# Specific Pathogen Free Ten Gene-Edited Pig Donor for Xenotransplantation

**DOI:** 10.1101/2025.05.06.652168

**Authors:** Kaixiang Xu, Heng Zhao, Baoyu Jia, Jiaoxiang Wang, Nazar Ali Mohammed Ali Siddig, Muhammad Ameen Jamal, Aqiang Mao, Kai Liu, Wenjie Cheng, Chang Yang, Taiyun Wei, Feiyan Zhu, Xiaoyin Huo, Deling Jiao, Jianxiong Guo, Hongfang Zhao, Wenmin Cheng, Yuemiao Zhang, Xiangyu Zhang, Lei Jiang, Zijie Zhang, Wei Zhang, Tingbo Liang, Hong-Ye Zhao, Bei-Cheng Sun, Hong-Jiang Wei

**Affiliations:** Yunnan Province Key Laboratory for Porcine Gene Editing and Xenotransplantation, Yunnan Agricultural University, Kunming, China; Yunnan Province Xenotransplantation Engineering Research Center, Yunnan Agricultural University, Kunming, 650201, China; The First Affiliation Hospital of Anhui Medical University, Anhui, 230022, China; The First Affiliated Hospital, Zhejiang University School of Medicine, Hangzhou, 311103, China; Faculty of Animal Science and Technology, Yunnan Agricultural University, Kunming, 650201, China; College of Veterinary Medicine, Yunnan Agricultural University, Kunming, 650201, China; Renal Division, Department of Medicine, Peking University First Hospital, Renal Pathology Center, Institute of Nephrology, Peking University, Key Laboratory of Renal Disease, Ministry of Health of China; Bio-X Center for Interdisciplinary Innovation, Yunnan University, Kunming, Yunnan 650091, China

**Author notes:** Authors contributed equally to this work (Kaixiang Xu, Heng Zhao, Baoyu Jia, Jiaoxiang Wang, Nazar Ali Mohammed Ali Siddig, Muhammad Ameen Jamal). Correspondence: Tingbo Liang, Hong-Ye Zhao; Beicheng Sun; Hong-Jiang Wei.

**Keywords:** Donor pig, gene editing, immune rejection, pathogenic microorganisms, xenotransplantation

## Abstract

Xenotransplantation has entered the clinical phase to fulfill the global organ shortage. However, recent clinical studies revealed that the xenograft from current gene-edited (GE) pigs still poses the risk of immune rejection, and biosafety concerns. In this study, we successfully constructed a large batch of 10- (GTKO/CMAHKO/ β4GalNT2KO/hCD46/hCD55/hCD59/hTBM/hEPCR/hCD39/hCD47) GE cloned (GEC) donor pigs by utilizing gene editing and somatic cell cloning technology, and successfully obtained F1 generation. Phenotypic characterization of 10-GEC pigs showed the deletion of three xenoantigens along with expression of seven human transgenes in various tissues. Digital droplet polymerase chain reaction, and whole genome sequencing indicated 2 copies of hCD46/hCD55/hCD59/hTBM/hCD39 and 1 copy of hEPCR/hCD47 in pig genome without significant off-target and damage to the porcine own functional genes. The validation results showed that 10-GEC pigs effectively inhibited the attacks of human antibodies, complement, and macrophages on porcine endothelial cells and alleviated the coagulation abnormalities between pigs and humans. 10-GEC pigs were negative for all zoonotic pathogens (48) including cytomegalovirus, except streptococcal infections. Kidney, heart, and liver xenografts from these 10-GE pigs were transplanted to non-human primates (NHP), which started working normally without hyperacute rejection. Among them, the heart and liver transplant recipient died without resuscitation due to unexpected interruption of oxygen supply, while the 2 kidney transplant recipients survived for 23 and 16 days, respectively. Pathological analysis showed that 10-GE pig kidney xenografts showed mesenchymal congestion, and fibrosis, cellular hyperplasia, with minor antibody and complement deposition, and significantly reduced the infiltration of CD68+ macrophage. In summary, we successfully produced a group of specific pathogen free GEC donor pigs that effectively mitigated immune rejection upon multi-organ transplantation to NHP.

## Introduction

Xenotransplantation is on the verge of entering the clinics to save the lives of patients with end-stage organ failure. In 2022, the University of Maryland Medical Center carried out the world’s first heart transplant from a 10-GE pig to a 57-year-old patient which sustained the life for up to 60 days without significant immune rejection ^[1]^. A year later, the University of California, USA performed the world’s second heart transplant from same 10-GE pig to a 58-year-old patient which survived for 40-days without significant immune rejection ^[2]^. Similarly, in 2024, the world’s first kidney xenograft from 11-GE pig was transplanted to a 62-year-old patient at Massachusetts General Hospital, which supported the life for 52 days without apparent immune-rejection ^[3]^. Subsequently, NYU Langone Medical Center performed the world’s second kidney transplant from a 1-GE pig to a 54-year-old patient which expired after 47 days. At the end of 2024, the same center performed the world’s third kidney transplant from a 10-GE pig to a 53-year-old patient which exhibited the longest survival of 130 days in live patients. More recently, Massachusetts General Hospital also performed the world’s fourth kidney xenotransplant from 11-GE pig-to-living-patient which is functioning well, hereby indicating the great potential of GE-pigs to address the global organs shortage.

Pig-to-human xenotransplantation still faces many obstacles like immune incompatibility including hyperacute rejection (HAR), antibody mediated rejection (AMR), cell-mediated rejection, coagulation disorders, and blood-mediated transient inflammatory responses, which has been effectively mitigated by gene editing ^[4]^. Nonetheless, the development of immune rejection involves the activation of multiple signaling pathways, and participation of multiple immune regulatory molecules, thus acquiring extensive gene editing in donor pigs. However, it is still unclear that how many genes should be edited, and which genetic combinations are suitable for which organ transplantation.

Knockout (KO) of the porcine GGTA1 gene (GTKO) was performed to eliminate HAR mediated by binding of the αGal xenoantigen to human natural antibodies which is necessary for long-term survival of porcine organs in humans ^[5–9]^. On this basis, it was further confirmed that KO of two other xenoantigens (CMAH, β4GalNT2) further reduces the risk of HAR ^[10–12]^. Thus 3-GE (GTKO/CMAHKO/β4GalNT2KO) donor pig is currently recognized as the basic configuration for xenotransplantation ^[13]^. Moreover, these 3-GE donor pigs have been initially attempted in human preclinical and clinical studies i.e., kidney xenotransplantation into brain-dead human recipients successfully alleviated immune-rejection, and thrombotic microangiopathy ^[14]^.

Human complement regulatory protein (CRP) can effectively inhibit most of the AMR leading to complement system activation as evidenced by in-vitro and in-vivo studies that expression of hCD46 or hCD55 significantly reduced cytotoxicity in porcine cells exposed to human serum ^[15, 16]^, and significantly prolonged the survival time of kidney and heart xenograft after transplantation to NHP ^[17–19]^. Similarly, the expression of hCRP along with GTKO reduced the early graft rejection ^[20]^), as the kidney xenograft from GTKO/hCD55 pigs survived up to 499 days into NHP ^[21]^. Nevertheless, the recent clinical trials of heart and kidney xenografts from 10-GE and 11-GE donor pigs still exhibited the complement deposition though both of CRP (hCD46/hCD55) were overexpressed ^[2, 3]^. These findings suggests that overexpression of 2 complement regulatory proteins is not sufficient to completely inhibit the cytotoxic effect of the complement system on porcine grafts, hereby necessitating the further genetic manipulations for complement system activation. Therefore, in this study, we proposed the simultaneous expression of three CRPs (hCD46, hCD55, and hCD59), on top of 3 xenoantigens (GTKO/CMAHKO/β4GalNT2KO) to further control the immune injury caused by complement system activation.

The development of thrombotic microangiopathy (TMA) due to coagulation system incompatibilities between pig and human often lead to the xenograft failure ^[22–24]^. Nevertheless, the expression of human coagulation regulatory proteins such as thrombomodulin (TBM) and vascular endothelial cell protein C receptor (EPCR) can act synergistically and increase the efficiency of protein C activation, which in turn inhibit graft microcirculatory thrombus formation, hereby increasing graft survival ^[25, 26]^. In addition, CD39 molecules hydrolyze the ATP and ADP which avoids excessive inflammatory response and tissue damage hereby inhibits the thrombus formation due to platelet aggregation ^[27]^. Therefore, we also proposed the simultaneous expression of those three coagulation regulatory proteins, (hTBM, hEPCR and hCD39) to reduce thrombotic microangiopathy.

Macrophage activation is also critical for xenograft survival as evidenced by our previous pig-to-NHP, and pig-to-brain-dead studies that a large number of CD68-positive macrophages infiltrated in pig kidney xenografts ^[28–30]^, which directly or indirectly led to loss of kidney xenograft function. Moreover, a large number of sedentary macrophages (Kupffer cells) are also present in the sinusoidal lining of porcine livers, and these cells continuously phagocytose human platelets, leading to a dramatic decrease in platelet count. Whereas in-vitro and in-vivo experiments revealed that CD47 expression can inhibit macrophage activation, inflammatory factor production and macrophage cytotoxicity ^[31, 32]^, and prolongs the survival time of kidney xenograft ^[33]^. Therefore, we also considered the expression human CD47 to prolong xenograft survival by inhibiting macrophage phagocytosis.

In recent years, zoonotic pathogenic microorganisms have been found to be critical for the survival of xenografts in both NHP, and human recipients ^[34, 35]^. For example, in the world’s first pig-to-living human patient cardiac xenograft, the reactivation and replication of porcine cytomegalovirus (PCMV/PRV) was a significant finding that might trigger a devastating inflammatory response leading to patient demise ^[36]^. Moreover, patients undergoing xenotransplantation underwent strong immunosuppressive treatment than allotransplantation, hereby increasing the likeliness of transmission of porcine pathogenic microorganisms to the recipient. The World Health Organization has clearly announced the serious biosafety concerns with 55 species of swine pathogenic microorganisms being potentially transmissible to humans ^[37]^. Therefore, the development of multi-GE pigs free from human-animal commensal pathogenic microorganisms could potentiate the clinical application of GE donor pigs for xenotransplantation.

Our team has conducted xenotransplantation research for almost a decade ^[28, 30, 38–41]^, and successfully inactivated the PERV to prevent cross-species viral transmission^[40]^, and produced 1-GE (GGTA1), 3-GE (GGTA1, B2M, CIITA)^[39]^, 4-GE (GGTA1, hCD55, hTBM, hCD39)^[30]^, and 8-GE (GGTA1, CMAH, β4GalNT2, hCD46, hCD55, hCD59, hTBM, hCD39)^[28]^ donor pigs.

In this study, we used gene editing, and somatic cell nuclear transfer (SCNT) technologies to construct 10-GEC (GTKO/CMAHKO/β4GalNT2KO/hCD46/hCD55/hCD59/hTBM/hEPCR/hCD39/hCD47) *Diannan* miniature donor pigs and monitored safety of these GE donor pigs for zoonotic pathogen transmission, and performed pig-to-nonhuman primate xenotransplantation to provide safe and effective donor pig types for the clinical application of xenotransplantation.

## 2. Materials and methods

All experimental animals and surgical procedures were approved by the Animal Ethics Committee of Yunnan Agricultural University.

### 2.1 Vector construction

sgRNAs were designed based on CRISPR/Cas9 technology to simultaneously target porcine GGTA1, CMAH, β4GalNT2 genes. Meanwhile, the PiggyBac system was utilized to overexpress human genes using human EF-1α promoter (hCD46, hCD55 and hCD59), the human endothelial cell-specific promoter ICAM2 promoter (hTBM and hCD39)^[28]^, and CAG promoter (hEPCR and hCD47) (Figure S1).

### 2.2 Cell transfection and screening and characterization

Cas9, sgRNA and human transgene overexpression vectors were transfected into fetal fibroblasts using electroporation, and antibiotic was added for screening at 48 h post-transfection, and surviving cells were cultured by limited dilution to obtain single-cell colonies. The genotypes of the obtained colonies were detected by polymerase chain reaction (PCR) and Sanger sequencing (Primers are shown in Table S1), and the cells with biallelic triple KO (TKO) of xenoantigens along with integration of human transgenes were selected as the donor for SCNT.

### 2.3 Genotyping of cell lines, cloned fetal pigs and piglets

The screened cell colonies were directly lysed, and the whole genomic DNA of fetal tissues and ear tissues of cloned piglets were extracted using high-salt method, and used as template for PCR, and the target gene fragments were amplified by PCR reaction. For cell colonies, we first confirmed the integration of five transgenes by gel electrophoresis, and then used the Sanger sequencing analysis to determine the genotypes of the target regions of three xenoantigens and finally the cells with the simultaneous KO of the three genes along with successful integration of the human transgenes were selected as donor cells for cloning. The 10-GE fetuses and GEC piglets were analyzed by PCR, and Sanger sequencing. The PCR reaction system (50 µL) was 400 ng of template, 2 µL of top and bottom primers, 1 µL of dNTP, 1 µL of Taq DNA polymerase (Cat #P505, Vazyme), 25 µL of buffer, waterwere made up.The PCR reaction program was 95°C for 5 min; 95°C for 30 s, 68~58°C (annealing temperature decreased by 1°C per cycle) for 30 s, 72°C for 1 min, 10 cycles; 95°C for 30 s, 58°C for 30 s, 72°C for 1 min, 25 cycles; and 72°C for 7 min. The primers are shown in Table S1.

### 2.4. Somatic cell nuclear transfer and embryo transfer

Oocyte collection, in vitro maturation, SCNT and embryo transfer were performed as described previously ^[42]^. Briefly, cultured cumulus-oocyte complexes were isolated from cumulus cells by treating with 0.1% (w/v) hyaluronidase. The oocytes with first polar body were selected and cultured in porcine zygote medium-3 (PZM-3) media containing 10% FBS, 0.0171g/mL sucrose, and 0.01μg/mL colchicine and incubated at 38°C, 5 % CO_2_ incubator for 0.5~1 h. The first polar body was enucleated via gentle aspiration using a beveled pipette in TLH-PVA, while the donor cells were injected into the perivitelline space of the enucleated oocytes by using micromanipulator (HMS-320D, Guangzhou Huayuehang Instrument Technology Co., Ltd.). The reconstructed embryos were fused with a single direct current pulse of 25V/mm for 20 µs using the Electro Cell Fusion Generator (HCF-230, Guangzhou Huayuehang Instrument Technology Co., Ltd.) in fusion medium. Embryos were then cultured in PZM-3 for 0.5–1 h and activated with a single pulse of 150 V/mm for 100 µs in activation medium. The embryos were equilibrated in PZM-3 supplemented with 5 µg/ml cytochalasin B for 2 h at 38.5°C in a four-compartment incubator (HincubatorDI-4R, Guangzhou Huayuehang Instrument Technology Co., Ltd.) and then cultured in PZM-3 medium with the same culture conditions described above until embryo transfer. The SCNT embryos were surgically transferred into the oviducts of the recipients, and a viable fetus was obtained through caesarian section after 33 days of pregnancy. A fetal fibroblast cell line was established, and subjected to PCR, and, Sanger sequencing for identification. Fetuses with biallelic TKO, and targeted overexpression of human transgenes were selected as donor cells for recloning. After completion of gestation period (~114 days), GEC piglets were obtained through natural delivery and identified by the same way of fetus identification.

### 2.5 Immunofluorescence staining

The paraffin-embedded tissue blocks were cut into 5 µm, transferred to glass slides, dewaxed using xylene and gradient alcohol, retrieved antigens with EDTA buffer (Cat#G1207, Servicebio Bio) at 92–98°C for 15 min, and cooled at room temperature. Then, sections were washed with PBS for three times (each time 3 min), incubated with autofluorescence quencher A (Cat#G1221, Servicebio Bio) at RT in dark for 15 min, washed with PBS for three times again, incubated with FBS at RT for 30 min, and dried. The dried sections were incubated with corresponding antibodies (Table S2). For visualization, corresponding secondary antibodies (Table S2) were diluted with PBS containing 10% FBS (v/v = 1:200) and used to incubate sections at 4°C in dark for 2 h, and a negative control was incubated with PBS containing 10% FBS. Then, sections were washed with PBS three times and stained with DAPI (Cat#G1012, Servicebio Bio) for 3 min. After washing with PBS for 1 min, autofluorescence quencher B (Cat#G1221, Servicebio Bio) was added for 5 min and washed three times again. Finally, sections were mounted with anti-fluorescence quencher (Cat#G1401, Servicebio Bio) and imaged using an OLYMPUS BX53 fluorescence microscope, and the fluorescence intensity of different samples was compared using ImageJ software.

### 2.6 H&E staining

Tissue samples were fixed in 4% paraformaldehyde for 48–72 h, processed by an automatic tissue processor (Yd-12p, Jinhua Yidi, medical appliance Co., Ltd, China) and embedded in a paraffin block (Yd-6D, Jinhua Yidi, medical appliance Co., Ltd, China). The paraffin blocks were cut into 5-um-thick sections using a Microm HM 325 microtome (Thermo Scientific, USA) and allowed to dry on glass slides overnight at 37°C. Thereafter, the tissue sections were deparaffinized in xylene and rehydrated through graded ethanol dilutions. Sections were stained with hematoxylin–eosin (H&E) (Cat. no. G1120, Solarbio) according to manufacturer’s instruction.

### 2.7 Flow cytometry

The cells were collected when confluence reaches 70% - 80%, digested with trypsin, and centrifuged at 1200rpm for 3 min. Then cells were resuspended with 50 μL PBS and incubated with primary antibodies (Supplementary table S2) according to 1×103 cells/μL at 37L in the dark for 15-30 min. After that, cells were washed with PBS and incubated with corresponding secondary antibodies at 37□ in the dark for 30 min. Cells were washed with PBS and detected by using CytoFLEX flow cytometer (Beckman Coulter, USA). Data were analyzed by FlowJo-V10.8.1 software.

### 2.8 Quantitative polymerase chain reaction (qPCR)

In order to evaluate the mRNA expression levels of seven transgenes, total RNA from heart, liver, lung, kidney and other tissues of WT and GEC pigs, as well as human umbilical cord tissues, was extracted using TRIzol reagent (Cat#ET111-01-V2, TransGen Biotech) according to the manufacturer’s instructions, and the concentration and RNA quality were detected. Complementary DNA (cDNA) was synthesized from total RNA using a PrimeScript RT reagent Kit (Cat#RR047B, TaKaRa) and was used as a template to perform qPCR in TB green-based qPCR instrument (CFX-96, Bio-Rad, USA). The reaction was performed in a 20 µL reaction mixtures comprising 10 µL of 2×TB Green® Premix Ex Taq™ (Tli RNaseH Plus) (Cat#RR420A, TaKaRa), 1 µL of cDNA, 1 µl of forward primer, 1 µL of reverse primer, and 7 µL of ddH2O (primers listed in Table S1). The reaction program is as follows: 95°C for 30 s, followed by 40 cycles of 95°C for 10 s, and 62°C for 45 s. Three technical replicates were conducted for each sample and the relative expression levels of target genes were quantified by 2^-ΔΔct.^

### 2.9 Droplet digital polymerase chain reaction (ddPCR) for transgenic copy numbers

The samples including the human blood (positive control), the pre-transfected porcine cells (negative control), the drug-selected cell lines, the fetal cells, and the heart, liver, lung, kidney, spleen and other tissues of the GEC pig were collected and extracted DNA. Then, these DNA samples were digested by MseI (Cat#FD0984, ThermoFisher), EagI (Cat#FD0334, ThermoFisher) and AsisI (Cat#FD2094, ThermoFisher) restriction endonuclease at 37°C for 2 h, and inactivated at 65°C and 85°C for 20 min. The digested product was diluted to 5ng/µl and used as a ddPCR template. The primers and probes corresponding to hCD46, hCD55, hCD59, hTBM, hCD39 and GAPDH genes (Table S1), ddPCR super mix (Cat#1863024, Bio-Rad), and DNA template were mixed to prepare a 25uL reaction system. Then, ddPCR droplets were generated with QX100 Droplet Generator (Bio-Rad) and transferred to a 96 well plate. PCR reactions were performed at 94°C for 5 min, 94°C for 30 s, 56°C for 1 min (40 cycle), and 98°C for 10 min. After the reaction, the 96 well plate was placed in Bio-Rad Droplet Reader and QunataSoft Software was used to set up the experimental design and read the experiment. Once the program was finished, the copy numbers were analyzed through gating, according to the manufacturer’s instructions (Bio-Rad).

### 2.10 Porcine mass screening for pathogenic microorganisms

Whole blood was collected from pigs through the anterior vena cava and serum was separated, and nasal, oral, and pharyngeal swabs were collected from GEC pigs, and DNA, RNA extractions, CDNA synthesis and PCR and qPCR were performed as reported elsewhere in the methods. The samples were processed for direct smear method ^[43]^, saturated saline floating method ^[44]^and centrifugal precipitation method ^[45]^ accordingly. Meanwhile, the samples collected above were sent to Suzhou Xishan Biotechnology Co., Ltd. in China (a third party) for pathogenic microorganisms’ detection, and the macro-genome sequencing analysis was carried out at Hangzhou Jieyi Biotechnology Co., Ltd. and Beijing Novozymes Technology Co.

### 2.11 Antigen-antibody binding assay

Whole blood (10 mL each) of healthy volunteers was collected and serum was separated. Meanwhile, 10 mL of whole blood from wild-type (WT), and 10-GEC donor pigs were collected to isolate peripheral blood mononuclear cells (PBMCs). Based on the antigen-antibody binding assay, inactivated serum from human was incubated with WT and 10-GEC donor pig PBMCs, and then treated with anti-IgG antibody (Cat#628411, Invitrogen) and anti-IgM antibody (Cat#A18842, Invitrogen), respectively), and the levels of binding of IgG and IgM to pig cells was detected by cell flow cytometry.

### 2.12 Complement-dependent cytotoxicity assay

The inactivated human serum was diluted 1:1 in staining buffer (PBS containing 1% FBS). And PBMCs derived from WT and 10-GEC pig were collected, washed twice and resuspended in staining buffer. 1×10^5^ cells were incubated with 50 µl inactive rhesus monkey serum (test group) or 50 µl PBS (negative control) for 30 min at RT. Then cells were washed with cold staining buffer to terminate reaction and incubated with rabbit complement sera (1:3 dilution, Cat. no. S7764, Sigma) for 30 min at RT. Finally, cells were stained with PI staining solution (Cat. no. GA1174, Servicebio Bio) for 2 min, and analyzed for cell death using a CytoFLEX flow cytometer (Beckman Coulter, USA).

### 2.13 Coagulation assay

Endothelial cells (1×10^6^ cells/well) were inoculated in 12-well plates and incubated at 37°C for 24 h. Cells were washed with PBS and then in assay buffer (50 mM Tris HCl, 150 mM NaCl, 25 mM CaCl□, 0.1% BSA, pH 7.5, 37°C). Then, added 100 μL of 20 U/mL thrombin (Ebixon - CAS# 9002-04-4) or human thrombin to each well and incubated for 30 min at 37°C. Then discarded the supernatant and transfer to a 96-well flat-bottomed plate, added 10 μL of 1 mM D-Lys(Z)-Pro-Arg-pNA (MCE-HY-P002A-1), and analyzed the supernatant in the zymometer at at absorbance of 405 nm at 37°C, and the concentration of unconjugated thrombin in the supernatant was calculated from the standard curve of thrombin.

### 2.14 Macrophage phagocytosis assay

Differentiation of the human macrophage cell line THP-1 was ensured by observing the differentiation status of the cells by incubating them for 3 days with 200 ng/mL of PMA (16561-29-8) in THP-1-specific medium. 8-TG PAECs, 10-TG PAECs, WT PAECs, and Human umbilical vein endothelial cells (HUVECs) (as the target cells) were stained with the fluorescent stain CFSE (ab113853), and then targeted cells were incubated with human differentiated THP-1 cells (as effector cells) at 1:1 ratio at 37°C for 6 h. Macrophages stained with anti-human CD11b antibody (abs180040-25T) phagocytosed the CFSE-labeled target cells, which were then measured using FACS, and phagocytic activity was calculated as phagocytes exerting phagocytosis/total number of phagocytes.

### 2.15 Kidney, heart and liver xenotransplantation from pigs to non-human primates

Pig-to-NHP xenotransplantation was carried out at the Yunnan Provincial Xenotransplantation Engineering Research Center of Yunnan Agricultural University, where the surgical team from the First Affiliated Hospital of Anhui Medical University, and First Affiliated Hospital of Zhejiang University School of Medicine performed the kidney, heart, and liver xenograft transplants to Tibetan macaque. Urine and blood biochemistry, immune index, coagulation function, infection, immunosuppressant concentration and other tests were conducted regularly after the operation, and the blood flow of transplanted kidneys was detected through ultrasound, and the clinical medical personnel and veterinary professionals took turns to take care of the patients, and according to the various test indexes as well as the mental status of the recipient monkeys, and treated accordingly.

### 2.16 Statistical analysis

The data were expressed as the mean ± standard deviation of at least three independent repeated experiments, and the comparison between the two groups of data was performed by student t-test for significance (*P<0.05, **P<0.01). Figure formation and data analysis were performed by GraphPad prism.

## 3 Results

### 3.1 Generation of 10-GEC pigs

To produce GGTA1/CMAH/β4GalNT2/hCD46/hCD55/hCD59/hTBM/hCD39 hEPCR/hCD47 10-GEC donor pigs, we utilized the CRISPR/Cas9 system, PiggyBac transcriptase system combined with somatic cell cloning technology, and first co-transfected the previously constructed 3KO (GGTA1/CMAH/β4GalNT2) vectors, with 5 transgenes (hCD46/hCD55/hCD59/hTBM/hCD39) vectors, and PB transcriptase vectors into the fetal fibroblasts of male *Diannan* miniature pigs. Fibroblasts were screened for Puro resistance and genotyped to obtain 8-GE cell colonies which were then used as donor for the first round of SCNT and successfully obtained 8-GEC fetuses, and after the fetuses were identified, the 8-GEC fetuses with good traits and phenotypes were selected for recloning to obtain 8-GEC pigs. Keeping in view the 8-GEC pigs with high survival rate, the corresponding fetal cell lines were selected for the second editing. The PB transcriptase vector of 2 transgenes (hEPCR/hCD47) was transfected into 8-GE fibroblasts, which were screened for HygR resistance and cultured in limited dilution to obtain 10-GE single cell colonies, which were then used as a donor cell for SCNT, and 10-GEC fetuses were obtained and genotyped, used as a donor for recloning to obtain 10-GEC pigs (Figure S1A). For simultaneous KO of GGTA1, CMAH and β4GalNT2 genes, 2, 2 and 3 sgRNAs at exon 3, exon 4 and exon 2 of corresponding genes were designed, (Figure S1B). The expression of 5 transgenes was obtained using the human EF-1α promoter (hCD46, hCD55 and hCD59), the human endothelial cell-specific promoter ICAM2 promoter (hTBM and hCD39). While, the CAG promoter was used to drive the expression of hEPCR and hCD47 genes (Figure S1C).

After electroporation and drug selection, a total of 49 cell colonies were obtained. At first, these colonies were identified by PCR, out of which 34 carried the 5 transgenes (Figure 1A). Then 32 colonies were randomly selected for Sanger sequencing and results showed that 10 colonies (C2#, C3#, C5#, C6#, C10#, C21#, C24#, C26#, C34#, and C44#) for GGTA1, 6 colonies for β4GalNT2 (C2#, C3#, C10#, C12#, C21#, and C24#), and 4 colonies (C2#, C3#, C10#, and C24#) for CMAH were biallelic KO. Among these 20 colonies, 4 (C2#, C3#, C10#, and C24#) were biallelic TKO carrying human transgenes (Table S3-4), thus colony C3# was used as a donor for SCNT and eighteen live 33-day-old cloned fetuses were obtained (Figure 1B). Among them, 13 live fetuses (C3F01 to C3F13) successfully carried the 5 transgenes (hCD46/hCD55/hCD59/hTBM/hCD39, Figure 1C), and were biallelic TKO (Figure S2, Supplementary Table S5). qPCR results showed strong mRNAs expression of hCD46, hCD55 and hCD59 transgenes, and weak expression of hTBM and hCD39 transgenes in 8-GEC fetal fibroblasts (Figure 1D), which might be due to the specific expression of endothelial cell promoter. While the WB results showed that hCD46, hCD55, hCD59, hTBM and hCD39 proteins were expressed in (C3F01, C3F03 and C3F07) 8-GEC fetal tissues (Figure 1E). Therefore, these three 8-GEC fetal fibroblasts were used for re-cloning and reconstructed embryos were transferred into eight surrogate sows, giving birth to 38 piglets, of which 18 survived (Supplementary Table S6), and their genotype for GGTA1, CMAH and β4GalNT2 was consistent with 8-GEC fetal fibroblasts (Supplementary Table S7). Additionally, we also cloned a batch of piglets (n=13) directly through the C3# cell colony (Figure 1F and Supplementary Table S8), and these individuals grew healthy with competent reproductivity. Their genomes successfully carried 5 transgenes (Figure 1G), and some of them also were biallelic TKO (Figure 1H and Supplementary Table S9).

**Figure 1.**
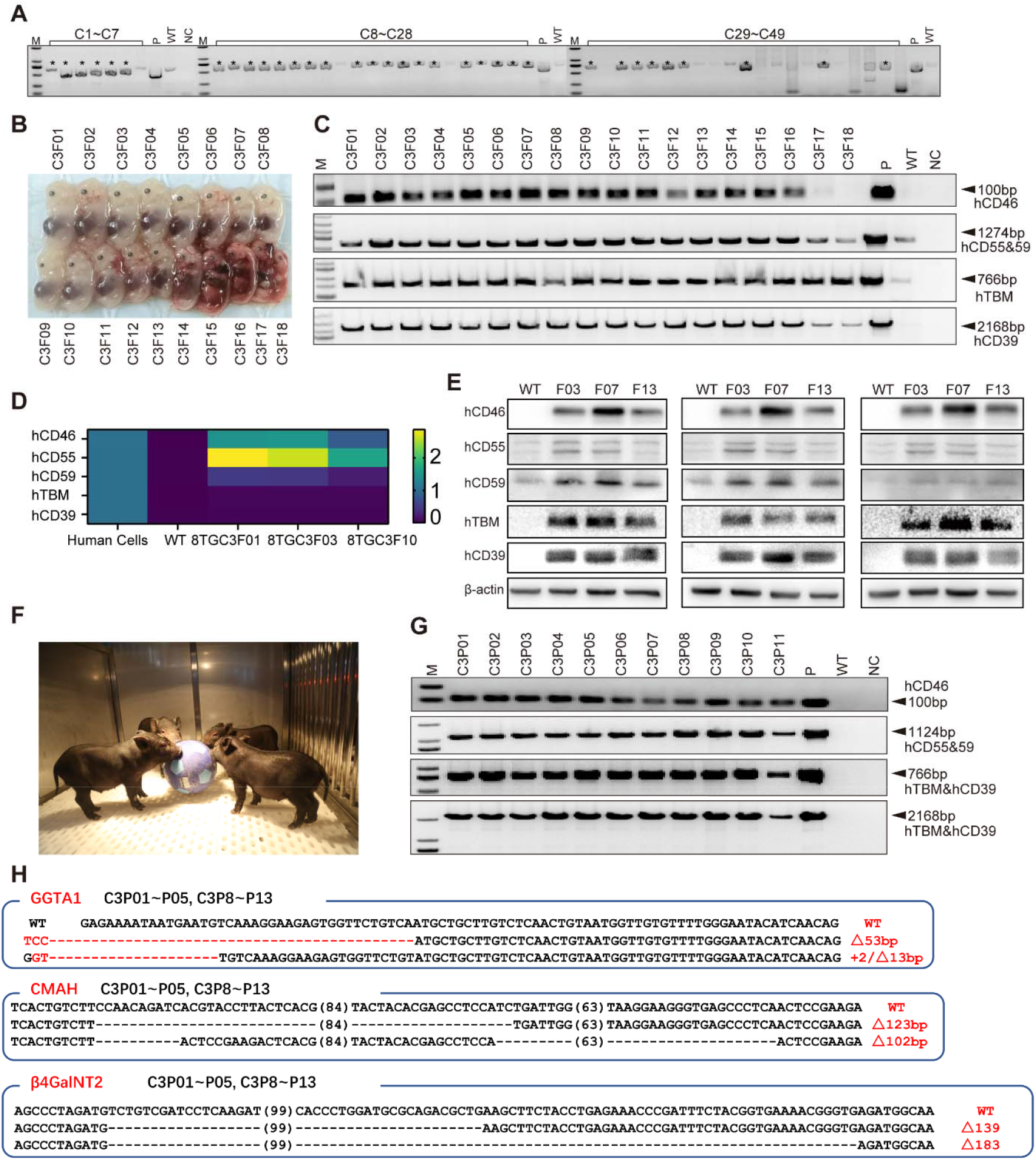
Construction of gene editing vectors for KO of xenoantigens, with 5 genes overexpression of human transgenes and screening colonies. (A). PCR identification of cell colonies for successful integration of 5 human transgenes (hCD46, hCD55, hCD59, hTBM and hCD39) into pig genome. (B) Photo of GEC fetuses obtained after 33-day pregnancy. (C-E) Identification of GEC fetuses for successful integration of 5 human transgenes by PCR (C), qPCR (D) and western blot (E). (F) Photo of 8-GEC pigs obtained after completion of pregnancy. (G) PCR identification of 8-GEC pigs for successful integration of 5 human transgenes. (H) The sequence of the targeting region of GGTA1, CMAH, and β4GalNT2 genes in 8-GEC pigs by Sanger sequencing.

We next performed the copy number analysis of 5 human transgenes on heart, liver, spleen, lung, and kidney tissues of 8-GEC pigs, and the results showed that these tissues possess 3 copies of each transgene (Figure 2A). The qPCR analysis showed the mRNA expression of all transgenes on heart, kidney and liver tissues, of 8-GEC pigs (Figure 2B). Moreover, the immunofluorescence staining showed that except αGal antigen, the other 2 xenoantigens (Neu5Gc and Sda) were deficit and 5 human transgenes were expressed on the kidneys of 8-GEC pigs (Figure 2C). We carefully analyzed the genotype of the GGTA1 gene in the 8-GEC pigs and found that, despite the pure mutation of the GGTA1 gene in exons, the allele with a deletion of 53 bp, which was calculated from the start codon ATG, was exactly 30 bp missing, presenting an integer multiplication of 3 (Figure 2D), and did not result in a shifted-code mutation of the GGTA1 gene, but rather made the GGTA1 gene to form a truncated protein, probably because this truncated protein can still catalyze the synthesis of αGal antigen, which ultimately led to the success of incomplete knockdown of GGTA1. Therefore, in the process of generating 10 GEC pigs, we redesigned the targeting vector at exon 8 of the GGTA1 gene (Figure S1B).

**Figure 2.**
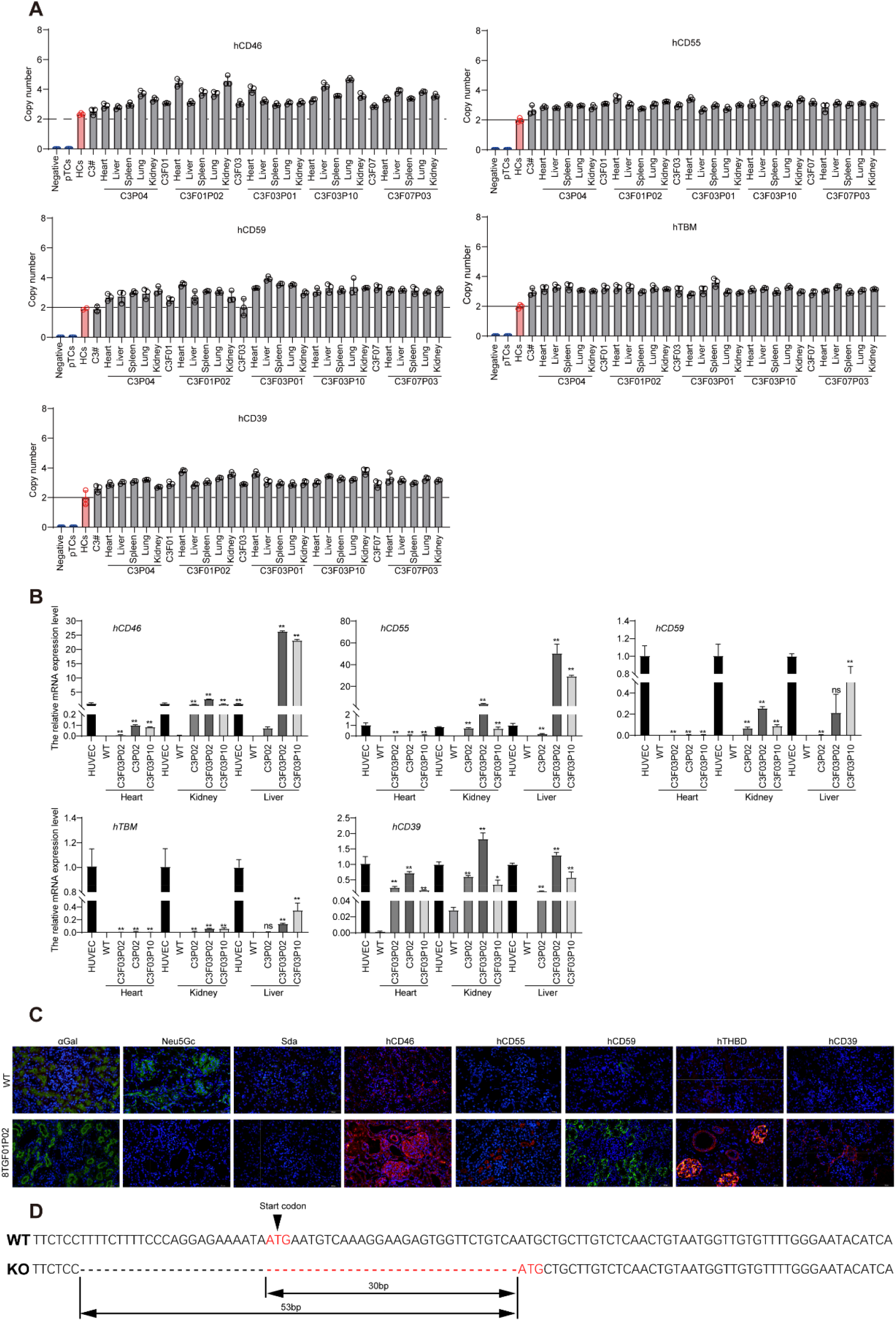
Copy number and Phenotype of 8-GEC pigs. (A) Copy numbers of transgenes hCD46, hCD55, hCD59, hTBM and hCD39 in donor cell lines, cloned fetus, and piglets. (B) The mRNA expression levels of hCD46, hCD55, hCD59, hTBM, and hCD39 genes in heart, kidney, liver, and lung tissues of GEC pigs; HUVEC, Human umbilical vein endothelial cell. (C) The expression of xenoantigens with 5 transgenes in GEC pig kidney was confirmed by immunofluorescence. (D) Sanger sequencing of the targeting region of GGTA1 in 8-GEC pigs.

On the basis of 8 GEC fetus (C3F01), we performed a second transfection and obtained 58 cell colonies after HygR resistance screening and limited dilution culture. First, PCR identification of the target region of exon 8 of the GGTA1 gene showed that 16 colonies (C1#, C3#, C4#, C6#, C7#, C12#, C15#, C16#, C19#, C22#, C23#, C28#, C31#, C36#, C54#, C58#) had obvious split bands in the target region (Figure S3A), which were predicted to be cloning sites with large fragment deletions on the GGTA1 gene. Sanger sequencing showed that all these cloning sites contained biallelic mutation of more than 100 bp (Supplementary Table S10). Then, these 16 cloning sites were identified by PCR, and the results showed that all 16 cloning sites had hEPCR and hCD47 gene insertions (Figure S3B). Then, we selected one colony (C3#) with a large deletion mutation in exon 8 of the GGTA1 gene as a donor for SCNT, and five 33-day-old viable fetuses were obtained (Figure S3C). Genotyping results (Supplementary Table S11), showed that all of the 10-GEC fetuses were genotypically consistent with the donor cell line i.e., biallelic TKO, along with integration of 7 human transgenes (Figure S3D, E). Moreover, the mRNA of 7 human transgenes was expressed in the 10-GEC fetuses (F01-F03, Figure S3F), hence it was used as a donor fetal fibroblast for recloning and reconstructed embryos were transferred into 118 surrogate sows, and delivered a total of 582 GEC piglets with an average of 4.9 cloned piglets per surrogate sow (Supplementary Table S12, Figure 3A). Genotyping results showed that 10-GEC pigs were genotypically consistent with the donor fetal fibroblast. Similarly, we characterized some of the cloned pigs by PCR and Sanger sequencing, which showed that all of the cloned pigs were biallelic TKO (Supplementary Table S13, Figure 3B). Genome-wide off-target analysis showed that only 1 intergenic region (RRAGD and ANKRD6 genes) was off-target in the 10-GEC pigs in the presence of 4 base mismatches (Table S14). Transgene copy number analysis revealed that the copy number of the hCD46, hCD55, hCD59, hTBM, and hCD39 transgenes changed from 3 in 8-GEC pigs to 2 copies in 10-GEC pigs, while the copy number of the hEPCR and hCD47 genes was 1 in 10-GEC pigs (Figure 3C). Genome-wide insertion site analysis showed that the three transgene copies were inserted into the intergenic regions or introns of chromosomes 5, 6, and 13, respectively, and did not cause damage to the pig’s own functional genes (Table S15). Immunofluorescence staining of kidney tissues of 10-GEC pigs showed that 3 xenoantigens were deficit, and 7 human transgenes were expressed, with significantly higher expression of hEPCR and hCD47 (Figure 3D). The qPCR analysis showed the mRNA expression of all transgenes on heart, kidney and liver tissues of 10-GEC pigs (Figure S4A). In order to obtain F1 generation, 10-GEC male pigs (n=4) were crossed with 8-GEC sows (n=4), giving birth to 24 F1 generation piglets (Figure S4B, Table S16), which carried TKO biallelic mutation with integration of 7 human transgenes (Table S17-18), hereby demonstrated that our 10-GEC pigs can reproduce offspring, which lays the foundation for the expansion and breeding of xenotransplantation donor pig populations.

**Figure 3.**
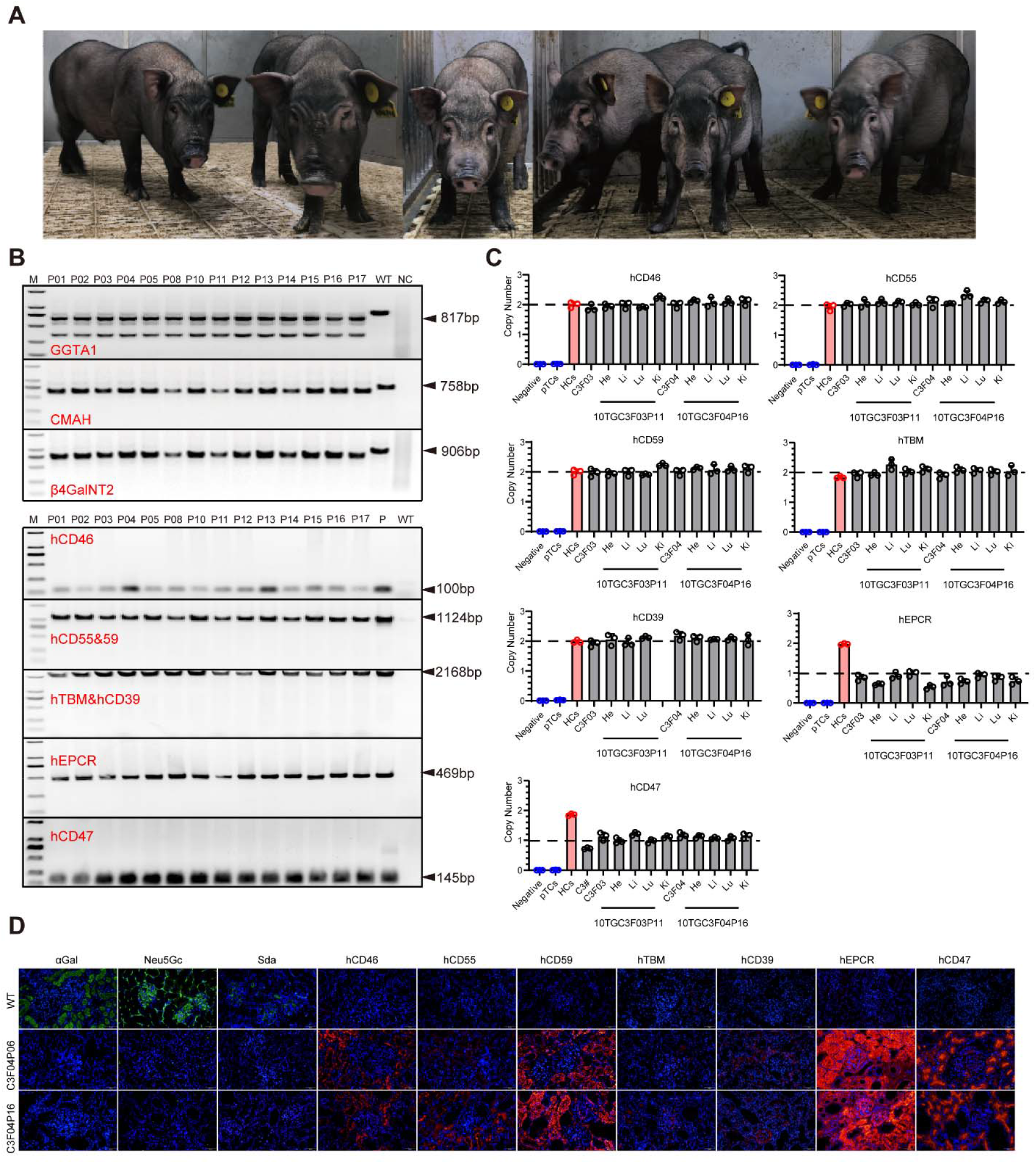
Identification of 10-GEC pigs. (A) Photo of 10-GEC pigs (B) PCR identification of 10-GEC pigs for TKO with successful integration of 7 transgenes. (C) Identification of copy number in 10-GEC pigs by ddPCR. (D) The expression of 3 xenoantigens with 7 transgenes in 10-GEC pig kidney was confirmed by immunofluorescence.

Next, we collected nose/mouth/throat swabs, feces, dander, skin, serum, and whole blood samples from 6-to 9-month-old 10-GEC pigs housed in SPF-grade facilities, and performed large-scale screening for pathogenic microorganisms and results showed that our 10-GEC pigs were negative for a total of 47 pathogenic microorganisms, including porcine cytomegalovirus, except streptococcal infection (Table S19).

### 3.2 Functional validation of ten gene edited xenograft donor pigs

Flowcytometric analysis showed the expression of αGal, Nue5Gc and Sda antigens were absent, and 7 human transgenes (hCD46, hCD55, hCD59, hTBM, hCD39, hEPCR and hCD47) were expressed in 10-GEC pigs as compared to WT (Figure 4A). The antigen-antibody binding assay revealed that binding ability of human IgG and IgM to 10-GEC pig’s aortic endothelial cells (PAECs) was significantly reduced than the WT control (Figure 4B). And the complement-mediated assay showed that the survival rate of 10-GEC pig’s PBMCs was higher than that of WT control (Figure 4C). Coagulation assay showed that 10-GEC pig’s PAECs were significantly more bound to thrombin than WT control (Figure 4D). To confirm that hCD47 effectively avoids phagocytosis of porcine cells by human macrophages, porcine cells were incubated by mixing with PMA-induced human THP-1 cells, and the results showed that 10 GEC pig’s PAECs were phagocytosed significantly less by human macrophages (Figure 4E). In conclusion, 10 GEC pigs can effectively resist immune rejection caused by antigen-antibody binding, complement system activation, coagulation dysregulation and macrophage activation in xenotransplantation.

**Figure 4.**
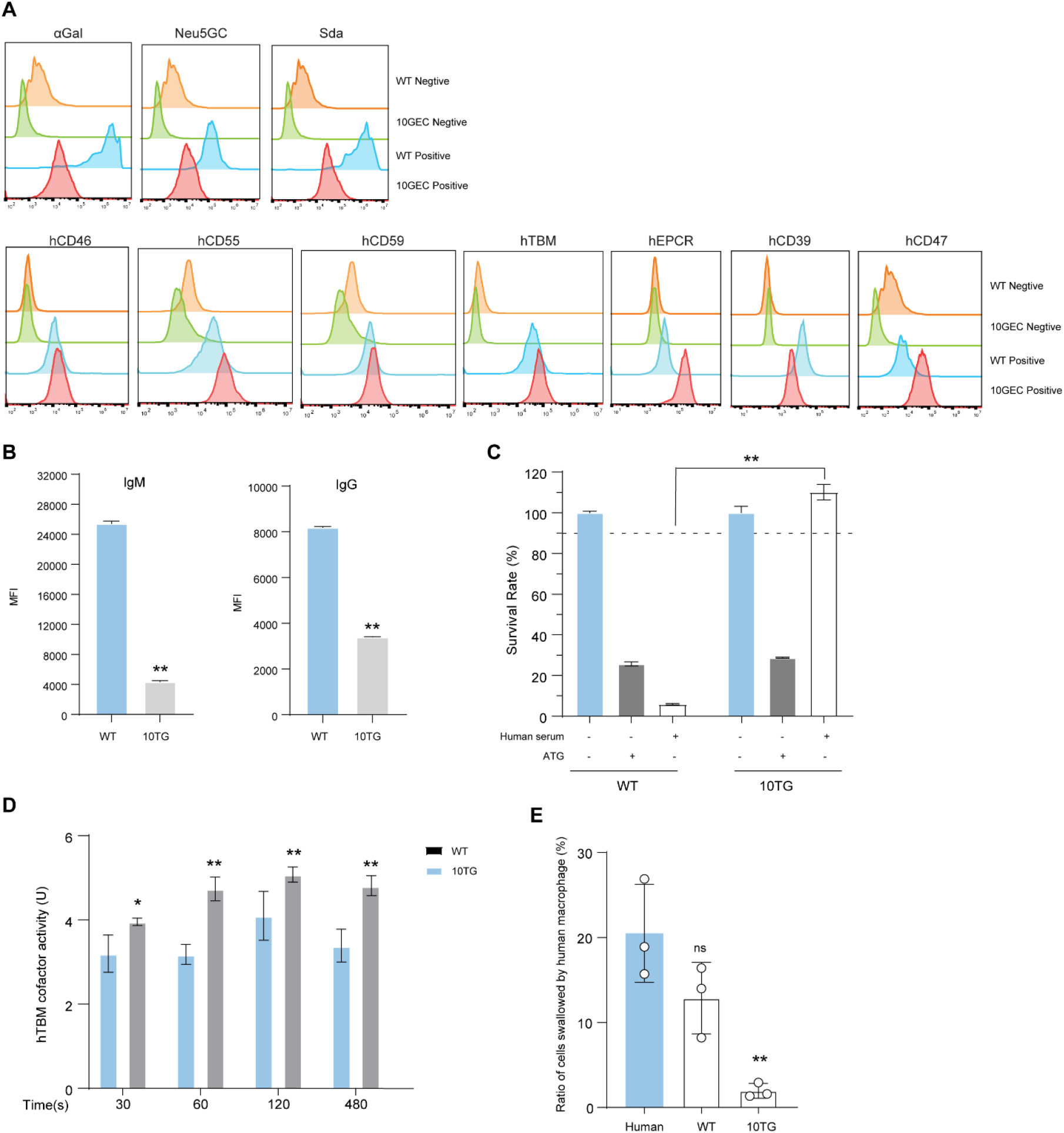
Identification and functional verification of 10-GEC pigs. (A) The expression of 3 xenantigens and 7 human transgenes in 10-GEC pig aortic endothelial cells (PAECs) was confirmed by flow cytometry. The levels of human IgG and IgM binding to 10-GE porcine PAECs. (C). The survival rate of 10-GEC porcine PBMCs. (D) Coagulation activity of 10-GEC pigs’ PAECs. (E) Phagocytosis of 10-GEC PAECs by human macrophages.

### 3.3 Organ transplantation from 10-gene edited xenotransplantation donor pigs to nonhuman primates

In order to evaluate the effectiveness of 10-GEC pigs as donor for xenotransplantation pigs, we performed multiple organ xenotransplantation i.e., kidney, heart, and liver from 10-GEC pigs to Tibetan macaques. First, we collected serum from 20 Tibetan macaques and isolated PBMCs from 10-GEC pigs (10TGC3F02P02, 10TGC3F02P12), and performed IgM and IgG antibody binding assays as well as complement-dependent cytotoxicity assays, from which 3 Tibetan macaques, and 1 rhesus monkey, were selected to carry out xenotransplantation (Table S 20). Two days before surgery, immunosuppression was induced with anti-CD20 mAb and ATG, respectively, On the day of surgery, we simultaneously obtained 1 kidney, 1 heart, and 1 lung from one 10-GEC pig (10TGC3F02P02), and transplanted to 3 Tibetan macaques after in vitro perfusion, and 1 kidney from another 10-GEC pig (10TGC3F04P12) was transplanted to 1 Tibetan macaque after in vitro perfusion. Soon after completion of surgery, the xenograft started functioning normally i.e., kidney produced urine, heart began to beat, and the liver secreted bile, without apparent HAR (Figure S5 and Figure S6A, B). The heart and liver transplant recipients were accessed to an anesthesia ventilator on the operating table for clinical observation and treatment after surgery, and at 3 days postoperatively, unexpected interruption of the oxygen supply led to cardiac arrest in the heart transplant recipients, which resulted in unsuccessful resuscitation, and death in the liver transplant recipients at day 4 post operation. We performed HE staining of the liver tissues, and the results showed that on postoperative day 1, there was partial vascular endothelial breakdown (Figure S6C), but no complement C4d deposition was detected (Figure S6D); and on postoperative day 4, there was necrotic disintegration of stem cells (Figure S6E, F). Because of the unexpected deaths of heart and liver transplant recipients, we summarized and analyzed mainly the 2 kidney transplant recipients in subsequent experiment.

During postoperative care, medications were administered regularly according to the developed immunosuppression protocol, and appropriate treatment or nutritional supplements were given accordingly ^[28]^. The 2 Tibetan macaques (M1 and M2) with transplanted kidneys survived for 23 and 16 days, respectively. In both recipient monkeys, the renal function indices of serum creatinine, urea and cystatin C remained relatively stable up to 10 days after surgery, and then gradually increased (Figure 5A-D). The red blood cell count remained stable at all times (Figure 5E), and the platelet count dropped to its lowest point on the day of transplantation and was maintained at about 1×10^11^/L thereafter (Figure 5F). The dissection of the xenografted kidney showed soft texture and large number of hemorrhagic spots (Figure 5G). The pathological analysis showed that M1 recipient transplanted kidney manifested interstitial hemorrhage, and inflammation, tubular infarction, and vacuolar degeneration, while the M2 transplanted kidney showed erythrocyte titularity, neutrophil infiltration, tubular calcium salt deposition, and small arteriolar thrombosis (Figure 5H). Masson staining showed small arteriolar-thrombus, interstitial fibrosis, cell proliferation, and tethered membrane hyperplasia in M1 transplanted kidney; lesions such as thrombus in the entry glomerular small artery, interstitial hemorrhage, glomerular cell proliferation, and glomerular platinum ear in M2 kidney recipient (Figure 5I).

**Figure 5.**
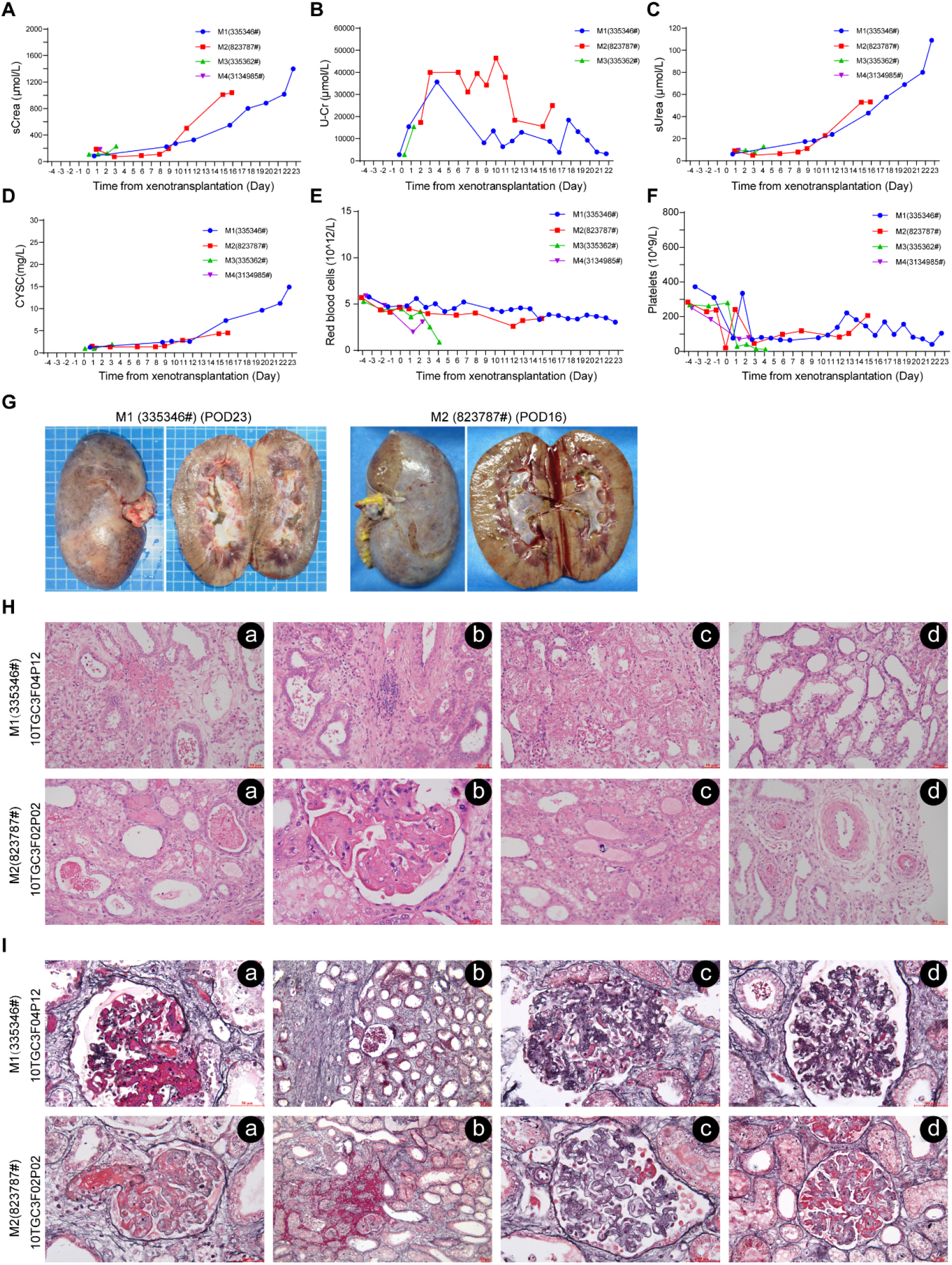
The renal function, erythrocytes indexes of recipient monkey and pathological sections of kidney xenograft from 10-GEC pigs. (A-D). The levels of (A)creatinine in serum, and (B) urine. Levels of (C) urea, and (D)cystatin C in the serum of recipient monkey. (E-F) The number of red blood cells, and platelets (F). (G-I) Pathological section of kidney xenograft. (G) Gross sectioning of kidney xenograft of day 23 and 16 post-transplantation. (H) H&E staining of two kidney xenografts. (I) Manson staining of two kidney xenografts from 10-GEC pigs

In terms of humoral immune rejection, the serum antibodies IgA, IgM and IgG of the 2 recipient monkeys were maintained at relatively stable levels (Figure 6A), and complement C3 and C4 were maintained at relatively stable levels in M1 recipient, whereas increased significantly in M2 recipient after 10 days of transplantation (Figure 6B). We found fluctuation in the leukocyte and neutrophil counts in both recipients. Monocyte counts in both recipients increased significantly at the late stages of transplantation, lymphocytes and basophils remained relatively stable, eosinophils remained relatively stable in M2 recipient, and basophils fluctuated greatly in the late stage of transplantation in M1 recipient (Figure 6C). Immunofluorescence staining showed IgM and IgG antibody deposition, without complement C4d and C5b-C9 deposition in both recipients, except C3c deposition in the M2 recipient (Figure 6D). In terms of cellular immunity, the transplanted kidneys of the both recipient monkeys showed a large number of CD68+ macrophages as well as sporadic CD3+CD8+ T-lymphocyte infiltration, but no CD57+ NK cells as well as CD3+CD4+ T-cell infiltration (Figure 6E).

**Figure 6.**
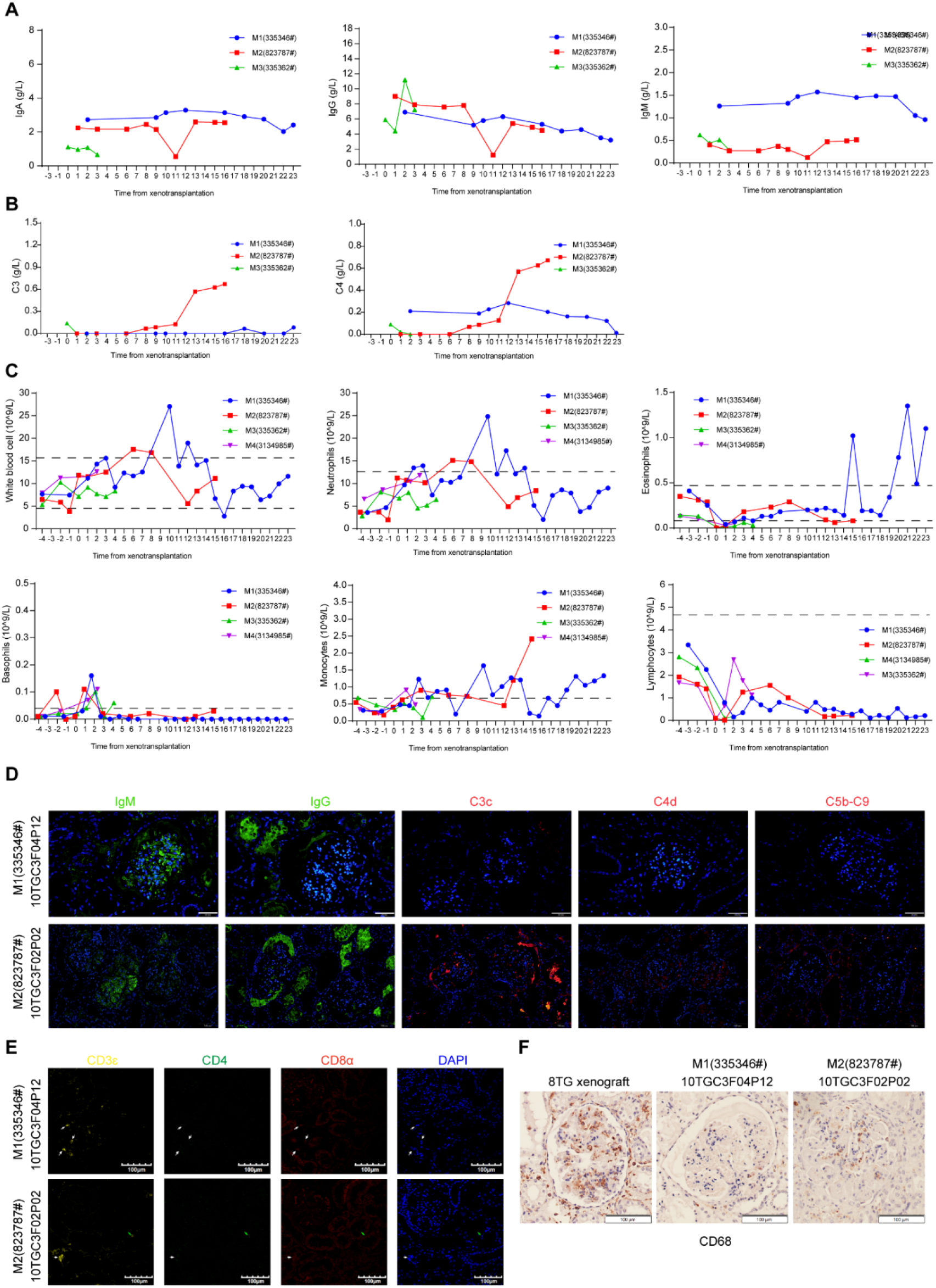
Humoral, cellular immunity recipient monkey, and pathological sectioning of kidney xenograft. (A). The levels of IgA, IgG, and IgM in serum of recipient monkey. (B). The levels of complement C3 and C4 in serum of recipient monkey. (C). The numbers of white blood cells (WBC), neutrophils, monocytes, lymphocytes, eosinophils and basophils in the whole blood of recipient monkey. (D). Immunoglobulins and complement deposition in 10-GE porcine kidney xenograft confirmed by immunofluorescence. (E). T cell infiltration in porcine kidney xenograft confirmed by immunofluorescence (scale bar=100 μm). (F). CD68+ macrophages infiltration in porcine kidney xenograft confirmed by immunohistochemistry, scale bar=100 μm.

Further, most of the liver function indexes, including TP, ALB, GLB, TBIL, TBA, AST, ALT, and ADA, remained relatively stable, and ALP showed a trend of gradual increase (Figure S7 A-I). In terms of electrolyte balance, electrolytes Na, K, Cl, Ca, Mg, HCO3, were maintained at a stable level, but blood P increased significantly in the late stage of transplantation (Figure S7 J-P).

## 4 Discussion and conclusion

Xenotransplantation is about to enter the clinic so it is urgent to construct and select the donor pigs with suitable gene combination. In recent years, with the rapid development of gene editing technology, GE donor pigs with various gene combinations have been emerging, and 10-GE donor pigs seems to be the standard for clinical xenotransplantation. However, editing more genes is comparatively difficult and time taking. Meanwhile, cross-species transmission of pathogenic microorganisms poses a great challenge to the clinical application of xenotransplantation. Xenotransplantation is allowed to be used for clinical phase only if multiple GE donor pigs are safe and have stably inheritance of the genes. In this study, we successfully constructed 10-GE donor pigs using the CRISPR/Cas9 system, PiggyBac transposase system and somatic cell cloning. These donor pigs are free from many pathogenic microorganisms such as porcine cytomegalovirus, and exhibit normal reproductive ability which makes us a step forward in the entry of clinical xenotransplantation.

Two companies, Revivicor and eGenesis, have been working on the development of xenotransplantation donor pigs for a long time. Currently, both companies have obtained more than 10-GE xenotransplantation donor pigs, and have carried out several cases of pig to non-human primate ^[46–48]^, pig to brain dead human ^[48, 49]^, and pig-to-living-patient ^[2, 3, 35]^, thus have greatly advanced the clinical use of xenotransplantation. The 10-GE donor pig developed by Revivicor knocked out 3 xenoantigens, 1 growth hormone receptor, 2 complement regulatory proteins (hCD46 and hCD55), 2 coagulation regulatory proteins (hTBM and hEPCR), 1 macrophage phagocytosis inhibiting gene (hCD47) and 1 anti-inflammation gene (hHO-1). The 11-GE donor pig developed by eGenesis also KO 3 xenoantigens, knocked in 2 complement regulatory proteins (hCD46 and hCD55), 2 coagulation regulatory proteins (hTBM and hEPCR), 1 macrophage phagocytosis-inhibiting gene (hCD47), and 2 anti-inflammatory genes (hHO-1 and hA20), and inactivated all endogenous retro viral viruses ^[46]^. In present study, we developed 10 GE donor pigs with KO of three xenoantigens, and overexpressed three complement regulatory proteins (hCD46, hCD55, and hCD59), three coagulation regulatory proteins (hTBM, hEPCR, and hCD39), and one macrophage phagocytosis suppressor gene (hCD47). Compared with the previous two companies, our donor pigs enhanced complement system activation inhibition and anticoagulant effects. Among them, hCD59 inhibits the formation of membrane-attacking complexes, which may further inhibit graft injury by human complement ^[50]^. hCD39 has an inhibitory effect on inflammation along with anticoagulation ^[51]^. In addition, the pigs we used were a locally endemic small pig strain with adult weights between 50-80 kg, which is similar to the average weight of humans, and the heart and kidney organ sizes were closer to those of humans as shown by a large number of slaughtering experiments and measurements. These characteristics avoided metabolic regulation disorders in pigs caused by knockdown of the GHR gene.

While the development cycle of donor pigs can be shortened by transfecting multiple positive cells for multiple gene editing with multiple mass screening particles in a single transfection, and it is also necessary to screen and characterize more single-cell colonies before obtaining compliant cell lines for somatic cell cloning. During the first transfection and screening process, we obtained a total of 49 single-cell cloning sites, and eventually only 4 colonies (C2#, C3#, C10#, and C24#) were 8-GE. Moreover, these four colonies contained all wild-type genotypes, and among them, three cloning sites, C2#, C10# and C24#, might have single allele knockout of the CMAH gene and the knockout fragment was an integer multiple of 3. Therefore, we finally chose the C3 cloning site for cloning. Among the 13 fetal pigs obtained by cloning, the genotypes of GGTA1, CMAH and β4GalNT2 genes were identical, and all of them were double allelic KO. However, a 53-bp deletion in one of the alleles of the GGTA1 gene in fetal pigs and piglets produced by cloning from cloning site C3# resulted in a 30-bp deletion in the corresponding coding region, leading to the possible formation of a truncated protein in the GGTA1 gene ^[52]^, which can still catalyze the synthesis of the αGal antigen. Therefore, during the second transfection, we finally succeeded in inactivating the GGTA1 gene by secondary targeting at exon 8 of the GGTA1 gene.

Currently, there is a lack of effective and safe clinical drugs or methods for monocyte inhibition in xenotransplantation. In a previous 8-GE pig to NHP kidney xenograft transplantation, varying degrees of renal failure with oliguria or anuria were observed at the end of the transplantation period, possibly due to extensive and severe glomerulosclerosis. The presence of a large number of macrophage aggregates in the glomeruli of transplanted porcine kidneys suggests that glomerulosclerosis may be related to the abnormal activation of macrophages ^[28]^. In the present study, overexpression of hCD47 in 8-GE pigs significantly reduced the number of macrophage infiltration in the glomeruli of transplanted porcine kidneys and prolonged the survival time of the grafts. In addition, while observing the both recipient (M1 and M2), a significantly lower number of macrophages in M1 was observed in the renal glomeruli of transplanted pigs than M2. This may be attributed to the dislodgement of the central venous retention tube in M2 on postoperative day 6, followed by a reintroduction of the central venous retention tube operation on postoperative day 8. The absence of immunosuppressant administration during this period may have been the direct cause of the surge in complement C3, C4, and monocyte counts in M2, which may also have contributed to the renal failure in M2. M1’s blood complement concentration and lymphocytes were consistently and effectively suppressed, but the gradual increase in creatinine concentration in the blood indicated progressive renal failure.M1 was suspected of having a gram-positive bacterial infection postoperative day 9-14, which was largely controlled by antibiotic therapy. From postoperative day 19 to the endpoint of the experiment, M1 detected signs of fungal infection, which may have contributed to the progressive failure of M1’s transplanted kidney.

IgM and IgG antibody deposition was observed in both porcine kidney grafts in this study, which was consistent with porcine heart xenografts utilizing the 10-GE pig developed by Revivicor ^[53]^, suggesting that porcine to NHP xenotransplantation is still characterized by the presence of other xenoantigens, leading to the development of antibody-mediated rejection. Contrarily, no significant AMR was found in multiple cases of multiple GE pig-to-human xenotransplantation in which three xenoantigens were knocked out ^[14, 54]^, which also suggests that NHP may not be an ideal model for studying the effectiveness of GE donor pigs in humans.

Biosafety risks are also an important impediment to clinical application of xenotransplantation. In human history, there has never been a shortage of viral invasions that have caused profound disasters for nations. For example, the HIV virus that spread globally in the 1980s infected more than 40 million people worldwide ^[55]^. In 2019, a global outbreak of COVID-19, with more than 700 million people infected and more than 7 million deaths due to COVID-19 infection ^[56]^. Porcine organ transplantation to humans also faces the transmission and spread of pathogenic microorganisms in the population, and only through strict screening and prevention and control measures during the breeding and production of donor pigs can avoid the spread of pathogenic microorganisms be. According to the WHO requirements for the control of pathogenic microorganisms in xenotransplantation donor pigs ^[37]^, we conducted multi-unit, multi-sampled, multi-method testing of 10-GE pigs that may carry most pathogenic microorganisms, and only found the presence of streptococcal infection, which can be completely cleared by the use of antibiotics, which indicates bio-secure production of our donor 10-GE pigs. However, a small number of pathogenic microorganisms cannot be determined at present due to the existence of many variants and lack of reliable detection methods. Therefore, the establishment of a sound screening method for pathogens screening in GE donor pigs is extremely important to promote the clinical application of xenotransplantation.

It is worthwhile to mention that on May 17, 2025, our 10 GEC pig liver was transplanted into a 71-year-old patient with massive liver cancer, making it the fifth case of xenotransplantation from a pig to a human patient worldwide. The patient is in good condition after the operation, and the pig liver can secrete bile in the human body without any signs of hyperacute or acute rejection ^[57]^, further confirming the effectiveness of our 10 gene-edited pigs.

In conclusion, we successfully obtained GTKO/CMAHKO/β4GalNT2KO/hCD46/hCD55/hCD59/hTBM/hCD39/hEPCR/hCD47 in 10-GE donor pigs by gene editing and somatic cell cloning techniques, which have normal reproductive ability with low biosafety risk and are in the pig to NHP. The effectiveness of the 10-GE pigs was confirmed by pig-to-NHP xenotransplantation, providing a usable donor pig for future clinical xenotransplantation.

## Supporting information

Supplementa fles

## Compliance with ethics guidelines

## Consent to participate

Informed consent was obtained from volunteer for being included in the study for obtaining serum sample.

## Acknowledgements

We thank the Xingdian Talent Support Program (XDYC-KJLJ-2022-0004), Department of Ministry of Science and Technology of Yunnan Province (202102AA310047; 202102AA100054) and Ministry of Science and Technology of the People’s Republic of China (2019YFA0110700) for providing funds.

## Data availability

All the data generated is available in the manuscript and can also be acquired from the corresponding authors upon special request.

## Authors Contribution

H-J.W. conceived the study. H-J.W. H-Y.Z. B-C.S. and T.L. designed and supervised the study. K.X. H.Z. B.J. J.W. N.A.M.A.S. A.M. K.L. W.C. C.Y. T.W. F.Z. X.H. D.J. J.G. H.Z. W.C. Y.Z. X.Z. L.J. Z.Z. W.Z. and T.L. performed the experiment. X.K., and M.A.J. wrote the manuscript. H-J.W. K.X and M.A.J. revised manuscript. All authors read the manuscript and approved for publication.

---

**Figure S1: Schematic diagram of generation of GEC pigs**. (A) Procedure of generation GEC pigs using gene-editing and SCNT technology. (B-C) designing of sgRNAs for KO of xenoantigens (B) and expression of human transgenes (C).

**Figure S2: Identification of 8-GEC pigs**

(A) PCR identification of 8-GEC pigs for KO of 3 xenoantigens (B) PCR identification of 8-GEC pigs for integration of 5 human transgenes. (C) Sanger sequencing of the targeting region of 3 xenoantigens in 8-GEC pigs. (D) The sequence of the targeting region of GGTA1, CMAH, and β4GalNT2 genes in 8-GEC pigs by Sanger sequencing.

**Figure S3: Cell colonies screening for TKO with integration of 8 transgenes, and generation and identification of 10-GEC fetuses**. (A-B) Screening of cell colonies for successful KO of α-Gal xenoantigen at exon 8 (A) and successful integration of 2 human transgenes (hEPCR and hCD47) into pig genome (B). (C) Photo of GEC fetuses obtained after 33-day pregnancy. (D) PCR identification of 10-GEC fetuses for KO of 3 xenoantigens. (E) PCR identification of 10-GEC fetuses for successful integration of 7 human transgenes by PCR (F), mRNA expression of 7 human transgenes in 10-GEC fetuses.

**Figure S4:** (A) Photo of 10-GEC pigs (B) The mRNA expression levels of 7 transgenes in heart, kidney, liver, and lung tissues of 10-GEC pigs. HUVEC, Human umbilical vein endothelial cell.

**Figure S5:** Changes of pig kidney, and liver before and after opening the blood vessel and heart transplant perfused and resuscitated during transplantation into rhesus monkeys.

**Figure S6**. (A) Bile production soon after liver transplant from 10-GEC pigs. (B)

(C-D) H&E staining of liver tissue with partial vascular endothelial breakdown and C4d deposition postoperative day 1. (E-F) H&E staining of liver tissue postoperative day 4 indicating necrotic disintegration.

**Figure S7** The liver functions indexes and electrolytes of recipient monkey

**Table S1** Primers for genotyping, qPCR and ddPCR

**Table S2** A summary of antibodies used by immunofluorescence, immunohistochemistry and flow cytometry

**Table S3** Genotyping the GGTA1, CMAH and β4GalNT2 genes of colonies by Sanger sequencing in the first gene editing.

**Table S4** The summary of genotype in cell colonies for 8 gene editing male pig production.

**Table S5** Genotyping the GGTA1, CMAH and β4GalNT2 genes of cloned fetuses with 8 gene editing by Sanger sequencing.

**Table S6** The generation of cloned pigs with 8 gene editing derived from cloned fetuses.

**Table S7** Genotyping the GGTA1, CMAH and β4GalNT2 genes of cloned pigs with 8 gene editing by Sanger sequencing.

**Table S8** The generation of cloned pigs with 8 gene editing.

**Table S9** Genotyping the GGTA1, CMAH and β4GalNT2 genes of cloned pigs with 8 gene editing by Sanger sequencing.

**Table S10** Genotyping the GGTA1 gene of colonies Sanger sequencing in the second gene editing.

**Table S11** Genotyping the GGTA1, CMAH and β4GalNT2 genes of cloned fetuses with 10 gene editing by Sanger sequencing.

**Table S12** The generation of cloned pigs with 10 gene editing.

**Table S13** Genotyping the GGTA1 gene of cloned pigs with 10 gene editing by Sanger sequencing.

**Table S14** Off-target analysis of 10-GEC pigs

**Table S15** Annotation information of transgenic insertion location

**Table S 16** The F1 generation of 10 GEC male (♂) pigs naturally mated with 8 GEC female (♀) pigs.

**Table S17** The genotype of F1 generation.

**Table S18** The summary of genotype in F1 generation of 10 GEC male pigs naturally mated with 8-GEC female pigs.

**Table S19** Screening of pathogenic microorganism in 10-GEC pigs

**Table S20** Information of recipient monkeys and donor pigs

