## Supplementa fles for "Specific Pathogen Free Ten Gene-Edited Pig Donor for Xenotransplantation": Supplementary Tables.docx

Table S1 Primers for genotyping, qPCR and ddPCR

| Primers | Sequence (5’-3’) | Size |
| --- | --- | --- |
| Genotyping | | |
| GGTA1-F | GGCAACATGGCAGGAAGGAA | 775bp |
| GGTA1-R | AGACGGCCCTGTCAGTTCAT |  |
| CMAH-F | AGCCTTAGAAGCCAGTGCAG | 904bp |
| CMAH-R | GCCCAAGCGAATCTACCTCA |  |
| β4GalNT2-F | TGCTCCATCGCAAGTGAC | 758bp |
| β4GalNT2-R | CCCGCAGTCAGGATTCAC |  |
| hCD55&hCD59-F | TCCAGCACCACCACAAATTGAC | 1124bp |
| hCD55&hCD59-R | CGGTGACCCGCTCGATGTG |  |
| hCD46-F | CTGATGAGACCCACAGAG | 100bp |
| hCD46-F | GCTCCACCATCTGCTTTC |  |
| hTBM&hCD39-F | GCTCGTGCATTCGGGCTTGC | 766bp |
| hTBM&hCD39-R | CCTGGCACCCTGGAAGTCAAAG |  |
| hTBM&hCD39-F | CCTTCCTCAATGCCAGTCAG | 2168bp |
| hTBM&hCD39-R | CCTGGCACCCTGGAAGTCAAAG |  |
| hEPCR-qPCR-F | GCTGAATGATCGTGGTGTTG | 140bp |
| hEPCR-qPCR-R | TGGCCTCCAAAGACTTCAT |  |
| hCD47PCR-F | GATGGATAAGAGTGATGCTGTC | 469bp |
| hCD47PCR-R | TCCAACCACAGCGAGGATATAG |  |
| qPCR | | |
| hCD46-F | ccaggtgcaggatcacaact | 146bp |
| hCD46-R | gcaatttggagcggtaagc |  |
| hCD55-F | TCCAGCACCACCACAAATTGAC | 207bp |
| hCD55-R | GGTGGGACCTTGGAAGTTAGAG |  |
| hCD59-F | TCATAGCCTGCAGTGCTACAAC | 103bp |
| hCD59-R | CCCAGCTTTGGTAATGAGACAC |  |
| hTBM-F | CATCCTGGACGACGGTTTCA | 107bp |
| hTBM-R | CGCAGATGCACTCGAAGGTA |  |
| hCD39-F | GGTGCCTATGGCTGGATTA | 211bp |
| hCD39-R | CCTTGCCATAGAGGCGAAA |  |
| hEPCR-qPCR-F | GCTGAATGATCGTGGTGTTG | 140bp |
| hEPCR-qPCR-R | TGGCCTCCAAAGACTTCAT |  |
| hCD47-qPCR-F | TTTCTCATCCATACCACCG | 229bp |
| hCD47-qPCR-R | GATGGATAAGAGTGATGCTGTC |  |
| pGAPDH-F | AGGGCATCCTGGGCTACACT | 367bp |
| pGAPDH-R | TCCACCACCCTGTTGCTGTAG |  |
| hGAPDH-F | GAGTCAACGGATTTGGTCGT | 160bp |
| hGAPDH-R | TGGAAGATGGTGATGGGATT |  |
| ddPCR | | |
| hCD46-F | CGTGGTCTCTTCTGCCTATTT | 95bp |
| hCD46-R | AAGGAAACTGAGACGCTACTG |  |
| hCD46 Probe | FAM TCCAGTGAAAGAAGCCAAGATCAGTAAGC |  |
| hCD55-F | GGTGCCAACAAGGCTAAATTC | 99bp |
| hCD55-R | CCTGGACGGCACTCATATTC |  |
| hCD55 Probe | FAM TGCATCCCTCAAACAGCCTTATATCACT |  |
| hCD59-F | CTGCAAGAAGGACCTGTGTAA | 93bp |
| hCD59-R | AATGGAGTCACCAGCAGAAG |  |
| hCD59 Probe | FAM AATGGTGGGACATCCTTATCAGAGAAA |  |
| hTBM-F | ACGTGGATGACTGCATACTG | 100bp |
| hTBM-R | ACCAGGTCGTAGTTAGGGTAG |  |
| hTBM Probe | FAM TCAACACACAGGGTGGCTTCGAG |  |
| hCD39-F | GCTCTGCAATTTCGCCTCTAT | 106bp |
| hCD39-R | GAATGTCCTTGGCCAGTTTCT |  |
| hCD39 Probe | FAM CTTCTTGTGCTATGGGAAGGATCAGGC |  |
| hCD47 -F | CAGCGATTGGATTAACCTCCT | 107bp |
| hCD47 -R | TGGTATACACGCCGCAATAC |  |
| hCD47 Probe | FAM CCTATATCCTCGCTGTGGTTGGACTGA |  |
| hEPCR -F | CAACCGCACTCGGTATGAA | 97bp |
| hEPCR -R | TTGGCTCCCTTTCGTGTTT |  |
| hEPCR Probe | FAM TTCTGCACATACTGCACACAGGTGTC |  |
| hGAPDH-F | CCTAGGGCTGCTCACATATTC | 85bp |
| hGAPDH-R | CGCCCAATACGACCAAATCTA |  |
| hGAPDH Probe | HEX CTCATGCCTTCTTGCCTCTTGTCTCT |  |
| pGAPDH-F | ccgcgatctaatgttctctttc | 114bp |
| pGAPDH-R | ttcactccgaccttcaccat |  |
| pGAPDH Probe | HEX cagccgcgtccctgagacac |  |

Table S2 A summary of antibodies used by immunofluorescence, immunohistochemistry and flow cytometry

|  | Primary antibody | | Secondary antibody | |
| --- | --- | --- | --- | --- |
|  | Catalog | Manufacturer | Catalog | Manufacturer |
| Immunofluorescence | | | | |
| GGTA1 | ALX-650-001F-MC05 | Enzo |  |  |
| CMAH(Neu5Gc) | 146903 | BioLegend | 550021 | Zenbio |
| B4GalNT2(DBA) | FL-1031 | Vector Labs |  |  |
| hCD46 | ab108307 | Abcam | GB21303 | Servicebio |
| hCD55 | ab230638 | Abcam | GB21303 | Servicebio |
| hCD55 | ab133684 | abcam | GB21303 | Servicebio |
| hCD59 | BF0017 | Affinity | GB21301 | Servicebio |
| hCD59 | ab9183 | abcam | GB21301 | Servicebio |
| hTBM(CD141) | sc-13164 | Santa Cruz | GB21301 | Servicebio |
| hCD39 | ab223842 | Abcam | GB21303 | Servicebio |
| hCD47 | ab226837 | Abcam | GB21303 | Servicebio |
| hEPCR | UM870092 | Origene | GB21301 | Servicebio |
| anti-CD3e | 85061T | CST | GB21303 | Servicebio |
| anti-CD4 | A22773 | ABClone |  |  |
| anti-CD8 | A23346 | ABClone |  |  |
| IgG | 62-8411 | Invitrogen |  |  |
| IgM | A18842 | Invitrogen |  |  |
| C3c | RAB-0027 | MXB Biotech. | GB21303 | Servicebio |
| C4d | RMA-0857 | MXB Biotech. | GB21303 | Servicebio |
| C5b-C9 | MA5-28502 | Invitrogen | GB21301 | Servicebio |
| Immunohistochemistry | | | | |
| CD57 | 14-0577-82 | Invitrogen | PV-9000 | OriGene |
| CD68 | 76437S | Cell Signaling | PV-9000 | OriGene |
| Flow cytometry | | | | |
| GGTA1 | ALX-650-001F-MC05 | Enzo |  |  |
| CMAH(Neu5Gc) | 146903 | BioLegend | 550021 | Zenbio |
| B4GalNT2(DBA) | FL-1031 | Vector Labs |  |  |
| hCD46 | 315304 | Biolegend |  |  |
| hCD55 | 555694 | BD |  |  |
| hCD59 | MA1-19463 | Invitrogen |  |  |
| hTBM | 564123 | BD |  |  |
| hCD39 | 560239 | BD |  |  |
| hCD47 | 556045 | BD |  |  |
| hEPCR | 557950 | BD |  |  |

**Table S3** Genotyping the GGTA1, CMAH and β4GalNT2 genes of colonies by Sanger sequencing in the first gene editing

| **Gene** | **No. of colonies** | **Sequence** | **Mutation** |
| --- | --- | --- | --- |
| GGTA1 | WT | TGATGTATTCCCAAAACACAACCATTACAGTTGAGACAAGCAGCATTGACAGAACCACTCTTCCTTTGACATTCATTATTTTCTCCTGGGAAAAGAAAAG |  |
|  | C2# | TGATGTATTCCCAAAACACAACCATTACAGTTGAGACAAGCAGCATTGACAGAACCACTCTTCCTT--------CATTATTTTCTCCTGGGAAAAGAAAAG  TGATGTATTCCCAAAACACAACCATTACAGTTGAGACAAGCAGCATTGACAGAACCACTCTTCC-----------ATTATTTTCTCCTGGGAAAAGAAAAG | -7bp 4/6  -1bp 2/6 |
|  | C3# | TGATGTATTCCCAAAACACAACCATTACAGTTGAGACAAGCAGCATTGACAGAACCACTCTTCCTTTGACATTCATTATTTTCTCCTGGGAAAAGAAAAG | WT 1/6 |
|  |  | TGATGTATTCCCAAAACACAACCATTACAGTTGAGACAAGCAGCAT-----------------------------------------------------G | -53bp 2/6 |
|  |  | TGATGTATTCCCAAAACACAACCATTACAGTTGAGACAAGCAGCATTGACAGAACCACTCTTCCTTTGACAAC-------------CCTGGGAAAAGAAAAG | -13bp/+2bp 2/6 |
|  | C5# | TGATGTATTCCCAAAACACAACCATTACAGTTGAGACAAGCAGCATTGACAGAACCACTCTTCCTTTGACATTCATTATTTTCTCCTGGGAAAAGAAAAG | WT 1/13 |
|  |  | TGATGTATTCCCAAAACACAACCATTACAGTTGAGACAAGCAGCATTGACAGAACCACTCTTCCTTTGTG---TTCATTATTTTCTCCTGGGAAAAGAAAAG | -3bp/+2bp 3/13 |
|  |  | TGATGTATTCCCAAAACACAACCATTACAGTTGAGACAAGCAGCATTGACAGAACCACTCTTCCTTT---------TATTTTCTCCTGGGAAAAGAAAAG | -9bp 1/13 |
|  |  | -----------------------------------------------------------------------------------ATTATTTTCTCCTGGGAAAAGAAAAG | -402bp 8/13 |
|  | C6# | TGATGTATTCCCAAAACACAACCATTACAGTTGAGACAAGCAGCATTGACAGAACCACTCTTCCTTTGACATTCATTATTTTCTCCTGGGAAAAGAAAAG | WT 1/4 |
|  |  | TGATGTATTCCCAAAACACAACCATTACAGTTGAGACAAGCAGCATTGACAGAACCACTCTTCCTTTGTG---TTCATTATTTTCTCCTGGGAAAAGAAAAG | -3bp/+2bp 2/4 |
|  |  | TGATGTATTCCCAAAACACAACCATTACAGTTGAGACAAGCAGCATTGACAGAACCACTCTTCCTTT---------TATTTTCTCCTGGGAAAAGAAAAG | -9bp 1/4 |
|  | C10# | TGATGTATTCCCAAAACACAACCATTACAGTTGAGACAAGCAGCATTGACAGAACCACTCTTCCTTTGACATTCATTATTTTCTCCTGGGAAAAGAAAAG | WT 2/8 |
|  |  | TGATGTATTCCCAAAACACAACCATTACAGTTGAGACAAGCAGCATTGACAGAACCACTCTTCCTT--------CATTATTTTCTCCTGGGAAAAGAAAAG | -7bp 2/8 |
|  |  | TGATGTATTCCCAAAACACAACCATTACAGTTGAGACAAGCAGCATTGACAGAACCACTCTTCC-----------ATTATTTTCTCCTGGGAAAAGAAAAG | -10bp 4/8 |
|  | C21# | TGATGTATTCCCAAAACACAACCATTACAGTTGAGACAAGCAGCATTGACAGAACCACTCTTCCTTTGACATTCATTATTTTCTCCTGGGAAAAGAAAAG | WT 6/11 |
|  |  | -----------------------------------------------------------------------------------------TTCTCCTGGGAAAAGAAAAG | -378bp 5/11 |
|  | C24# | TGATGTATTCCCAAAACACAACCATTACAGTTGAGACAAGCAGCATTGACAGAACCACTCTTCCTTTGACATTCATTATTTTCTCCTGGGAAAAGAAAAG | WT 3/7 |
|  |  | TGATGTATTCCCAAAACACAACCATTACAGTTGAGACAAGCAGCATTGACAGAACCACTCTTCCTT--------CATTATTTTCTCCTGGGAAAAGAAAAG | -7bp 2/7 |
|  |  | TGATGTATTCCCAAAACACAACCATTACAGTTGAGACAAGCAGCATTGACAGAACCACTCTTCCTTTGAACATTCATTATTTTCTCCTGGGAAAAGAAAAG | +1bp 2/7 |
|  | C26# | TGATGTATTCCCAAAACACAACCATTACAGTTGAGACAAGCAGCATTGACAGAACCACTCTTCCTTTGACATTCATTATTTTCTCCTGGGAAAAGAAAAG | WT 3/6 |
|  |  | TGATGTATTCCCAAAACACAACCATTACAGTTGAGACAAGCAGCATTGACAGAACCACTCTTCCTTTGA-ATTCATTATTTTCTCCTGGGAAAAGAAAAG | -1bp 3/6 |
|  | C34# | TGATGTATTCCCAAAACACAACCATTACAGTTGAGACAAGCAGCATTGACAGAACCACTCTTCCTTTGACATTCATTATTTTCTCCTGGGAAAAGAAAAG | WT 1/6 |
|  |  | TGATGTATTCCCAAAACACAACCATTACAGTTGAGACAAGCAGCATTGACAGAACCACTCTTCCTTTGA--TTCATTATTTTCTCCTGGGAAAAGAAAAG | -2bp 3/6 |
|  |  | TGATGTATTCCCAAAACACAACCATTACAGTTGAGACAAGCAGCATTGACAGAACCACTCTTCCTTTGAACATTCATTATTTTCTCCTGGGAAAAGAAAAG | **+**1bp 2/6 |
|  | C44# | TGATGTATTCCCAAAACACAACCATTACAGTTGAGACAAGCAGCATTGACAGAACCACTCTTCCTTTGACATTCATTATTTTCTCCTGGGAAAAGAAAAG | WT 5/11 |
|  |  | TGATGTATTCCCAAAA-----------------------------------------------------------ACATTCATTATTTTCTCCTGGGAAAAGAAAAG | -52bp 1/11 |
|  |  | -----------------------------------------------------------------------------------------TTCTCCTGGGAAAAGAAAAG | -380bp 5/11 |
| CMAH | WT | TCACTGTCTTCCAACAGATCACGTACCTTACTCACG(84)TACTACACGAGCCTCCATCTGATTGG(62)GTAAGGAAGGGTGAGCCCTCAACTCCGAAGA |  |
|  | C2# | TCACTGTCTTCCAACAGATCACGTACCTTACTCACG(84)TACTACACGAGCCTCCATCTGATTGG(62)GTAAGGAAGGGTGAGCCCTCAACTCCGAAGA | WT 3/7 |
|  |  | TCACTGTCTT--------------------------**(84)**-------------------TGATTGG(62)GTAAGGAAGGGTGAGCCCTCAACTCCGAAGA | -123 4/7 |
|  | C3# | TCACTGTCTTCCAACAGATCACGTACCTTACTCACG(84)TACTACACGAGCCTCCATCTGATTGG(62)GTAAGGAAGGGTGAGCCCTCAACTCCGAAGA | WT 7/18 |
|  |  | TCACTGTCTT--------------------------**(84)**-------------------TGATTGG(62)GTAAGGAAGGGTGAGCCCTCAACTCCGAAGA | -123bp 5/18 |
|  |  | TCACTGTCTTCCAACA----------CTTACTCACG(84)TACTACACGAGCCTCCA---------**(62)**---------------------ACTCCGAAGA | 102bp 5/18 |
|  |  | TCACTGTCTTCCAACAGATCACGTACCTTACTCACG(84)TACTACACGAGCCTCCA---------**(62)**---------------------ACTCCGAAGA | 92bp 1/18 |
|  | C10# | TCACTGTCTTCCAACAGATCACGTACCTTACTCACG(84)TACTACACGAGCCTCCATCTGATTGG(62)GTAAGGAAGGGTGAGCCCTCAACTCCGAAGA | WT 2/7 |
|  |  | TCACTGTCTT--------------------------**(84)**-------------------TGATTGG(62)GTAAGGAAGGGTGAGCCCTCAACTCCGAAGA | -123bp 5/7 |
|  | C24# | TCACTGTCTTCCAACAGATCACGTACCTTACTCACG(84)TACTACACGAGCCTCCATCTGATTGG(62)GTAAGGAAGGGTGAGCCCTCAACTCCGAAGA | WT 2/8 |
|  |  | TCACTGTCTT--------------------------**(84)**-------------------TGATTGG(62)GTAAGGAAGGGTGAGCCCTCAACTCCGAAGA | -123bp 6/8 |
| B4GalNT2 | WT | AGCCCTAGATGTCTGTCGATCCTCAAGAT(99)CACCCTGGATGCGCAGACGCTGAAGCTTCTACC(15)CTACGGTGAAAACGGGTGAGATGGCAA |  |
|  |  | AGCCCTAGATGTCTGTCGATCCTCAAGAT(99)CACCCTGGATGCGCAGACGCTGAAGCTTCTACC(27)CTACAGTGGAATCTGGTGAGATGGCAA |  |
|  | C2# | AGCCCTAGATGTCTGTCGATCCTCAAGAT(99)CACCCTGGATGCGCAGACGCTGAAGCTTCTACC(27)CTACAGTGGAATCTGGTGAGATGGCAA | WT 1/4 |
|  |  | AGCCCTAGAT-------------------**(99)**---------------------------------**(15)**------------------AGATGGCAA | -184bp 3/4 |
|  | C3# | AGCCCTAGATGTCTGTCGATCCTCAAGAT(99)CACCCTGGATGCGCAGACGCTGAAGCTTCTACC(15)CTACGGTGAAAACGGGTGAGATGGCAA | WT 3/9 |
|  |  | AGCCCTAGATGTCTGTCGATCCTCAAGAT(99)CACCCTGGATGCGCAGACGCTGAAGCTTCTACC(27)CTACAGTGGAATCTGGTGAGATGGCAA | WT 2/9 |
|  |  | AGCCCTAGATG--------------------**(99)**--------------------------------**(15)**-----------------AGATGGCAA | -183bp 2/9 |
|  |  | AGCCCTAGATG------------------**(99)**----------------------AAGCTTCTACC(27)CTACAGTGGAATCTGGTGAGATGGCAA | -139bp 2/9 |
|  | C10# | AGCCCTAGATGTCTGTCGATCCTCAAGAT(99)CACCCTGGATGCGCAGACGCTGAAGCTTCTACC(15)CTACGGTGAAAACGGGTGAGATGGCAA | WT 1/10 |
|  |  | AGCCCTAGATGTCTGTCGATCCTCAAGAT(99)CACCCTGGATGCGCAGACGCTGAAGCTTCTACC(27)CTACAGTGGAATCTGGTGAGATGGCAA | WT 1/10 |
|  |  | AGCCCTAGATG--------------------**(99)**-------------------------------**(15)**-----------------AGATGGCAA | -183bp 4/10 |
|  |  | AGCCCTAGATGTCTGTCGATCCTCAAGAT(99)CACCCTGGAT-----------------------**(15)**-----------------AAGATGGCAA | -61bp 1/10 |
|  |  | AGCCCTAGATGTCTGTCGATCCTCAAGAT(99)CACCCTGGATGCGCAGACGCTGAAGCTTCTACC(15)CTACGGTGAA---------GATGGCAA | -9bp 3/10 |
|  | C12# | AGCCCTAGATGTCTGTCGATCCTCAAGAT(99)CACCCTGGATGCGCAGACGCTGAAGCTTCTACC(15)CTACGGTGAAAACGGGTGAGATGGCAA | WT 2/6 |
|  |  | AGCCCTAGATGTCTGTCGATCCTCAAGAT(99)CACCCTGGATGCGCAGACGCTGAAGCTTCTACC(27)CTACAGTGGAATCTGGTGAGATGGCAA | WT 1/6 |
|  |  | AGCCCTAGATG--------------------**(99)**-------------------------------**(15)**-----------------AGATGGCAA | -183bp 2/6 |
|  |  | AGCCCTAGATGTCTGTCGATCCTCAAGAT(99)CACCCTGGATGCGCAGACGCTGAAGCTTCTACC(15)CTACGGTGAA---------GATGGCAA | -9bp 1/6 |
|  | C21# | AGCCCTAGATGTCTGTCGATCCTCAAGAT(99)CACCCTGGATGCGCAGACGCTGAAGCTTCTACC(15)CTACGGTGAAAACGGGTGAGATGGCAA | WT 2/4 |
|  |  | AGCCCTAGATGTCTGTCGATCCTCAAGAT(99)CACCCTGGATGCGCAGACGCTGAAGCTTCTACC(27)CTACAGTGGAATCTGGTGAGATGGCAA | WT 1/4 |
|  |  | AGCCCTAGATG--------------------**(99)**-------------------------------**(15)**-----------------AGATGGCAA | -183bp 1/4 |
|  | C24# | AGCCCTAGATGTCTGTCGATCCTCAAGAT(99)CACCCTGGATGCGCAGACGCTGAAGCTTCTACC(27)CTACAGTGGAATCTGGTGAGATGGCAA | WT 5/7 |
|  |  | AGCCCTAGATG--------------------**(99)**-------------------------------**(15)**-----------------AGATGGCAA | -183bp 2/7 |

**Table S4 The summary of genotype in cell colonies for 8 gene editing male pig production**

| No. | ID | GGTA1 | CMAH | β4GalNT2 | hCD46 | hCD55 | hCD59 | hTBM | hCD39 |
| --- | --- | --- | --- | --- | --- | --- | --- | --- | --- |
| 1 | C2 | +1/△7/△10/+2△9 | WT/△123 | WT/WT/△184/△184 | ○ | ○ | ○ | ○ | ○ |
| 2 | C3 | WT/△53/+2△13 | WT/△123/△102/△92 | WT/△139/WT/△183 | ○ | ○ | ○ | ○ | ○ |
| 3 | C5 | WT/+2△3/△9/△402 | - | WT | ○ | ○ | ○ | ○ | ○ |
| 4 | C6 | WT/△10/+1△3 | - | WT | ○ | ○ | ○ | ○ | ○ |
| 5 | C10 | WT/△10/△7 | WT/△123 | WT/WT/△9/+2△59/△184 | ○ | ○ | ○ | ○ | ○ |
| 6 | C12 | WT/△10/△7 | - | WT/WT/△9/△184 | ○ | ○ | ○ | ○ | ○ |
| 7 | C21 | WT/△378 | - | WT/WT/△182/△182 | ○ | ○ | ○ | ○ | ○ |
| 8 | C24 | WT/+1/+11△2/△7 | WT/△123 | WT/△119/△183/△183 | ○ | ○ | ○ | ○ | ○ |
| 9 | C26 | WT/△1/△1 | - | WT | ○ | ○ | ○ | ○ | ○ |
| 10 | C34 | WT/+2/△1 | - | WT | ○ | ○ | ○ | ○ | ○ |
| 11 | C44 | WT/△52/△380 | - | WT | ○ | ○ | ○ | ○ | ○ |
| 12 | C4 | WT | - | - | ○ | ○ | ○ | ○ | ○ |
| 13 | C8 | WT | - | - | ○ | ○ | ○ | ○ | ○ |
| 14 | C9 | WT | - | - | ○ | ○ | ○ | ○ | ○ |
| 15 | C11 | WT | - | - | ○ | ○ | ○ | ○ | ○ |
| 16 | C13 | WT | - | - | ○ | ○ | ○ | ○ | ○ |
| 17 | C14 | WT | - | - | ○ | ○ | ○ | ○ | ○ |
| 18 | C15 | WT | - | - | ○ | ○ | ○ | ○ | ○ |
| 19 | C17 | WT | - | - | ○ | ○ | ○ | ○ | ○ |
| 20 | C18 | WT | - | - | ○ | ○ | ○ | ○ | ○ |
| 21 | C19 | WT | - | - | ○ | ○ | ○ | ○ | ○ |
| 22 | C20 | WT | - | - | ○ | ○ | ○ | ○ | ○ |
| 23 | C22 | WT | - | - | ○ | ○ | ○ | ○ | ○ |
| 24 | C27 | WT | - | - | ○ | ○ | ○ | ○ | ○ |
| 25 | C28 | WT | - | - | ○ | ○ | ○ | ○ | ○ |
| 26 | C29 | WT | - | - | ○ | ○ | ○ | ○ | ○ |
| 27 | C31 | WT | - | - | ○ | ○ | ○ | ○ | ○ |
| 28 | C32 | WT | - | - | ○ | ○ | ○ | ○ | ○ |
| 29 | C33 | WT | - | - | ○ | ○ | ○ | ○ | ○ |
| 30 | C35 | WT | - | - | ○ | ○ | ○ | ○ | ○ |
| 31 | C39 | WT | - | - | ○ | ○ | ○ | ○ | ○ |
| 32 | C48 | Sequencing Defeat | - | - | ○ | ○ | ○ | ○ | ○ |
| 33 | Remaining 17 colonies | - | - | - | × | × | × | × | × |

Note: A total of 49 cell colonies were screened, among which C2, C3, C4, C5, C6, C8, C9, C10, C11, C12, C13, C14, C15, C17, C18, C19, C20, C21, C22, C24, C26, C27, C28, C29, C31, C32, C33, C34, C35, C39, C44, C48 (a total of 32) were PCR-positive cell colonies, indicating successful insertion of hTBM/hCD39/hCD46/hCD55/hCD59. These colonies were used for GGTA1 gene target fragment PCR and Sanger sequencing, and the mutated colonies are shown in the table. Then, β4GalNT2 gene target fragment PCR and Sanger sequencing were performed, and the mutated colonies were C2, C3, C10, C12, C21, and C24 (a total of 6). Among them, C12 colony was a single knockout and existed in multiples of 3 (△9), and C21 clone was confirmed to be a single knockout for both GGTA1 and beta4GalNT2. Finally, Sanger sequencing was performed on the CMAH gene target fragment for C2, C3, C10, and C24 clones (a total of 4). (“-”: not tested; “○”: insertion detected; “×”: no insertion detected)

**Table S5** Genotyping the GGTA1, CMAH and β4GalNT2 genes of cloned fetuses with 8 gene editing by Sanger sequencing

| **Gene** | **No. of pigs** | **Sequence** | **Mutation** |
| --- | --- | --- | --- |
| GGTA1  On exon 1 | WT | TGATGTATTCCCAAAACACAACCATTACAGTTGAGACAAGCAGCATTGACAGAACCACTCTTCCTTTGACATTCATTATTTTCTCCTGGGAAAAGAAAAG |  |
|  | C3F01 | TGATGTATTCCCAAAACACAACCATTACAGTTGAGACAAGCAGCATTGACAGAACCACTCTTCCTTTGACAAC-------------CCTGGGAAAAGAAAAG | -13bp/+2bp 4/10 |
|  |  | TGATGTATTCCCAAAACACAACCATTACAGTTGAGACAAGCAGCAT-----------------------------------------------------G | -53bp 6/10 |
|  | C3F02 | TGATGTATTCCCAAAACACAACCATTACAGTTGAGACAAGCAGCATTGACAGAACCACTCTTCCTTTGACAAC-------------CCTGGGAAAAGAAAAG | -13bp/+2bp 5/8 |
|  |  | TGATGTATTCCCAAAACACAACCATTACAGTTGAGACAAGCAGCAT-----------------------------------------------------G | -53bp 3/8 |
|  | C3F03 | TGATGTATTCCCAAAACACAACCATTACAGTTGAGACAAGCAGCATTGACAGAACCACTCTTCCTTTGACAAC-------------CCTGGGAAAAGAAAAG | -13bp/+2bp 4/6 |
|  |  | TGATGTATTCCCAAAACACAACCATTACAGTTGAGACAAGCAGCAT-----------------------------------------------------G | -53bp 2/6 |
|  | C3F04 | TGATGTATTCCCAAAACACAACCATTACAGTTGAGACAAGCAGCATTGACAGAACCACTCTTCCTTTGACAAC-------------CCTGGGAAAAGAAAAG | -13bp/+2bp 4/7 |
|  |  | TGATGTATTCCCAAAACACAACCATTACAGTTGAGACAAGCAGCAT-----------------------------------------------------G | -53bp 3/7 |
|  | C3F05 | TGATGTATTCCCAAAACACAACCATTACAGTTGAGACAAGCAGCATTGACAGAACCACTCTTCCTTTGACAAC-------------CCTGGGAAAAGAAAAG | -13bp/+2bp 10/15 |
|  |  | TGATGTATTCCCAAAACACAACCATTACAGTTGAGACAAGCAGCAT-----------------------------------------------------G | -53bp 5/15 |
|  | C3F06 | TGATGTATTCCCAAAACACAACCATTACAGTTGAGACAAGCAGCATTGACAGAACCACTCTTCCTTTGACAAC-------------CCTGGGAAAAGAAAAG | -13bp/+2bp 3/7 |
|  |  | TGATGTATTCCCAAAACACAACCATTACAGTTGAGACAAGCAGCAT-----------------------------------------------------G | -53bp 4/7 |
|  | C3F07 | TGATGTATTCCCAAAACACAACCATTACAGTTGAGACAAGCAGCATTGACAGAACCACTCTTCCTTTGACAAC-------------CCTGGGAAAAGAAAAG | -13bp/+2bp 3/9 |
|  |  | TGATGTATTCCCAAAACACAACCATTACAGTTGAGACAAGCAGCAT-----------------------------------------------------G | -53bp 6/9 |
|  | C3F08 | TGATGTATTCCCAAAACACAACCATTACAGTTGAGACAAGCAGCATTGACAGAACCACTCTTCCTTTGACAAC-------------CCTGGGAAAAGAAAAG | -13bp/+2bp 5/9 |
|  |  | TGATGTATTCCCAAAACACAACCATTACAGTTGAGACAAGCAGCAT-----------------------------------------------------G | -53bp 4/9 |
|  | C3F09 | TGATGTATTCCCAAAACACAACCATTACAGTTGAGACAAGCAGCATTGACAGAACCACTCTTCCTTTGACAAC-------------CCTGGGAAAAGAAAAG | -13bp/+2bp 8/11 |
|  |  | TGATGTATTCCCAAAACACAACCATTACAGTTGAGACAAGCAGCAT-----------------------------------------------------G | -53bp 11/11 |
|  | C3F10 | TGATGTATTCCCAAAACACAACCATTACAGTTGAGACAAGCAGCATTGACAGAACCACTCTTCCTTTGACAAC-------------CCTGGGAAAAGAAAAG | -13bp/+2bp 3/8 |
|  |  | TGATGTATTCCCAAAACACAACCATTACAGTTGAGACAAGCAGCAT-----------------------------------------------------G | -53bp 5/8 |
|  | C3F11 | TGATGTATTCCCAAAACACAACCATTACAGTTGAGACAAGCAGCATTGACAGAACCACTCTTCCTTTGACAAC-------------CCTGGGAAAAGAAAAG | -13bp/+2bp 8/9 |
|  |  | TGATGTATTCCCAAAACACAACCATTACAGTTGAGACAAGCAGCAT-----------------------------------------------------G | -53bp 1/9 |
|  | C3F12 | TGATGTATTCCCAAAACACAACCATTACAGTTGAGACAAGCAGCATTGACAGAACCACTCTTCCTTTGACAAC-------------CCTGGGAAAAGAAAAG | -13bp/+2bp 5/10 |
|  |  | TGATGTATTCCCAAAACACAACCATTACAGTTGAGACAAGCAGCAT-----------------------------------------------------G | -53bp 5/10 |
|  | C3F13 | TGATGTATTCCCAAAACACAACCATTACAGTTGAGACAAGCAGCATTGACAGAACCACTCTTCCTTTGACAAC-------------CCTGGGAAAAGAAAAG | -13bp/+2bp 6/9 |
|  |  | TGATGTATTCCCAAAACACAACCATTACAGTTGAGACAAGCAGCAT-----------------------------------------------------G | -53bp 3/9 |
| CMAH | WT | TCACTGTCTTCCAACAGATCACGTACCTTACTCACG(84)TACTACACGAGCCTCCATCTGATTGG(62)GTAAGGAAGGGTGAGCCCTCAACTCCGAAGA |  |
|  | C3F01 | TCACTGTCTT--------------------------**(84)**-------------------TGATTGG(62)GTAAGGAAGGGTGAGCCCTCAACTCCGAAGA | -123bp 6/9 |
|  |  | TCACTGTCTTCCAACA----------CTTACTCACG(84)TACTACACGAGCCTCCA---------**(62)**---------------------ACTCCGAAGA | -102bp 3/9 |
|  | C3F02 | TCACTGTCTT--------------------------**(84)**-------------------TGATTGG(62)GTAAGGAAGGGTGAGCCCTCAACTCCGAAGA | -123bp 6/8 |
|  |  | TCACTGTCTTCCAACA----------CTTACTCACG(84)TACTACACGAGCCTCCA---------**(62)**---------------------ACTCCGAAGA | -102b 2/8 |
|  | C3F03 | TCACTGTCTT--------------------------**(84)**-------------------TGATTGG(62)GTAAGGAAGGGTGAGCCCTCAACTCCGAAGA | -123bp 3/8 |
|  |  | TCACTGTCTTCCAACA----------CTTACTCACG(84)TACTACACGAGCCTCCA---------**(62)**---------------------ACTCCGAAGA | -102b 5/8 |
|  | C3F04 | TCACTGTCTT--------------------------**(84)**-------------------TGATTGG(62)GTAAGGAAGGGTGAGCCCTCAACTCCGAAGA | -123bp 2/8 |
|  |  | TCACTGTCTTCCAACA----------CTTACTCACG(84)TACTACACGAGCCTCCA---------**(62)**---------------------ACTCCGAAGA | -102b 6/8 |
|  | C3F05 | TCACTGTCTT--------------------------**(84)**-------------------TGATTGG(62)GTAAGGAAGGGTGAGCCCTCAACTCCGAAGA | -123bp 3/10 |
|  |  | TCACTGTCTTCCAACA----------CTTACTCACG(84)TACTACACGAGCCTCCA---------**(62)**---------------------ACTCCGAAGA | -102bp 7/10 |
|  | C3F06 | TCACTGTCTT--------------------------**(84)**-------------------TGATTGG(62)GTAAGGAAGGGTGAGCCCTCAACTCCGAAGA | -123bp 7/11 |
|  |  | TCACTGTCTTCCAACA----------CTTACTCACG(84)TACTACACGAGCCTCCA---------**(62)**---------------------ACTCCGAAGA | -102bp 4/11 |
|  | C3F07 | TCACTGTCTT--------------------------**(84)**-------------------TGATTGG(62)GTAAGGAAGGGTGAGCCCTCAACTCCGAAGA | -123bp 6/11 |
|  |  | TCACTGTCTTCCAACA----------CTTACTCACG(84)TACTACACGAGCCTCCA---------**(62)**---------------------ACTCCGAAGA | -102bp 6/11 |
|  | C3F08 | TCACTGTCTT--------------------------**(84)**-------------------TGATTGG(62)GTAAGGAAGGGTGAGCCCTCAACTCCGAAGA | -123bp 7/11 |
|  |  | TCACTGTCTTCCAACA----------CTTACTCACG(84)TACTACACGAGCCTCCA---------**(62)**---------------------ACTCCGAAGA | -102bp 4/11 |
|  | C3F09 | TCACTGTCTT--------------------------**(84)**-------------------TGATTGG(62)GTAAGGAAGGGTGAGCCCTCAACTCCGAAGA | -123bp 5/8 |
|  |  | TCACTGTCTTCCAACA----------CTTACTCACG(84)TACTACACGAGCCTCCA---------**(62)**---------------------ACTCCGAAGA | -102bp 3/8 |
|  | C3F10 | TCACTGTCTT--------------------------**(84)**-------------------TGATTGG(62)GTAAGGAAGGGTGAGCCCTCAACTCCGAAGA | -123bp 5/10 |
|  |  | TCACTGTCTTCCAACA----------CTTACTCACG(84)TACTACACGAGCCTCCA---------**(62)**---------------------ACTCCGAAGA | -102bp 5/10 |
|  | C3F11 | TCACTGTCTT--------------------------**(84)**-------------------TGATTGG(62)GTAAGGAAGGGTGAGCCCTCAACTCCGAAGA | -123bp 6/12 |
|  |  | TCACTGTCTTCCAACA----------CTTACTCACG(84)TACTACACGAGCCTCCA---------**(62)**---------------------ACTCCGAAGA | -102bp 6/12 |
|  | C3F12 | TCACTGTCTT--------------------------**(84)**-------------------TGATTGG(62)GTAAGGAAGGGTGAGCCCTCAACTCCGAAGA | -123bp 5/8 |
|  |  | TCACTGTCTTCCAACA----------CTTACTCACG(84)TACTACACGAGCCTCCA---------**(62)**---------------------ACTCCGAAGA | -102bp 3/8 |
|  | C3F13 | TCACTGTCTT--------------------------**(84)**-------------------TGATTGG(62)GTAAGGAAGGGTGAGCCCTCAACTCCGAAGA | -123bp 5/11 |
|  |  | TCACTGTCTTCCAACA----------CTTACTCACG(84)TACTACACGAGCCTCCA---------**(62)**---------------------ACTCCGAAGA | -102bp 6/11 |
| β4GalNT2 | WT | AGCCCTAGATGTCTGTCGATCCTCAAGAT(99)CACCCTGGATGCGCAGACGCTGAAGCTTCTACC(15)CTACGGTGAAAACGGGTGAGATGGCAA |  |
|  |  | AGCCCTAGATGTCTGTCGATCCTCAAGAT(99)CACCCTGGATGCGCAGACGCTGAAGCTTCTACC(27)CTACAGTGGAATCTGGTGAGATGGCAA |  |
|  | C3F01 | AGCCCTAGATG--------------------**(99)**--------------------------------**(15)**-----------------AGATGGCAA | -183bp 6/11 |
|  |  | AGCCCTAGATG------------------**(99)**----------------------AAGCTTCTACC(27)CTACAGTGGAATCTGGTGAGATGGCAA | -139bp 5/11 |
|  | C3F02 | AGCCCTAGATG--------------------**(99)**--------------------------------**(15)**-----------------AGATGGCAA | -183bp 2/9 |
|  |  | AGCCCTAGATG------------------**(99)**----------------------AAGCTTCTACC(27)CTACAGTGGAATCTGGTGAGATGGCAA | -139bp 7/9 |
|  | C3F03 | AGCCCTAGATG--------------------**(99)**--------------------------------**(15)**-----------------AGATGGCAA | -183bp 3/7 |
|  |  | AGCCCTAGATG------------------**(99)**----------------------AAGCTTCTACC(27)CTACAGTGGAATCTGGTGAGATGGCAA | -139bp 4/7 |
|  | C3F04 | AGCCCTAGATG--------------------**(99)**--------------------------------**(15)**-----------------AGATGGCAA | -183bp 2/5 |
|  |  | AGCCCTAGATG------------------**(99)**----------------------AAGCTTCTACC(27)CTACAGTGGAATCTGGTGAGATGGCAA | -139bp 3/5 |
|  | C3F05 | AGCCCTAGATG--------------------**(99)**--------------------------------**(15)**-----------------AGATGGCAA | -183bp 4/7 |
|  |  | AGCCCTAGATG------------------**(99)**----------------------AAGCTTCTACC(27)CTACAGTGGAATCTGGTGAGATGGCAA | -139bp 3/7 |
|  | C3F06 | AGCCCTAGATG--------------------**(99)**--------------------------------**(15)**-----------------AGATGGCAA | -183bp 4/7 |
|  |  | AGCCCTAGATG------------------**(99)**----------------------AAGCTTCTACC(27)CTACAGTGGAATCTGGTGAGATGGCAA | -139bp 3/7 |
|  | C3F07 | AGCCCTAGATG--------------------**(99)**--------------------------------**(15)**-----------------AGATGGCAA | -183bp 5/9 |
|  |  | AGCCCTAGATG------------------**(99)**----------------------AAGCTTCTACC(27)CTACAGTGGAATCTGGTGAGATGGCAA | -139bp 4/9 |
|  | C3F08 | AGCCCTAGATG--------------------**(99)**--------------------------------**(15)**-----------------AGATGGCAA | -183bp 6/11 |
|  |  | AGCCCTAGATG------------------**(99)**----------------------AAGCTTCTACC(27)CTACAGTGGAATCTGGTGAGATGGCAA | -139bp 5/11 |
|  | C3F09 | AGCCCTAGATG--------------------**(99)**--------------------------------**(15)**-----------------AGATGGCAA | -183bp 5/9 |
|  |  | AGCCCTAGATG------------------**(99)**----------------------AAGCTTCTACC(27)CTACAGTGGAATCTGGTGAGATGGCAA | -139bp 4/9 |
|  | C3F10 | AGCCCTAGATG--------------------**(99)**--------------------------------**(15)**-----------------AGATGGCAA | -183bp 6/11 |
|  |  | AGCCCTAGATG------------------**(99)**----------------------AAGCTTCTACC(27)CTACAGTGGAATCTGGTGAGATGGCAA | -139bp 5/11 |
|  | C3F11 | AGCCCTAGATG--------------------**(99)**--------------------------------**(15)**-----------------AGATGGCAA | -183bp 4/8 |
|  |  | AGCCCTAGATG------------------**(99)**----------------------AAGCTTCTACC(27)CTACAGTGGAATCTGGTGAGATGGCAA | -139bp 4/8 |
|  | C3F12 | AGCCCTAGATG--------------------**(99)**--------------------------------**(15)**-----------------AGATGGCAA | -183bp 5/10 |
|  |  | AGCCCTAGATG------------------**(99)**----------------------AAGCTTCTACC(27)CTACAGTGGAATCTGGTGAGATGGCAA | -139bp 5/10 |
|  | C3F13 | AGCCCTAGATG--------------------**(99)**--------------------------------**(15)**-----------------AGATGGCAA | -183bp 7/9 |
|  |  | AGCCCTAGATG------------------**(99)**----------------------AAGCTTCTACC(27)CTACAGTGGAATCTGGTGAGATGGCAA | -139bp 2/9 |

**Table S6** The generation of cloned pigs with 8 gene editing derived from cloned fetuses

| No. | ID | Body weight (g) | Status at birth | Survival time (day) | Surrogate mother |
| --- | --- | --- | --- | --- | --- |
| 1 | 33dDN-8TGC3F01P01 | 100 | mummy | 0 | P754 |
| 2 | 33dDN-8TC3GF01P02 | 1000 | healthy | 130 |  |
| 3 | 33dDN-8TGC3F01P03 | 1050 | healthy | - |  |
| 4 | 33dDN-8TGC3F01P04 | 600 | stillbirth | 0 |  |
| 5 | 33dDN-8TGC3F01P05 | 1120 | stillbirth | 0 |  |
| 6 | 33dDN-8TGC3F01P06 | 500 | healthy | 3 | P774 |
| 7 | 33dDN8TGC3F03P01 | 850 | healthy | 5 | P814 |
| 8 | 33dDN8TGC3F03P02 | 800 | healthy | 3 |  |
| 9 | 33dDN8TGC3F03P03 | 500 | stillbirth | 0 |  |
| 10 | 33dDN8TGC3F03P04 | 850 | healthy | 9 |  |
| 11 | 33dDN8TGC3F03P05 | 800 | healthy | 3 |  |
| 12 | 33dDN8TGC3F03P06 | 700 | stillbirth | 0 |  |
| 13 | 33dDN8TGC3F03P07 | 850 | stillbirth | 0 |  |
| 14 | 33dDN8TGC3F03P08 | 650 | healthy | 96 | P738 |
| 15 | 33dDN8TGC3F03P09 | 800 | healthy | - |  |
| 16 | 33dDN8TGC3F03P10 | 850 | healthy | 89 |  |
| 17 | 33dDN8TGC3F03P11 | 650 | healthy | 55 |  |
| 18 | 33dDN8TGC3F07P08 | 500 | healthy | 2 | P832 |
| 19 | 33dDN8TGC3F07P09 | 600 | healthy | - |  |
| 20 | 33dDN8TGC3F07P10 | 450 | stillbirth | 0 |  |
| 21 | 33dDN8TGC3F07P11 | 600 | stillbirth | 0 |  |
| 22 | 33dDN8TGC3F07P12 | 400 | stillbirth | 0 |  |
| 23 | 33dDN8TGC3F07P13 | 450 | stillbirth | 0 |  |
| 24 | 33dDN8TGC3F07P14 | 500 | stillbirth | 0 |  |
| 25 | 33dDN8TGC3F07P15 | 700 | stillbirth | 0 |  |
| 26 | 33dDN8TGC3F07P16 | 250 | mummy | 0 |  |
| 27 | 33dDN8TGC3F07P01 | 650 | healthy | 2 | P856 |
| 28 | 33dDN8TGC3F07P02 | 750 | healthy | 2 |  |
| 29 | 33dDN8TGC3F07P03 | 800 | healthy | 7 |  |
| 30 | 33dDN8TGC3F07P04 | 200 | mummy | 0 | P757 |
| 31 | 33dDN8TGC3F07P05 | 250 | stillbirth | 0 |  |
| 32 | 33dDN8TGC3F07P06 | 400 | stillbirth | 0 |  |
| 33 | 33dDN8TGC3F07P07 | 240 | mummy | 0 |  |
| 34 | 33dDN8TGC3F07P17 | 600 | stillbirth | 0 |  |
| 35 | 33dDN8TGC3F07P18 | 300 | mummy | 0 |  |
| 36 | 33dDN8TGC3F07P19 | 950 | healthy | - | P697 |
| 37 | 33dDN8TGC3F07P20 | 1050 | healthy | - |  |
| 38 | 33dDN8TGC3F07P21 | 200 | mummy | 0 |  |

**Table S7** Genotyping the GGTA1, CMAH and β4GalNT2 genes of cloned pigs with 8 gene editing by Sanger sequencing

| **Gene** | **No. of pigs** | **Sequence** | **Mutation** |
| --- | --- | --- | --- |
| GGTA1  on exon 1 | WT | TGATGTATTCCCAAAACACAACCATTACAGTTGAGACAAGCAGCATTGACAGAACCACTCTTCCTTTGACATTCATTATTTTCTCCTGGGAAAAGAAAAG |  |
|  | F01P02 | TGATGTATTCCCAAAACACAACCATTACAGTTGAGACAAGCAGCATTGACAGAACCACTCTTCCTTTGACAAC-------------CCTGGGAAAAGAAAAG | -13bp/+2bp 3/8 |
|  |  | TGATGTATTCCCAAAACACAACCATTACAGTTGAGACAAGCAGCAT-----------------------------------------------------G | -53bp 5/8 |
|  | F03P01 | TGATGTATTCCCAAAACACAACCATTACAGTTGAGACAAGCAGCATTGACAGAACCACTCTTCCTTTGACAAC-------------CCTGGGAAAAGAAAAG | -13bp/+2bp 6/9 |
|  |  | TGATGTATTCCCAAAACACAACCATTACAGTTGAGACAAGCAGCAT-----------------------------------------------------G | -53bp 3/9 |
|  | F03P02 | TGATGTATTCCCAAAACACAACCATTACAGTTGAGACAAGCAGCATTGACAGAACCACTCTTCCTTTGACAAC-------------CCTGGGAAAAGAAAAG | -13bp/+2bp 4/5 |
|  |  | TGATGTATTCCCAAAACACAACCATTACAGTTGAGACAAGCAGCAT-----------------------------------------------------G | -53bp 1/5 |
|  | F03P10 | TGATGTATTCCCAAAACACAACCATTACAGTTGAGACAAGCAGCATTGACAGAACCACTCTTCCTTTGACAAC-------------CCTGGGAAAAGAAAAG | -13bp/+2bp 3/10 |
|  |  | TGATGTATTCCCAAAACACAACCATTACAGTTGAGACAAGCAGCAT-----------------------------------------------------G | -53bp 7/10 |
|  | F07P03 | TGATGTATTCCCAAAACACAACCATTACAGTTGAGACAAGCAGCATTGACAGAACCACTCTTCCTTTGACAAC-------------CCTGGGAAAAGAAAAG | -13bp/+2bp 3/9 |
|  |  | TGATGTATTCCCAAAACACAACCATTACAGTTGAGACAAGCAGCAT-----------------------------------------------------G | -53bp 6/9 |
| CMAH | WT | TCACTGTCTTCCAACAGATCACGTACCTTACTCACG(84)TACTACACGAGCCTCCATCTGATTGG(62)GTAAGGAAGGGTGAGCCCTCAACTCCGAAGA |  |
|  | F01P02 | TCACTGTCTT--------------------------**(84)**-------------------TGATTGG(62)GTAAGGAAGGGTGAGCCCTCAACTCCGAAGA | -123bp 5/10 |
|  |  | TCACTGTCTTCCAACAGATCACGTACCTTACTCACG(84)TACTACACGAGCCTCCA---------**(62)**---------------------ACTCCGAAGA | -102bp 5/10 |
|  | F03P01 | TCACTGTCTT--------------------------**(84)**-------------------TGATTGG(62)GTAAGGAAGGGTGAGCCCTCAACTCCGAAGA | -123bp 6/9 |
|  |  | TCACTGTCTTCCAACAGATCACGTACCTTACTCACG(84)TACTACACGAGCCTCCA---------**(62)**---------------------ACTCCGAAGA | -102bp 3/9 |
|  | F03P02 | TCACTGTCTT--------------------------**(84)**-------------------TGATTGG(62)GTAAGGAAGGGTGAGCCCTCAACTCCGAAGA | -123bp 5/8 |
|  |  | TCACTGTCTTCCAACAGATCACGTACCTTACTCACG(84)TACTACACGAGCCTCCA---------**(62)**---------------------ACTCCGAAGA | -102bp 3/8 |
|  | F03P10 | TCACTGTCTT--------------------------**(84)**-------------------TGATTGG(62)GTAAGGAAGGGTGAGCCCTCAACTCCGAAGA | -123bp 2/4 |
|  |  | TCACTGTCTTCCAACAGATCACGTACCTTACTCACG(84)TACTACACGAGCCTCCA---------**(62)**---------------------ACTCCGAAGA | -102bp 7/4 |
|  | F07P03 | TCACTGTCTT--------------------------**(84)**-------------------TGATTGG(62)GTAAGGAAGGGTGAGCCCTCAACTCCGAAGA | -123bp 4/10 |
|  |  | TCACTGTCTTCCAACAGATCACGTACCTTACTCACG(84)TACTACACGAGCCTCCA---------**(62)**---------------------ACTCCGAAGA | -102bp 6/10 |
| β4GalNT2 | WT | AGCCCTAGATGTCTGTCGATCCTCAAGAT(99)CACCCTGGATGCGCAGACGCTGAAGCTTCTACC(15)CTACGGTGAAAACGGGTGAGATGGCAA |  |
|  |  | AGCCCTAGATGTCTGTCGATCCTCAAGAT(99)CACCCTGGATGCGCAGACGCTGAAGCTTCTACC(27)CTACAGTGGAATCTGGTGAGATGGCAA |  |
|  | F01P02 | AGCCCTAGATG--------------------**(99)**--------------------------------**(15)**-----------------AGATGGCAA | -183bp 2/9 |
|  |  | AGCCCTAGATG------------------**(99)**----------------------AAGCTTCTACC(27)CTACAGTGGAATCTGGTGAGATGGCAA | -139bp 7/9 |
|  | F03P01 | AGCCCTAGATG--------------------**(99)**--------------------------------**(15)**-----------------AGATGGCAA | -183bp 3/9 |
|  |  | AGCCCTAGATG------------------**(99)**----------------------AAGCTTCTACC(27)CTACAGTGGAATCTGGTGAGATGGCAA | -139bp 6/9 |
|  | F03P02 | AGCCCTAGATG--------------------**(99)**--------------------------------**(15)**-----------------AGATGGCAA | -183bp 3/5 |
|  |  | AGCCCTAGATG------------------**(99)**----------------------AAGCTTCTACC(27)CTACAGTGGAATCTGGTGAGATGGCAA | -139bp 2/5 |
|  | F03P10 | AGCCCTAGATG--------------------**(99)**--------------------------------**(15)**-----------------AGATGGCAA | -183bp 2/7 |
|  |  | AGCCCTAGATG------------------**(99)**----------------------AAGCTTCTACC(27)CTACAGTGGAATCTGGTGAGATGGCAA | -139bp 5/7 |
|  | F07P03 | AGCCCTAGATG--------------------**(99)**--------------------------------**(15)**-----------------AGATGGCAA | -183bp 5/16 |
|  |  | AGCCCTAGATG------------------**(99)**----------------------AAGCTTCTACC(27)CTACAGTGGAATCTGGTGAGATGGCAA | -139bp 11/16 |

Note：F01P02, 8TGC3F01P02; F03P01, 8TGC3F03P01; F03P02, 8TGC3F03P02; F03P10, 8TGC3F03P10; F07P03, 8TGC3F07P03

**Table S8** The generation of cloned pigs with 8 gene editing

| No. | ID | Body weight (g) | Status at birth | Survival time (day) | Surrogate mother |
| --- | --- | --- | --- | --- | --- |
| 1 | DN-8TG-C3P01 | 600 | healthy | 448 | P668 |
| 2 | DN-8TG-C3P02 | 600 | stillbirth | 0 |  |
| 3 | DN-8TG-C3P03 | 850 | healthy | - |  |
| 4 | DN-8TG-C3P04 | 550 | stillbirth | 0 |  |
| 5 | DN-8TG-C3P05 | 550 | healthy | 5 | P679 |
| 6 | DN-8TG-C3P06 | 600 | stillbirth | 0 |  |
| 7 | DN-8TG-C3P07 | - | mummy | 0 | P585 |
| 8 | DN-8TG-C3P08 | 750 | healthy | 5 |  |
| 9 | DN-8TG-C3P09 | 700 | healthy | 479 |  |
| 10 | DN-8TG-C3P10 | 800 | healthy | 4 |  |
| 11 | DN-8TG-C3P11 | 1010 | healthy | - |  |
| 12 | DN-8TG-C3P12 | 900 | stillbirth | 0 |  |
| 13 | DN-8TG-C3P13 | 800 | healthy | - |  |

**Table S9** Genotyping the GGTA1, CMAH and β4GalNT2 genes of cloned pigs with 8 gene editing by Sanger sequencing

| **Gene** | **No. of pigs** | **Sequence** | **Mutation** |
| --- | --- | --- | --- |
| GGTA1  on exon 1 | WT | TGATGTATTCCCAAAACACAACCATTACAGTTGAGACAAGCAGCATTGACAGAACCACTCTTCCTTTGACATTCATTATTTTCTCCTGGGAAAAGAAAAG |  |
|  | P01 | TGATGTATTCCCAAAACACAACCATTACAGTTGAGACAAGCAGCATTGACAGAACCACTCTTCCTTTGACAAC-------------CCTGGGAAAAGAAAAG | -13bp/+2bp 3/6 |
|  |  | TGATGTATTCCCAAAACACAACCATTACAGTTGAGACAAGCAGCAT-----------------------------------------------------G | -53bp 3/6 |
|  | P02 | TGATGTATTCCCAAAACACAACCATTACAGTTGAGACAAGCAGCATTGACAGAACCACTCTTCCTTTGACAAC-------------CCTGGGAAAAGAAAAG | -13bp/+2bp 2/5 |
|  |  | TGATGTATTCCCAAAACACAACCATTACAGTTGAGACAAGCAGCAT-----------------------------------------------------G | -53bp 3/5 |
|  | P03 | TGATGTATTCCCAAAACACAACCATTACAGTTGAGACAAGCAGCATTGACAGAACCACTCTTCCTTTGACAAC-------------CCTGGGAAAAGAAAAG | -13bp/+2bp 3/8 |
|  |  | TGATGTATTCCCAAAACACAACCATTACAGTTGAGACAAGCAGCAT-----------------------------------------------------G | -53bp 5/8 |
|  | P04 | TGATGTATTCCCAAAACACAACCATTACAGTTGAGACAAGCAGCATTGACAGAACCACTCTTCCTTTGACAAC-------------CCTGGGAAAAGAAAAG | -13bp/+2bp 3/7 |
|  |  | TGATGTATTCCCAAAACACAACCATTACAGTTGAGACAAGCAGCAT-----------------------------------------------------G | -53bp 4/7 |
| CMAH | WT | TCACTGTCTTCCAACAGATCACGTACCTTACTCACG(84)TACTACACGAGCCTCCATCTGATTGG(62)GTAAGGAAGGGTGAGCCCTCAACTCCGAAGA |  |
|  | P01 | TCACTGTCTT--------------------------**(84)**-------------------TGATTGG(62)GTAAGGAAGGGTGAGCCCTCAACTCCGAAGA | -123bp 6/15 |
|  |  | TCACTGTCTTCCAACAGATCACGTACCTTACTCACG(84)TACTACACGAGCCTCCA---------**(62)**---------------------ACTCCGAAGA | -102bp 9/15 |
|  | P02 | TCACTGTCTT--------------------------**(84)**-------------------TGATTGG(62)GTAAGGAAGGGTGAGCCCTCAACTCCGAAGA | -123bp 3/8 |
|  |  | TCACTGTCTTCCAACAGATCACGTACCTTACTCACG(84)TACTACACGAGCCTCCA---------**(62)**---------------------ACTCCGAAGA | -102bp 5/8 |
|  | P03 | TCACTGTCTT--------------------------**(84)**-------------------TGATTGG(62)GTAAGGAAGGGTGAGCCCTCAACTCCGAAGA | -123bp 1/6 |
|  |  | TCACTGTCTTCCAACAGATCACGTACCTTACTCACG(84)TACTACACGAGCCTCCA---------**(62)**---------------------ACTCCGAAGA | -102bp 5/6 |
|  | P04 | TCACTGTCTT--------------------------**(84)**-------------------TGATTGG(62)GTAAGGAAGGGTGAGCCCTCAACTCCGAAGA | -123bp 3/7 |
|  |  | TCACTGTCTTCCAACAGATCACGTACCTTACTCACG(84)TACTACACGAGCCTCCA---------**(62)**---------------------ACTCCGAAGA | -102bp 4/7 |
| β4GalNT2 | WT | AGCCCTAGATGTCTGTCGATCCTCAAGAT(99)CACCCTGGATGCGCAGACGCTGAAGCTTCTACC(15)CTACGGTGAAAACGGGTGAGATGGCAA |  |
|  |  | AGCCCTAGATGTCTGTCGATCCTCAAGAT(99)CACCCTGGATGCGCAGACGCTGAAGCTTCTACC(27)CTACAGTGGAATCTGGTGAGATGGCAA |  |
|  | P01 | AGCCCTAGATG--------------------**(99)**--------------------------------**(15)**-----------------AGATGGCAA | -183bp 2/4 |
|  |  | AGCCCTAGATG------------------**(99)**----------------------AAGCTTCTACC(27)CTACAGTGGAATCTGGTGAGATGGCAA | -139bp 2/4 |
|  | P02 | AGCCCTAGATG--------------------**(99)**--------------------------------**(15)**-----------------AGATGGCAA | -183bp 4/13 |
|  |  | AGCCCTAGATG------------------**(99)**----------------------AAGCTTCTACC(27)CTACAGTGGAATCTGGTGAGATGGCAA | -139bp 9/13 |
|  | P03 | AGCCCTAGATG--------------------**(99)**--------------------------------**(15)**-----------------AGATGGCAA | -183bp 12/18 |
|  |  | AGCCCTAGATG------------------**(99)**----------------------AAGCTTCTACC(27)CTACAGTGGAATCTGGTGAGATGGCAA | -139bp 6/18 |
|  | P04 | AGCCCTAGATG--------------------**(99)**--------------------------------**(15)**-----------------AGATGGCAA | -183bp 3/9 |
|  |  | AGCCCTAGATG------------------**(99)**----------------------AAGCTTCTACC(27)CTACAGTGGAATCTGGTGAGATGGCAA | -139bp 6/9 |

Note：P01, 8TGC3P01; P02, 8TGC3P02; P03, 8TGC3P03; P04, 8TGC3P04

**Table S10** Genotyping the GGTA1 gene of colonies Sanger sequencing in the second gene editing

| **Gene** | **No. of colonies** | **Sequence** | **Mutation** |
| --- | --- | --- | --- |
| GGTA1  on exon 8 | WT | ATGATGCGCATGAAGACCATCGGGGAGCACATCCTGGC(90)GGTGGCTCAGCTACAGGCCTGGTGGTACAAGGCACATCCTGACGAG |  |
|  | C1# | ATGATGCGCATGAAGACCATCGGGGAGCACATCCTGGC(90)GGTGGCTCAGCTACAGGCCTGGTGGTACAAGGCACATCCTGACGAG | WT 3/10 |
|  |  | ATGATGCGCATCA---------------------------**(90)**----------------------------------------TGACGAG | +2bp-156bp 6/10 |
|  |  | ATGATGCGCATGAAGACCATCGGGGAGCACATCCTGGC(90)GGTGGCTCAGCTACAGGCCTGGTGG-ACAAGGCACATCCTGACGAG | -1bp 1/10 |
|  | C3# | ----------------------GTGGAGACCCTGGGCCAGTCGGTGGCTCAGCTACAGGCCTGGTGG-------CACATCCTGACGAG | -469bp/-7bp 12/17 |
|  |  | ATGAT-----------------------------------**(90)**-------------------------------GCACATCCTGACGAG | -154bp 5/17 |
|  | C4# | ATGATGCGCATGAAGACCATCGGGGAGCACATCCTGGC(90)GGTGGCTCAGCTACAGGCCTGGTGGTACAAGGCACATCCTGACGAG | WT 4/8 |
|  |  | ATGATGCGCATGAAGACCAT------------------**(90)**--------------------------ACAAGGCACATCCTGACGAG | -134bp 3/8 |
|  |  | ATGATGCGCATGAAGACCATCGGGGAGCACATCCTGGC(90)GGTGGCTCAGCTACAGGCCTGGTGGTCTACAAGGCACATCCTGACGAG | +2bp 1/8 |
|  | C6# | ATGATGCGCATGAAGACCATCGGGGAGCACATCCTGGC(90)GGTGGCTCAGCTACAGGCCTGGTGGTACAAGGCACATCCTGACGAG | WT 1/7 |
|  |  | ATGATGCGCATGAAGACCAT------------------(90)--------------------------ACAAGGCACATCCTGACGAG | -134bp 4/7 |
|  |  | ATGATGCGCATGAAGACCATCGGGGAGCACATCCTGGC**(90)**GGTGGCTCAGCTACAGGCCTGGTGGTTACAAGGCACATCCTGACGAG | +1bp 2/7 |
|  | C7# | ATGATGCGCATGAAGACCATCGGGGAGCACATCCTGGC(90)GGTGGCTCAGCTACAGGCCTGGTGGTACAAGGCACATCCTGACGAG | WT 3/6 |
|  |  | ATGATGCGCATGAAGACCAT------------------**(90)**--------------------------ACAAGGCACATCCTGACGAG | -134bp 2/6 |
|  |  | ATGATGCGCATGAAGACCATCGGGGAGCACATCCTGGC**(90)**GGTGGCTCAGCTACAGGCCTGGTGGTTACAAGGCACATCCTGACGAG | +1bp 1/6 |
|  |  | ATGATGCGCATGAAG-----------------------**(90)**-------------------------TACAAGGCACATCCTGACGAG | -138bp 12/16 |
|  | C12# | ATGATGCGCATGAAGACCAT------------------**(90)**--------------------------ACAAGGCACATCCTGACGAG | -134bp 1/16 |
|  |  | ATGATGCGC---------------------ATCCTGGC(90)GGTGGCTCAGCTACAGGCCTGGTGGTATACAAGGCACATCCTGACGAG | +2bp/-21bp 3/16 |
|  | C15# | ATGATGCGCATGAAGACCAT------------------**(90)**--------------------------ACAAGGCACATCCTGACGAG | -134bp 6/11 |
|  |  | ATGATGCGCATGAAGACCAATCGGGGAGCACATCCTGGC(90)GGTGGCTCAGCTACAGGCCTGGTGGTTACAAGGCACATCCTGACGAG | +2bp 5/11 |
|  | C16# | ATGATGCGCATGAAGACCATCGGGGAGCACATCCTGGC(90)GGTGGCTCAGCTACAGGCCTGGTGGTACAAGGCACATCCTGACGAG | WT 2/8 |
|  |  | ATGATGCGCATCA-------------------------**(90)**---------------------------------------TGACGAG | -154bp 1/8 |
|  |  | ATGATGCGCATGAAGACCA----------CATCCTGGC(90)GGTGGCTCAGCTACAGGCC---------------CATCCTGACGAG | -25bp 4/8 |
|  |  | ATGATGCGCATGAAGACCAATCGGGGAGCACATCCTGGC(90)GGTGGCTCAGCTACAGGCCTGGTGGTTACAAGGCACATCCTGACGAG | +2bp 1/8 |
|  | C19# | ATGATGCGCATGAAGACCATCGGGGAGCACATCCTGGC(90)GGTGGCTCAGCTACAGGCCTGGTGGTACAAGGCACATCCTGACGAG | WT 1/9 |
|  |  | ATGATGCGCATGAAGACCAT------------------**(90)**--------------------------ACAAGGCACATCCTGACGAG | -134bp 8/9 |
|  | C22# | ATGATGCGCATGAAGACCATCGGGGAGCACATCCTGGC(90)GGTGGCTCAGCTACAGGCCTGGTGGTACAAGGCACATCCTGACGAG | WT 5/9 |
|  |  | ATGATGCGCATGAAGACCA-------------------**(90)**---------------------------CAAGGCACATCCTGACGAG | -136bp 4/9 |
|  | C23# | ATGATGCGCATGAAGACCATCGGGGAGCACATCCTGGC(90)GGTGGCTCAGCTACAGGCCTGGTGGTACAAGGCACATCCTGACGAG | WT 1/9 |
|  |  | -----------------------------CATCCTGGC(90)GGTGGCTCAGCTACAGGCCTGGTGGTACAAGGCACATCCTGACGAG | -347bp 4/6 |
|  |  | ATGATGCGCATGAAGACC--------------------**(90)**-------------------------TACAAGGCACATCCTGACGAG | -135bp 1/6 |
|  | C28# | ATGATGCGCATGAAGACCATCGGGGAGCACATCCTGGC(90)GGTGGCTCAGCTACAGGCCTGGTGGTACAAGGCACATCCTGACGAG | WT 1/9 |
|  |  | ATGATGCGCATGAAGACC--------------------**(90)**-------------------------TACAAGGCACATCCTGACGAG | -135bp 3/6 |
|  |  | ATGATGCGCATGAAGAC---------------------**(90)**---------------------------------ACATCCTGACGAG | -144bp 2/6 |
|  | C31# | ATGATGCGCATGAAGACCATCGGGGAGCACATCCTGGC(90)GGTGGCTCAGCTACAGGCCTGGTGGTACAAGGCACATCCTGACGAG | WT 5/7 |
|  |  | ATGATGCGCATGAAGACC--------------------**(90)**-------------------------TACAAGGCACATCCTGACGAG | -135bp 2/7 |
|  | C36# | ATGATGCGCATGAAGACCATCGGGGAGCACATCCTGGC(90)GGTGGCTCAGCTACAGGCCTGGTGGTACAAGGCACATCCTGACGAG | WT 3/4 |
|  |  | ATGATGCGCATGAAGACC--------------------**(90)**-------------------------TACAAGGCACATCCTGACGAG | -135bp 1/4 |
|  | C54# | ATGATGCGCATGAAGACCATCGGGGAGCACATCCTGGC(90)GGTGGCTCAGCTACAGGCCTGGTGGTACAAGGCACATCCTGACGAG | WT 1/8 |
|  |  | ATGATGCGCATGAAGAC--TCGGGGAGCACATCCTGGC(90)GGTGGCTCAGCTACAGGCCTGGTGGTTACAAGGCACATCCTGACGAG | -135bp 1/8 |
|  |  | ATGATGCGCATGAAGACCAT------------------**(90)**--------------------------ACAAGGCACATCCTGACGAG | WT 2/8 |
|  |  | ATGATGCGCATGA-------------------------(90)-------------------------------GCACATCCTGACGAG | -135bp 1/8 |
|  |  | ATGATGCGCAT---------------------------**(90)**---------------------------------------------- | WT 3/8 |
|  | C58# | --------------------------------------**(90)**----------------------------------CATCCTGACGAG  GTAGCTGAGCCACCGACTGGCCCAGGGTCTCCACCCCAAAGTTGTTTTGGAAGACCTGATCCACGTCCATGCAGAAGAGGAAGTCCACCTCGTCTTCATGCGCAT (inserted sequence) | -440bp/+105bp 2/6 |
|  |  | ATGATGCGCATGAA------CGGGGAGCACATCCTGGC**(90)**GGTGGCTCA---------------------------TCCTGACGAG | -135bp 4/6 |

**Table S11** Genotyping the GGTA1, CMAH and β4GalNT2 genes of cloned fetuses with 10 gene editing by Sanger sequencing

| **Gene** | **No. of pigs** | **Sequence** | **Mutation** |
| --- | --- | --- | --- |
| GGTA1  on exon 3 | WT | TGATGTATTCCCAAAACACAACCATTACAGTTGAGACAAGCAGCATTGACAGAACCACTCTTCCTTTGACATTCATTATTTTCTCCTGGGAAAAGAAAAG |  |
|  | C3F01 | TGATGTATTCCCAAAACACAACCATTACAGTTGAGACAAGCAGCATTGACAGAACCACTCTTCCTTTGACAAC-------------CCTGGGAAAAGAAAAG | -13bp/+2bp 5/8 |
|  |  | TGATGTATTCCCAAAACACAACCATTACAGTTGAGACAAGCAGCAT-----------------------------------------------------G | -53bp 3/8 |
|  | C3F02 | TGATGTATTCCCAAAACACAACCATTACAGTTGAGACAAGCAGCATTGACAGAACCACTCTTCCTTTGACAAC-------------CCTGGGAAAAGAAAAG | -13bp/+2bp 1/9 |
|  |  | TGATGTATTCCCAAAACACAACCATTACAGTTGAGACAAGCAGCAT-----------------------------------------------------G | -53bp 8/9 |
|  | C3F03 | TGATGTATTCCCAAAACACAACCATTACAGTTGAGACAAGCAGCATTGACAGAACCACTCTTCCTTTGACAAC-------------CCTGGGAAAAGAAAAG | -13bp/+2bp 4/10 |
|  |  | TGATGTATTCCCAAAACACAACCATTACAGTTGAGACAAGCAGCAT-----------------------------------------------------G | -53bp 6/10 |
|  | C3F04 | TGATGTATTCCCAAAACACAACCATTACAGTTGAGACAAGCAGCATTGACAGAACCACTCTTCCTTTGACAAC-------------CCTGGGAAAAGAAAAG | -13bp/+2bp 2/9 |
|  |  | TGATGTATTCCCAAAACACAACCATTACAGTTGAGACAAGCAGCAT-----------------------------------------------------G | -53bp 7/9 |
|  | C3F05 | TGATGTATTCCCAAAACACAACCATTACAGTTGAGACAAGCAGCATTGACAGAACCACTCTTCCTTTGACAAC-------------CCTGGGAAAAGAAAAG | -13bp/+2bp 4/8 |
|  |  | TGATGTATTCCCAAAACACAACCATTACAGTTGAGACAAGCAGCAT-----------------------------------------------------G | -53bp 4/8 |
| GGTA1  on exon 8 | WT | ATGATGCGCATGAAGACCATCGGGGAGCACATCCTGGC(90)GGTGGCTCAGCTACAGGCCTGGTGGTACAAGGCACATCCTGACGAG |  |
|  | C3F01 | -----------------------GTGGAGACCCTGGGCCAGTCGGTGGCTCAGCTACAGGCCTGGTGG-------CACATCCTGACGAG | -469/-7bp 2/5 |
|  |  | ATGAT--------------------------------------**(90)**------------------------------------GCACATCCTGACGAG | -154bp 3/5 |
|  | C3F02 | -----------------------GTGGAGACCCTGGGCCAGTCGGTGGCTCAGCTACAGGCCTGGTGG-------CACATCCTGACGAG | -469/-7bp 5/7 |
|  |  | ATGAT--------------------------------------**(90)**------------------------------------GCACATCCTGACGAG | -154bp 2/7 |
|  | C3F03 | -----------------------GTGGAGACCCTGGGCCAGTCGGTGGCTCAGCTACAGGCCTGGTGG-------CACATCCTGACGAG | -469/-7bp 2/5 |
|  |  | ATGAT--------------------------------------**(90)**------------------------------------GCACATCCTGACGAG | -154bp 5/6 |
|  | C3F04 | -----------------------GTGGAGACCCTGGGCCAGTCGGTGGCTCAGCTACAGGCCTGGTGG-------CACATCCTGACGAG | -469/-7bp 9/9 |
|  | C3F05 | -----------------------GTGGAGACCCTGGGCCAGTCGGTGGCTCAGCTACAGGCCTGGTGG-------CACATCCTGACGAG | -469/-7bp 5/9 |
|  |  | ATGAT--------------------------------------**(90)**------------------------------------GCACATCCTGACGAG | -154bp 4/9 |
| CMAH | WT | TCACTGTCTTCCAACAGATCACGTACCTTACTCACG(84)TACTACACGAGCCTCCATCTGATTGG(62)GTAAGGAAGGGTGAGCCCTCAACTCCGAAGA |  |
|  | C3F01 | TCACTGTCTT--------------------------**(84)**-------------------TGATTGG(62)GTAAGGAAGGGTGAGCCCTCAACTCCGAAGA | -123bp 6/10 |
|  |  | TCACTGTCTTCCAACA----------CTTACTCACG(84)TACTACACGAGCCTCCA---------**(62)**---------------------ACTCCGAAGA | -102bp 4/10 |
|  | C3F02 | TCACTGTCTT--------------------------**(84)**-------------------TGATTGG(62)GTAAGGAAGGGTGAGCCCTCAACTCCGAAGA | -123bp 7/11 |
|  |  | TCACTGTCTTCCAACA----------CTTACTCACG(84)TACTACACGAGCCTCCA---------**(62)**---------------------ACTCCGAAGA | -102bp 4/11 |
|  | C3F03 | TCACTGTCTT--------------------------**(84)**-------------------TGATTGG(62)GTAAGGAAGGGTGAGCCCTCAACTCCGAAGA | -123bp 5/10 |
|  |  | TCACTGTCTTCCAACA----------CTTACTCACG(84)TACTACACGAGCCTCCA---------**(62)**---------------------ACTCCGAAGA | -102bp 5/10 |
|  | C3F04 | TCACTGTCTT--------------------------**(84)**-------------------TGATTGG(62)GTAAGGAAGGGTGAGCCCTCAACTCCGAAGA | -123bp 4/10 |
|  |  | TCACTGTCTTCCAACA----------CTTACTCACG(84)TACTACACGAGCCTCCA---------**(62)**---------------------ACTCCGAAGA | -102bp 6/10 |
|  | C3F05 | TCACTGTCTT--------------------------**(84)**-------------------TGATTGG(62)GTAAGGAAGGGTGAGCCCTCAACTCCGAAGA | -123bp 6/10 |
|  |  | TCACTGTCTTCCAACA----------CTTACTCACG(84)TACTACACGAGCCTCCA---------**(62)**---------------------ACTCCGAAGA | -102bp 4/10 |
| β4GalNT2 | WT | AGCCCTAGATGTCTGTCGATCCTCAAGAT(99)CACCCTGGATGCGCAGACGCTGAAGCTTCTACC(15)CTACGGTGAAAACGGGTGAGATGGCAA |  |
|  |  | AGCCCTAGATGTCTGTCGATCCTCAAGAT(99)CACCCTGGATGCGCAGACGCTGAAGCTTCTACC(27)CTACAGTGGAATCTGGTGAGATGGCAA |  |
|  | C3F01 | AGCCCTAGATG--------------------**(99)**--------------------------------**(15)**-----------------AGATGGCAA | -183bp 5/9 |
|  |  | AGCCCTAGATG------------------**(99)**----------------------AAGCTTCTACC(27)CTACAGTGGAATCTGGTGAGATGGCAA | -139bp 4/9 |
|  | C3F02 | AGCCCTAGATG--------------------**(99)**--------------------------------**(15)**-----------------AGATGGCAA | -183bp 6/10 |
|  |  | AGCCCTAGATG------------------**(99)**----------------------AAGCTTCTACC(27)CTACAGTGGAATCTGGTGAGATGGCAA | -139bp 4/10 |
|  | C3F03 | AGCCCTAGATG--------------------**(99)**--------------------------------**(15)**-----------------AGATGGCAA | -183bp 4/10 |
|  |  | AGCCCTAGATG------------------**(99)**----------------------AAGCTTCTACC(27)CTACAGTGGAATCTGGTGAGATGGCAA | -139bp 6/10 |
|  | C3F04 | AGCCCTAGATG--------------------**(99)**--------------------------------**(15)**-----------------AGATGGCAA | -183bp 5/10 |
|  |  | AGCCCTAGATG------------------**(99)**----------------------AAGCTTCTACC(27)CTACAGTGGAATCTGGTGAGATGGCAA | -139bp 5/10 |
|  | C3F05 | AGCCCTAGATG--------------------**(99)**--------------------------------**(15)**-----------------AGATGGCAA | -183bp 3/10 |
|  |  | AGCCCTAGATG------------------**(99)**----------------------AAGCTTCTACC(27)CTACAGTGGAATCTGGTGAGATGGCAA | -139bp 8/10 |

**Table S13** Genotyping the GGTA1 gene of cloned pigs with 10 gene editing by Sanger sequencing

| **Gene** | **No. of pigs** | **Sequence** | **Mutation** |
| --- | --- | --- | --- |
| GGTA1  on exon 3 | WT | ATGATGCGCATGAAGACCATCGGGGAGCACATCCTGGC(90)GGTGGCTCAGCTACAGGCCTGGTGGTACAAGGCACATCCTGACGAG |  |
|  | F01P05 | -----------------------GTGGAGACCCTGGGCCAGTCGGTGGCTCAGCTACAGGCCTGGTGG-------CACATCCTGACGAG | -469/-7bp 3/6 |
|  |  | ATGAT--------------------------------------**(90)**------------------------------------GCACATCCTGACGAG | -154bp 3/6 |
|  | F01P06 | -----------------------GTGGAGACCCTGGGCCAGTCGGTGGCTCAGCTACAGGCCTGGTGG-------CACATCCTGACGAG | -469/-7bp 6/8 |
|  |  | ATGAT--------------------------------------**(90)**------------------------------------GCACATCCTGACGAG | -154bp 2/8 |
|  | F01P16 | -----------------------GTGGAGACCCTGGGCCAGTCGGTGGCTCAGCTACAGGCCTGGTGG-------CACATCCTGACGAG | -469/-7bp 5/6 |
|  |  | ATGAT--------------------------------------**(90)**------------------------------------GCACATCCTGACGAG | -154bp 1/6 |
|  | F03P11 | -----------------------GTGGAGACCCTGGGCCAGTCGGTGGCTCAGCTACAGGCCTGGTGG-------CACATCCTGACGAG | -469/-7bp 7/7 |
|  | F04P01 | -----------------------GTGGAGACCCTGGGCCAGTCGGTGGCTCAGCTACAGGCCTGGTGG-------CACATCCTGACGAG | -469/-7bp 5/8 |
|  |  | ATGAT--------------------------------------**(90)**------------------------------------GCACATCCTGACGAG | -154bp 3/8 |
|  | F04P02 | -----------------------GTGGAGACCCTGGGCCAGTCGGTGGCTCAGCTACAGGCCTGGTGG-------CACATCCTGACGAG | -469/-7bp 4/4 |
|  | F04P13 | -----------------------GTGGAGACCCTGGGCCAGTCGGTGGCTCAGCTACAGGCCTGGTGG-------CACATCCTGACGAG | -469/-7bp 7/7 |

Note: F01P05, 10TGC3F01P05; F01P06, 10TGC3F01P06; P04, 10TGC3F01P16; F03P11, 10TGC3F03P11; F04P01, 10TGC3F04P01; P07, 10TGC3F04P02; F04P13, 10TGC3F04P13

**Table S14 The F1 generation of 10 GEC male (♂) pigs naturally mated with 8 GEC female (♀) pigs**

| No. | ID | Parents | Body weight (g) | Status at birth | Survival time (day) | Gender |
| --- | --- | --- | --- | --- | --- | --- |
| 1 | G1F1P01 | DN10TGC3F02P02×  DN8TGC12F01P17 | 550 | weak | 2 | ♂ |
| 2 | G1F1P02 |  | 580 | healthy | 58 | ♀ |
| 3 | G1F1P03 |  | 510 | weak | 3 | ♀ |
| 4 | G1F1P04 | DN10TGC3F02P02  or  DN10TGC3F03P07  ×  DN8TGC12F01P08 | 260 | weak | 1 | ♂ |
| 5 | G1F1P05 |  | 430 | healthy | - | ♂ |
| 6 | G1F1P06 |  | 320 | stillbirth | 0 | ♂ |
| 7 | G1F1P07 |  | 290 | weak | 56 | ♂ |
| 8 | G1F1P08 |  | 400 | healthy | - | ♀ |
| 9 | G1F1P09 |  | 420 | healthy | - | ♂ |
| 10 | G1F1P10 |  | 380 | stillbirth | 0 | ♂ |
| 11 | G1F1P11 |  | 410 | healthy | - | ♀ |
| 12 | G1F1P12 | DN10TGC3F01P04  ×  DN8TGC12F01P02 | 630 | healthy | - | ♀ |
| 13 | G1F1P13 |  | 610 | healthy | 11 | ♀ |
| 14 | G1F1P14 |  | 550 | healthy | - | ♀ |
| 15 | G1F1P15 |  | 550 | healthy | - | ♂ |
| 16 | G1F1P16 |  | 480 | healthy | - | ♀ |
| 17 | G1F1P17 |  | 640 | healthy | - | ♂ |
| 18 | G1F1P18 |  | 400 | healthy | - | ♂ |
| 19 | G1F1P19 |  | 350 | healthy | 11 | ♂ |
| 20 | G1F1P20 | DN10TGC3F02P02  or  DN10TGC3F04P12  ×  DN8TGC12F01P25 | 550 | healthy | - | ♀ |
| 21 | G1F1P21 |  | 380 | healthy | - | ♂ |
| 22 | G1F1P22 |  | 320 | stillbirth | 0 | ♂ |
| 23 | G1F1P23 |  | 540 | healthy | - | ♀ |
| 24 | G1F1P24 |  | 230 | stillbirth | 0 | ♂ |

**Table S15 The genotype of F1 generation**

| **Gene** | **No. of pigs** | **Sequence** | **Mutation** |
| --- | --- | --- | --- |
| GGTA1  on exon 3 | WT | TGATGTATTCCCAAAACACAACCATTACAGTTGAGACAAGCAGCATTGACAGAACCACTCTTCCTTTGACATTCATTATTTTCTCCTGGGAAAAGAAAAG | WT |
|  | G1F1P01 | TGATGTATTCCCAAAA-----------------------------------------------------ACATTCATTATTTTCTCCTGGGAAAAGAAAAG | -52bp 3/10 |
|  |  | TGATGTATTCCCAAAACACAACCATTACAGTTGAGACAAGCAGCAT-----------------------------------------------------G | -53bp 7/10 |
|  | G1F1P02 | TGATGTATTCCCAAAA-----------------------------------------------------ACATTCATTATTTTCTCCTGGGAAAAGAAAAG | -52bp 5/10 |
|  |  | TGATGTATTCCCAAAACACAACCATTACAGTTGAGACAAGCAGCATTGACAGAACCACTCTTCCTTTGACAAC-------------CCTGGGAAAAGAAAAG | +2/-13bp 5/10 |
|  | G1F1P03 | TGATGTATTCCCAAAA-----------------------------------------------------ACATTCATTATTTTCTCCTGGGAAAAGAAAAG | -52bp 4/9 |
|  |  | TGATGTATTCCCAAAACACAACCATTACAGTTGAGACAAGCAGCATTGACAGAACCACTCTTCCTTTGACAAC-------------CCTGGGAAAAGAAAAG | +2/-13bp 5/9 |
|  | G1F1P04 | TGATGTATTCCCAAAA-----------------------------------------------------ACATTCATTATTTTCTCCTGGGAAAAGAAAAG | -52bp 2/9 |
|  |  | TGATGTATTCCCAAAACACAACCATTACAGTTGAGACAAGCAGCATTGACAGAACCACTCTTCCTTTGACAAC-------------CCTGGGAAAAGAAAAG | +2/-13bp 7/9 |
|  | G1F1P05 | TGATGTATTCCCAAAA-----------------------------------------------------ACATTCATTATTTTCTCCTGGGAAAAGAAAAG | -52bp 4/9 |
|  |  | TGATGTATTCCCAAAACACAACCATTACAGTTGAGACAAGCAGCATTGACAGAACCACTCTTCCTTTGACAAC-------------CCTGGGAAAAGAAAAG | +2/-13bp 5/9 |
|  | G1F1P06 | TGATGTATTCCCAAAA-----------------------------------------------------ACATTCATTATTTTCTCCTGGGAAAAGAAAAG | -52bp 5/9 |
|  |  | TGATGTATTCCCAAAACACAACCATTACAGTTGAGACAAGCAGCATTGACAGAACCACTCTTCCTTTGACAAC-------------CCTGGGAAAAGAAAAG | +2/-13bp 4/9 |
|  | G1F1P07 | TGATGTATTCCCAAAA-----------------------------------------------------ACATTCATTATTTTCTCCTGGGAAAAGAAAAG | -52bp 5/10 |
|  |  | TGATGTATTCCCAAAACACAACCATTACAGTTGAGACAAGCAGCATTGACAGAACCACTCTTCCTTTGACAAC-------------CCTGGGAAAAGAAAAG | +2/-13bp 5/10 |
|  | G1F1P08 | TGATGTATTCCCAAAA-----------------------------------------------------ACATTCATTATTTTCTCCTGGGAAAAGAAAAG | -52bp 2/10 |
|  |  | TGATGTATTCCCAAAACACAACCATTACAGTTGAGACAAGCAGCAT-----------------------------------------------------G | -53bp 8/10 |
|  | G1F1P09 | TGATGTATTCCCAAAA-----------------------------------------------------ACATTCATTATTTTCTCCTGGGAAAAGAAAAG | -52bp 3/9 |
|  |  | TGATGTATTCCCAAAACACAACCATTACAGTTGAGACAAGCAGCAT-----------------------------------------------------G | -53bp 6/9 |
|  | G1F1P10 | TGATGTATTCCCAAAA-----------------------------------------------------ACATTCATTATTTTCTCCTGGGAAAAGAAAAG | -52bp 5/10 |
|  |  | TGATGTATTCCCAAAACACAACCATTACAGTTGAGACAAGCAGCATTGACAGAACCACTCTTCCTTTGACAAC-------------CCTGGGAAAAGAAAAG | +2/-13bp 5/10 |
|  | G1F1P11 | TGATGTATTCCCAAAA-----------------------------------------------------ACATTCATTATTTTCTCCTGGGAAAAGAAAAG | -52bp 7/10 |
|  |  | TGATGTATTCCCAAAACACAACCATTACAGTTGAGACAAGCAGCATTGACAGAACCACTCTTCCTTTGACAAC-------------CCTGGGAAAAGAAAAG | +2/-13bp 3/10 |
| GGTA1  on exon 8 | WT | ATGATGCGCATGAAGACCATCGGGGAGCACATCCTGGC(90)GGTGGCTCAGCTACAGGCCTGGTGGTACAAGGCACATCCTGACGAG | WT |
|  | G1F1P01 | ATGATGCGCATGAAGACCATCGGGGAGCACATCCTGGC(90)GGTGGCTCAGCTACAGGCCTGGTGGTACAAGGCACATCCTGACGAG | WT 1/7 |
|  |  | ATGAT--------------------------------------**(90)**------------------------------------GCACATCCTGACGAG | -154bp 6/7 |
|  | G1F1P02 | ATGATGCGCATGAAGACCATCGGGGAGCACATCCTGGC(90)GGTGGCTCAGCTACAGGCCTGGTGGTACAAGGCACATCCTGACGAG | WT 2/10 |
|  |  | -----------------------GTGGAGACCCTGGGCCAGTCGGTGGCTCAGCTACAGGCCTGGTGG-------CACATCCTGACGAG | -7/-469bp 8/10 |
|  | G1F1P03 | ATGATGCGCATGAAGACCATCGGGGAGCACATCCTGGC(90)GGTGGCTCAGCTACAGGCCTGGTGGTACAAGGCACATCCTGACGAG | WT 4/9 |
|  |  | -----------------------GTGGAGACCCTGGGCCAGTCGGTGGCTCAGCTACAGGCCTGGTGG-------CACATCCTGACGAG | -7/-469bp 5/9 |
|  | G1F1P04 | ATGATGCGCATGAAGACCATCGGGGAGCACATCCTGGC(90)GGTGGCTCAGCTACAGGCCTGGTGGTACAAGGCACATCCTGACGAG | WT 3/10 |
|  |  | -----------------------GTGGAGACCCTGGGCCAGTCGGTGGCTCAGCTACAGGCCTGGTGG-------CACATCCTGACGAG | -7/-469bp 7/10 |
|  | G1F1P05 | ATGATGCGCATGAAGACCATCGGGGAGCACATCCTGGC(90)GGTGGCTCAGCTACAGGCCTGGTGGTACAAGGCACATCCTGACGAG | WT 2/10 |
|  |  | -----------------------GTGGAGACCCTGGGCCAGTCGGTGGCTCAGCTACAGGCCTGGTGG-------CACATCCTGACGAG | -7/-469bp 8/10 |
|  | G1F1P06 | ATGATGCGCATGAAGACCATCGGGGAGCACATCCTGGC(90)GGTGGCTCAGCTACAGGCCTGGTGGTACAAGGCACATCCTGACGAG | WT 3/10 |
|  |  | -----------------------GTGGAGACCCTGGGCCAGTCGGTGGCTCAGCTACAGGCCTGGTGG-------CACATCCTGACGAG | -7/-469bp 7/10 |
|  | G1F1P07 | ATGATGCGCATGAAGACCATCGGGGAGCACATCCTGGC(90)GGTGGCTCAGCTACAGGCCTGGTGGTACAAGGCACATCCTGACGAG | WT 1/12 |
|  |  | -----------------------GTGGAGACCCTGGGCCAGTCGGTGGCTCAGCTACAGGCCTGGTGG-------CACATCCTGACGAG | -7/-469bp 11/12 |
|  | G1F1P08 | ATGATGCGCATGAAGACCATCGGGGAGCACATCCTGGC(90)GGTGGCTCAGCTACAGGCCTGGTGGTACAAGGCACATCCTGACGAG | WT 4/7 |
|  |  | -----------------------GTGGAGACCCTGGGCCAGTCGGTGGCTCAGCTACAGGCCTGGTGG-------CACATCCTGACGAG | -7/-469bp 3/7 |
|  | G1F1P09 | ATGATGCGCATGAAGACCATCGGGGAGCACATCCTGGC(90)GGTGGCTCAGCTACAGGCCTGGTGGTACAAGGCACATCCTGACGAG | WT 6/10 |
|  |  | -----------------------GTGGAGACCCTGGGCCAGTCGGTGGCTCAGCTACAGGCCTGGTGG-------CACATCCTGACGAG | -7/-469bp 4/10 |
|  | G1F1P10 | ATGATGCGCATGAAGACCATCGGGGAGCACATCCTGGC(90)GGTGGCTCAGCTACAGGCCTGGTGGTACAAGGCACATCCTGACGAG | WT 3/10 |
|  |  | -----------------------GTGGAGACCCTGGGCCAGTCGGTGGCTCAGCTACAGGCCTGGTGG-------CACATCCTGACGAG | -7/-469bp 7/10 |
|  | G1F1P11 | ATGATGCGCATGAAGACCATCGGGGAGCACATCCTGGC(90)GGTGGCTCAGCTACAGGCCTGGTGGTACAAGGCACATCCTGACGAG | WT 2/8 |
|  |  | -----------------------GTGGAGACCCTGGGCCAGTCGGTGGCTCAGCTACAGGCCTGGTGG-------CACATCCTGACGAG | -7/-469bp 6/8 |
| CMAH | WT | TCACTGTCTTCCAACAGATCACGTACCTTACTCACG(84)TACTACACGAGCCTCCATCTGATTGG(62)GTAAGGAAGGGTGAGCCCTCAACTCCGAAGA | WT |
|  | G1F1P01 | TCACTGTCTTCCAAC-GATCACGTACCTTACTCACG(84)TACTACACGAGCCTCCA---GATTGG(62)GTAAGGAAGGGTGAGCCCTCAACTCCGAAGA | -4 bp 4/8 |
|  |  | TCACTGTCTT--------------------------**(84)**-------------------TGATTGG(62)GTAAGGAAGGGTGAGCCCTCAACTCCGAAGA | -123bp 4/8 |
|  | G1F1P02 | TCACTGTCTTCCAAC-GATCACGTACCTTACTCACG(84)TACTACACGAGCCTCCA---GATTGG(62)GTAAGGAAGGGTGAGCCCTCAACTCCGAAGA | -4 bp 4/10 |
|  |  | TCACTGTCTT--------------------------**(84)**-------------------TGATTGG(62)GTAAGGAAGGGTGAGCCCTCAACTCCGAAGA | -123bp 6/10 |
|  | G1F1P03 | TCACTGTCTTCCAAC-GATCACGTACCTTACTCACG(84)TACTACACGAGCCTCCA---GATTGG(62)GTAAGGAAGGGTGAGCCCTCAACTCCGAAGA | -4 bp 4/10 |
|  |  | TCACTGTCTT--------------------------**(84)**-------------------TGATTGG(62)GTAAGGAAGGGTGAGCCCTCAACTCCGAAGA | -123bp 6/10 |
|  | G1F1P04 | TCACTGTCTTCCAAC-GATCACGTACCTTACTCACG(84)TACTACACGAGCCTCCA---GATTGG(62)GTAAGGAAGGGTGAGCCCTCAACTCCGAAGA | -4 bp 4/10 |
|  |  | TCACTGTCTT--------------------------**(84)**-------------------TGATTGG(62)GTAAGGAAGGGTGAGCCCTCAACTCCGAAGA | -123bp 6/10 |
|  | G1F1P05 | TCACTGTCTTCCAACAGAT-----------------**(84)**-----------------------TGG(62)GTAAGGAAGGGTGAGCCCTCAACTCCGAAGA | -124bp 4/10 |
|  |  | TCACTGTCTTCCAACA----------CTTACTCACG(84)TACTACACGAGCCTCCA---------**(62)**---------------------ACTCCGAAGA | -10/-92bp 6/10 |
|  | G1F1P06 | TCACTGTCTTCCAAC-GATCACGTACCTTACTCACG(84)TACTACACGAGCCTCCA---GATTGG(62)GTAAGGAAGGGTGAGCCCTCAACTCCGAAGA | -4 bp 4/8 |
|  |  | TCACTGTCTTCCAACA----------CTTACTCACG(84)TACTACACGAGCCTCCA---------**(62)**---------------------ACTCCGAAGA | -10/-92bp 4/8 |
|  | G1F1P07 | TCACTGTCTTCCAAC-GATCACGTACCTTACTCACG(84)TACTACACGAGCCTCCA---GATTGG(62)GTAAGGAAGGGTGAGCCCTCAACTCCGAAGA | -4 bp 2/10 |
|  |  | TCACTGTCTTCCAACA----------CTTACTCACG(84)TACTACACGAGCCTCCA---------**(62)**---------------------ACTCCGAAGA | -10/-92bp 8/10 |
|  | G1F1P08 | TCACTGTCTTCCAAC-GATCACGTACCTTACTCACG(84)TACTACACGAGCCTCCA---GATTGG(62)GTAAGGAAGGGTGAGCCCTCAACTCCGAAGA | -4 bp 4/9 |
|  |  | TCACTGTCTT--------------------------**(84)**-------------------TGATTGG(62)GTAAGGAAGGGTGAGCCCTCAACTCCGAAGA | -123bp 5/9 |
|  | G1F1P09 | TCACTGTCTTCCAAC-GATCACGTACCTTACTCACG(84)TACTACACGAGCCTCCA---GATTGG(62)GTAAGGAAGGGTGAGCCCTCAACTCCGAAGA | -4 bp 5/10 |
|  |  | TCACTGTCTTCCAACA----------CTTACTCACG(84)TACTACACGAGCCTCCA---------**(62)**---------------------ACTCCGAAGA | -10/-92bp 5/10 |
|  | G1F1P10 | TCACTGTCTTCCAACAGAT-----------------**(84)**-----------------------TGG(62)GTAAGGAAGGGTGAGCCCTCAACTCCGAAGA | -124bp 3/10 |
|  |  | TCACTGTCTTCCAACA----------CTTACTCACG(84)TACTACACGAGCCTCCA---------**(62)**---------------------ACTCCGAAGA | -10/-92bp 7/10 |
|  | F1P11 | TCACTGTCTTCCAAC-GATCACGTACCTTACTCACG(84)TACTACACGAGCCTCCA---GATTGG(62)GTAAGGAAGGGTGAGCCCTCAACTCCGAAGA | -4bp 5/9 |
|  |  | TCACTGTCTT--------------------------**(84)**-------------------TGATTGG(62)GTAAGGAAGGGTGAGCCCTCAACTCCGAAGA | -123bp 4/9 |
|  |  | TCACTGTCTTCCAACAGAT-----------------**(84)**-----------------------TGG(62)GTAAGGAAGGGTGAGCCCTCAACTCCGAAGA | -124bp 6/7 |
| β4GalNT2 | WT | AGCCCTAGATGTCTGTCGATCCTCAAGAT(99)CACCCTGGATGCGCAGACGCTGAAGCTTCTACC(15)CTACGGTGAAAACGGGTGAGATGGCAA | WT |
|  |  | AGCCCTAGATGTCTGTCGATCCTCAAGAT(99)CACCCTGGATGCGCAGACGCTGAAGCTTCTACC(27)CTACAGTGGAATCTGGTGAGATGGCAA | WT |
|  | G1F1P01 | AGCCCTAGAG------------------**(99)**--------ATGCGCAGACGCTGAAGCTTCTACC(27)CTACAGTGGAATCTGGTGAGATGGCAA | -125bp 4/16 |
|  |  | AGCCCTAGAT--------------AAGAT(99)CACCCTGGAT-----------------------(15)------------------AGATGGCAA | -14/-56bp 4/16 |
|  |  | AGCCCTAGATG--------------------**(99)**--------------------------------**(15)**-----------------AGATGGCAA | -183bp 1/16 |
|  |  | AGCCCTAGATG------------------**(99)**----------------------AAGCTTCTACC(27)CTACAGTGGAATCTGGTGAGATGGCAA | -139bp 7/16 |
|  | G1F1P02 | AGCCCTAGAG------------------**(99)**--------ATGCGCAGACGCTGAAGCTTCTACC(27)CTACAGTGGAATCTGGTGAGATGGCAA | -125bp 6/16 |
|  |  | AGCCCTAGAT--------------AAGAT(99)CACCCTGGAT-----------------------(15)------------------AGATGGCAA | -14/-56bp 1/16 |
|  |  | AGCCCTAGATG--------------------**(99)**--------------------------------**(15)**-----------------AGATGGCAA | -183bp 5/16 |
|  |  | AGCCCTAGATG------------------**(99)**----------------------AAGCTTCTACC(27)CTACAGTGGAATCTGGTGAGATGGCAA | -139bp 4/16 |
|  | G1F1P03 | AGCCCTAGAT-------------------**(99)**---------TGCGCAGACGCTGAAGCTTCTACC(27)CTACAGTGGAATCTGGTGAGATGGCAA | -127bp 11/16 |
|  |  | AGCCCTAGATGTCTGTCGATCCTCAAGAT(99)CACCCTGGA------------------------(15)------------------AGATGGCAA | -57bp 5/16 |
|  | G1F1P04 | AGCCCTAGAG------------------**(99)**--------ATGCGCAGACGCTGAAGCTTCTACC(27)CTACAGTGGAATCTGGTGAGATGGCAA | -125bp 8/15 |
|  |  | AGCCCTAGAT--------------AAGAT(99)CACCCTGGAT-----------------------(15)------------------AGATGGCAA | -14/-56bp 3/15 |
|  |  | AGCCCTAGATG------------------**(99)**----------------------AAGCTTCTACC(27)CTACAGTGGAATCTGGTGAGATGGCAA | -139bp 4/15 |
|  | G1F1P05 | AGCCCTAGAG------------------**(99)**--------ATGCGCAGACGCTGAAGCTTCTACC(27)CTACAGTGGAATCTGGTGAGATGGCAA | -125bp 4/16 |
|  |  | AGCCCTAGAT--------------AAGAT(99)CACCCTGGAT-----------------------(15)------------------AGATGGCAA | -14/-56bp 5/16 |
|  |  | AGCCCTAGATG--------------------**(99)**--------------------------------**(15)**-----------------AGATGGCAA | -183bp 2/16 |
|  |  | AGCCCTAGATG------------------**(99)**----------------------AAGCTTCTACC(27)CTACAGTGGAATCTGGTGAGATGGCAA | -139bp 5/16 |
|  | G1F1P06 | AGCCCTAGAT-------------------**(99)**---------TGCGCAGACGCTGAAGCTTCTACC(27)CTACAGTGGAATCTGGTGAGATGGCAA | -127bp 4/16 |
|  |  | AGCCCTAGATGTCTGTCGATCCTCAAGAT(99)CACCCTGGA------------------------(15)------------------AGATGGCAA | -57bp 3/16 |
|  |  | AGCCCTAGATG--------------------**(99)**--------------------------------**(15)**-----------------AGATGGCAA | -183bp 3/16 |
|  |  | AGCCCTAGATG------------------**(99)**----------------------AAGCTTCTACC(27)CTACAGTGGAATCTGGTGAGATGGCAA | -139bp 6/16 |
|  | G1F1P07 | AGCCCTAGAG------------------**(99)**--------ATGCGCAGACGCTGAAGCTTCTACC(27)CTACAGTGGAATCTGGTGAGATGGCAA | -125bp 14/16 |
|  |  | AGCCCTAGAT--------------AAGAT(99)CACCCTGGAT-----------------------(15)------------------AGATGGCAA | -14/-56bp 2/16 |
|  | G1F1P08 | AGCCCTAGAG------------------**(99)**--------ATGCGCAGACGCTGAAGCTTCTACC(27)CTACAGTGGAATCTGGTGAGATGGCAA | -125bp 5/14 |
|  |  | AGCCCTAGATG--------------------**(99)**--------------------------------**(15)**-----------------AGATGGCAA | -183bp 3/14 |
|  |  | AGCCCTAGATG------------------**(99)**----------------------AAGCTTCTACC(27)CTACAGTGGAATCTGGTGAGATGGCAA | -139bp 6/14 |
|  | G1F1P09 | AGCCCTAGAG------------------**(99)**--------ATGCGCAGACGCTGAAGCTTCTACC(27)CTACAGTGGAATCTGGTGAGATGGCAA | -125bp 11/15 |
|  |  | AGCCCTAGAT--------------AAGAT(99)CACCCTGGAT-----------------------(15)------------------AGATGGCAA | -14/-56bp 4/15 |
|  | G1F1P10 | AGCCCTAGAG------------------**(99)**--------ATGCGCAGACGCTGAAGCTTCTACC(27)CTACAGTGGAATCTGGTGAGATGGCAA | -125bp 11/15 |
|  |  | AGCCCTAGAT--------------AAGAT(99)CACCCTGGAT-----------------------(15)------------------AGATGGCAA | -14/-56bp 4/15 |
|  | G1F1P11 | AGCCCTAGAG------------------**(99)**--------ATGCGCAGACGCTGAAGCTTCTACC(27)CTACAGTGGAATCTGGTGAGATGGCAA | -125bp 10/16 |
|  |  | AGCCCTAGAT--------------AAGAT(99)CACCCTGGAT-----------------------(15)------------------AGATGGCAA | -14/-56bp 6/16 |

**Table S16 The summary of genotype in F1 generation of 10 GEC male pigs naturally mated with 8 GEC female pigs**

| No. | Male  parents | Female parents | ID | GGTA1 | CMAH | β4GalNT2 | hCD46 | hCD55 | hCD59 | hTBM | hCD39 | hEPCR | hCD47 |
| --- | --- | --- | --- | --- | --- | --- | --- | --- | --- | --- | --- | --- | --- |
| 1 | DN10TGC3F02P02 | DN8TGC12F01P17 | G1F1P01 | △52/△53  WT/△154 | △123/△1△3 | △183/△139/  △125/△14△56 | ○ | ○ | ○ | ○ | ○ | × | × |
| 2 |  |  | G1F1P02 | △52/+2△13  WT/△7△469 | △124/△102 | △183/△139/  △125/△14△56 | ○ | ○ | ○ | ○ | ○ | ○ | ○ |
| 3 |  |  | G1F1P03 | △52/+2△13  WT/△7△469 | △123/△1△3 | △125/△14△56 | ○ | ○ | ○ | ○ | ○ | ○ | ○ |
| 4 | DN10TGC3F02P02  or  DN10TGC3F03P07 | DN8TGC12F01P08 | G1F1P04 | △52/+2△13  WT/△7△469 | △124/△102 | △139/△183 | ○ | ○ | ○ | ○ | ○ | × | × |
| 5 |  |  | G1F1P05 | △52/+2△13  WT/△7△469 | △123/△1△3 | △183/△139/  △125/△14△56 | ○ | ○ | ○ | ○ | ○ | × | × |
| 6 |  |  | G1F1P06 | △52/+2△13  WT/△7△469 | △124/△102 | △183/△139/  △125/△14△56 | ○ | ○ | ○ | ○ | ○ | × | × |
| 7 |  |  | G1F1P07 | △52/+2△13  WT/△7△469 | △124/△102 | △125/△14△56 | ○ | ○ | ○ | ○ | ○ | × | × |
| 8 |  |  | G1F1P08 | △52/△53  WT/△154 | △123/△1△3 | △139/△125/  △14△56 | ○ | ○ | ○ | ○ | ○ | ○ | ○ |
| 9 |  |  | G1F1P09 | △52/△53  WT/△154 | △124/△102 | △125/△14△56 | ○ | ○ | ○ | ○ | ○ | ○ | ○ |
| 10 |  |  | G1F1P10 | △52/+2△13  WT/△7△469 | △124/△102 | △139/△183 | ○ | ○ | ○ | ○ | ○ | ○ | ○ |
| 11 |  |  | G1F1P11 | △52/+2△13  WT/△7△469 | △124/△102 | △125/△14△56 | ○ | ○ | ○ | ○ | ○ | × | × |
| 12 | DN10TGC3F01P04 | DN8TGC12F01P02 | G1F1P12 |  |  |  | ○ | ○ | ○ | ○ | ○ | ○ | ○ |
| 13 |  |  | G1F1P13 |  |  |  | ○ | ○ | ○ | ○ | ○ | × | × |
| 14 |  |  | G1F1P14 |  |  |  | ○ | ○ | ○ | ○ | ○ | ○ | ○ |
| 15 |  |  | G1F1P15 |  |  |  | ○ | ○ | ○ | ○ | ○ | × | × |
| 16 |  |  | G1F1P16 |  |  |  | ○ | ○ | ○ | ○ | ○ | ○ | ○ |
| 17 |  |  | G1F1P17 |  |  |  | ○ | ○ | ○ | ○ | ○ | × | × |
| 18 |  |  | G1F1P18 |  |  |  | ○ | ○ | ○ | ○ | ○ | × | × |
| 19 |  |  | G1F1P19 |  |  |  | ○ | ○ | ○ | ○ | ○ | × | × |
| 20 | DN10TGC3F02P02  or  DN10TGC3F04P12 | DN8TGC12F01P025 | G1F1P20 |  |  |  |  |  |  |  |  |  |  |
| 21 |  |  | G1F1P21 |  |  |  |  |  |  |  |  |  |  |
| 22 |  |  | G1F1P22 |  |  |  |  |  |  |  |  |  |  |
| 23 |  |  | G1F1P23 |  |  |  |  |  |  |  |  |  |  |
| 24 |  |  | G1F1P24 |  |  |  |  |  |  |  |  |  |  |

**Table S17 Information of recipient monkeys and donor pigs**

| NO. | ID | Gender | Age (year) | Weight (Kg) | Blood type | Transplantation | Donor Information |
| --- | --- | --- | --- | --- | --- | --- | --- |
| 1 | 335346# | Male | 16 | 15.65 | A | Kidney | 33dDN10TGC3F04P12 (32.4 Kg, 11 month, Male, Blood type O) |
| 2 | 823787# | Male | 17 | 16.45 | A | Kidney | 33dDN10TGC3F02P02 (34.1Kg, 10 month, Male, Blood type O) |
| 3 | 335362# | Male | 17 | 16.70 | AB | Liver |  |
| 4 | 3134985# | Male | 17 | 19.3 | A | Heart |  |
