## Supplementary figures and images for "Specific Pathogen Free Ten Gene-Edited Pig Donor for Xenotransplantation"

### Supplemental Figures.docx

Figure S1

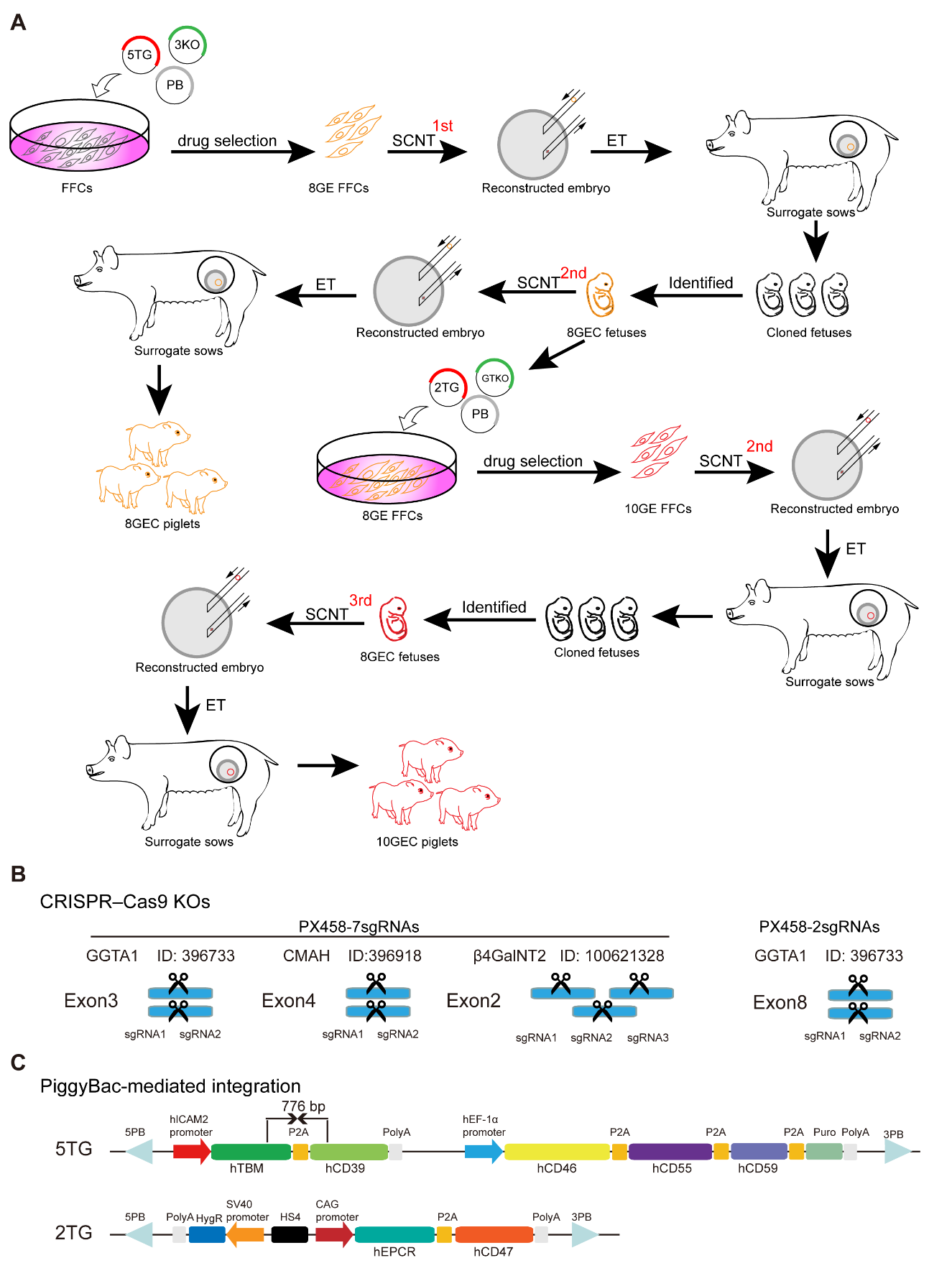

Figure S2

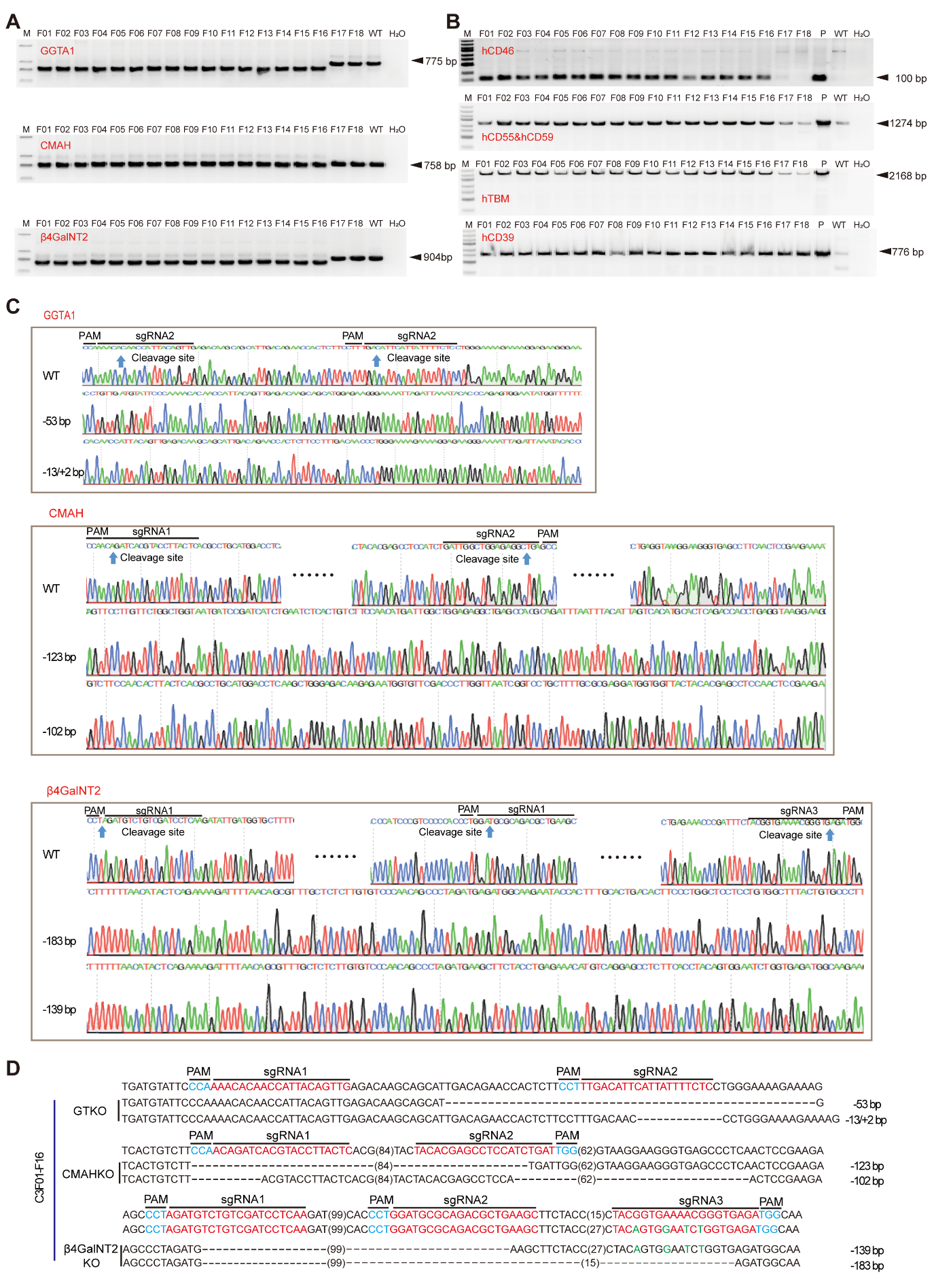

Figure S3

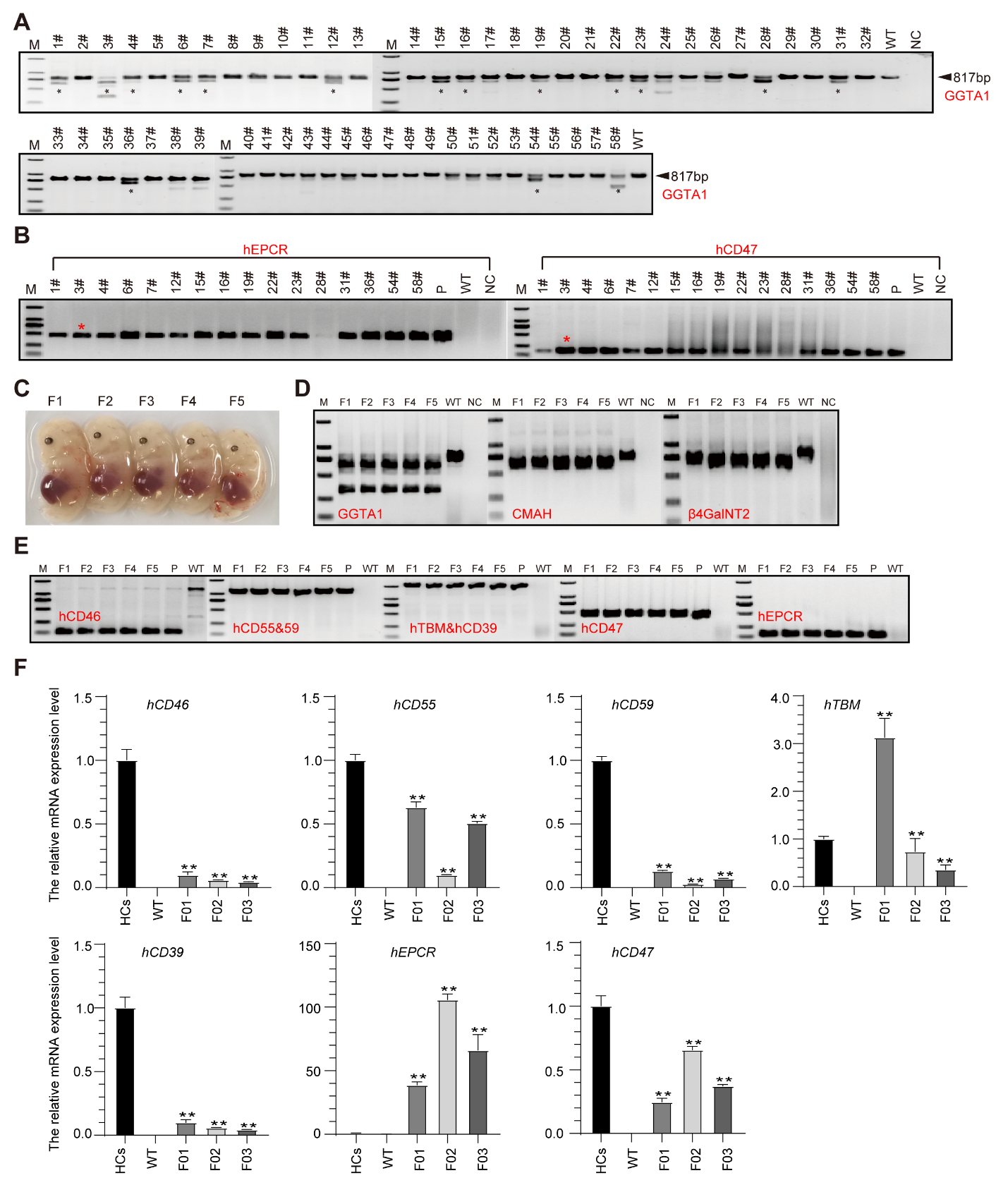

Figure S4

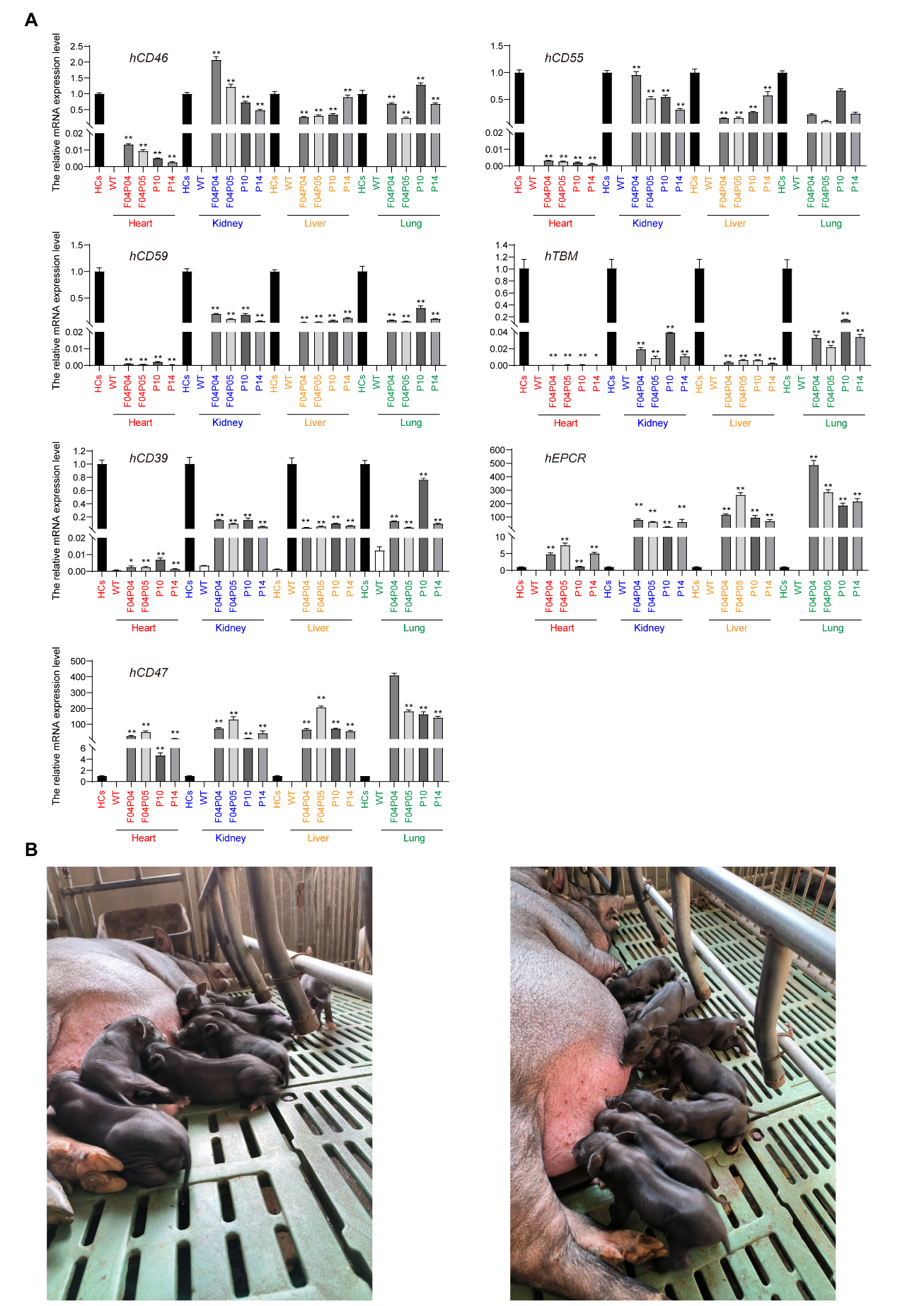

Figure S5

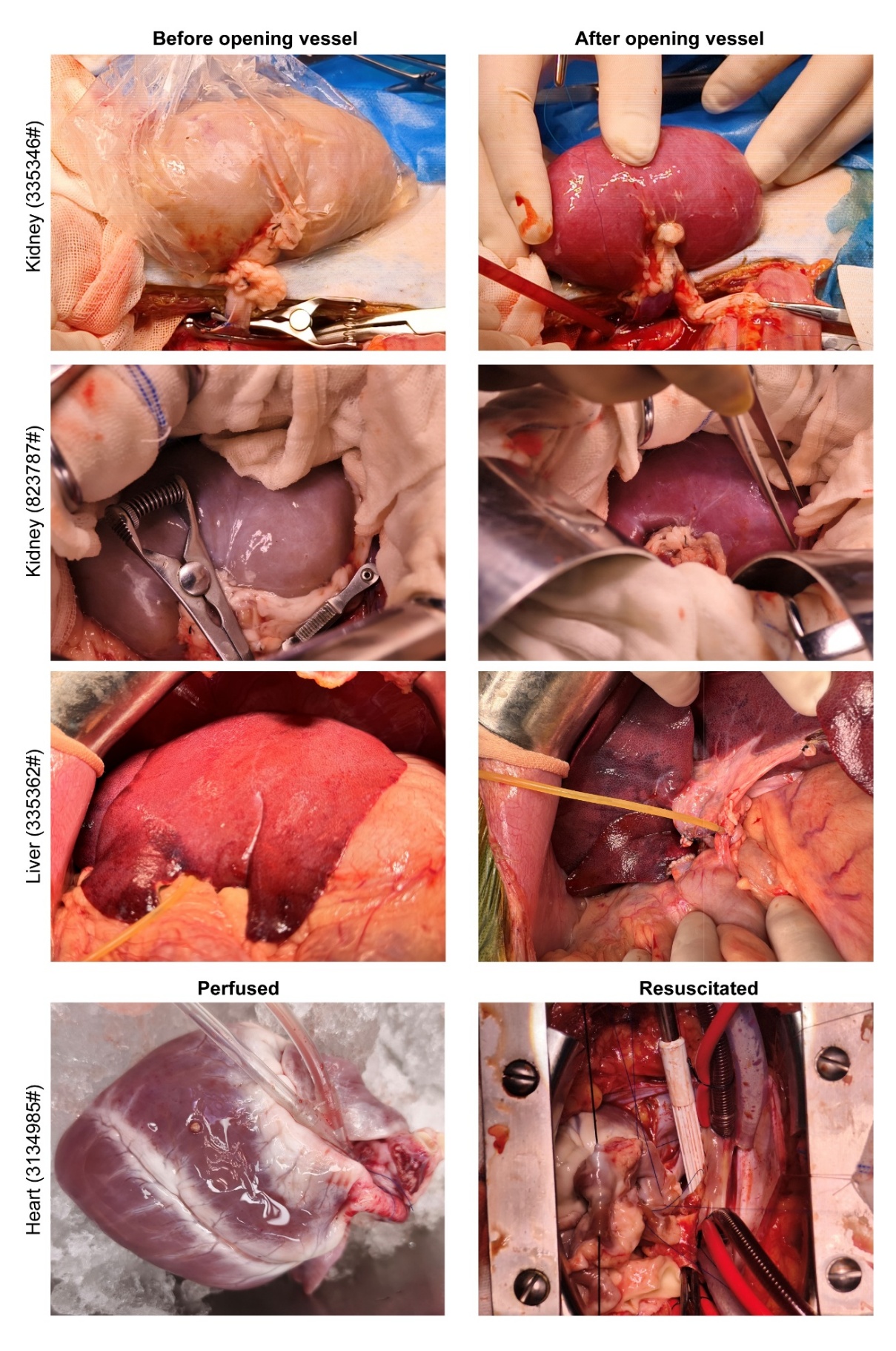

Figure S6

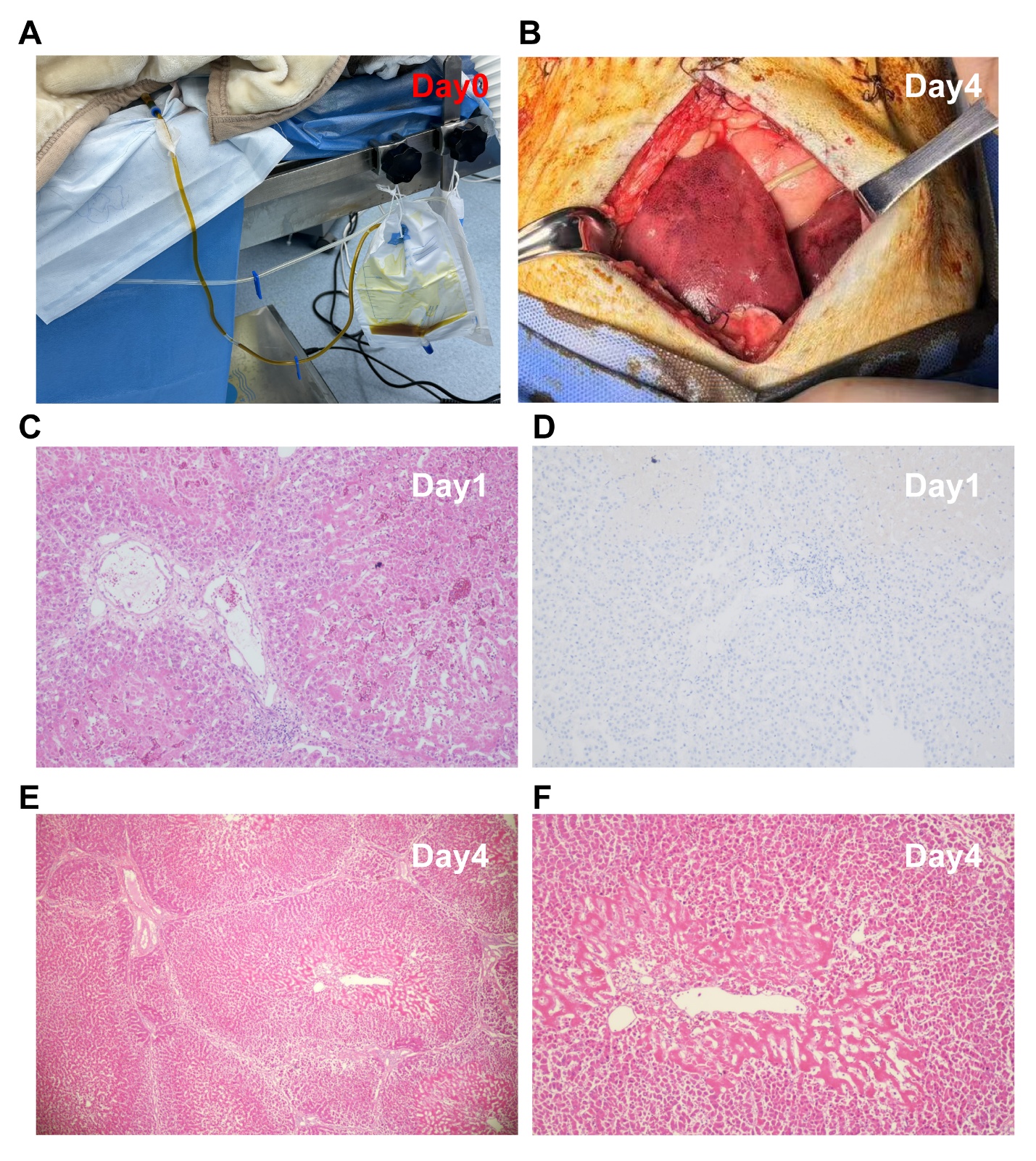

Figure S7

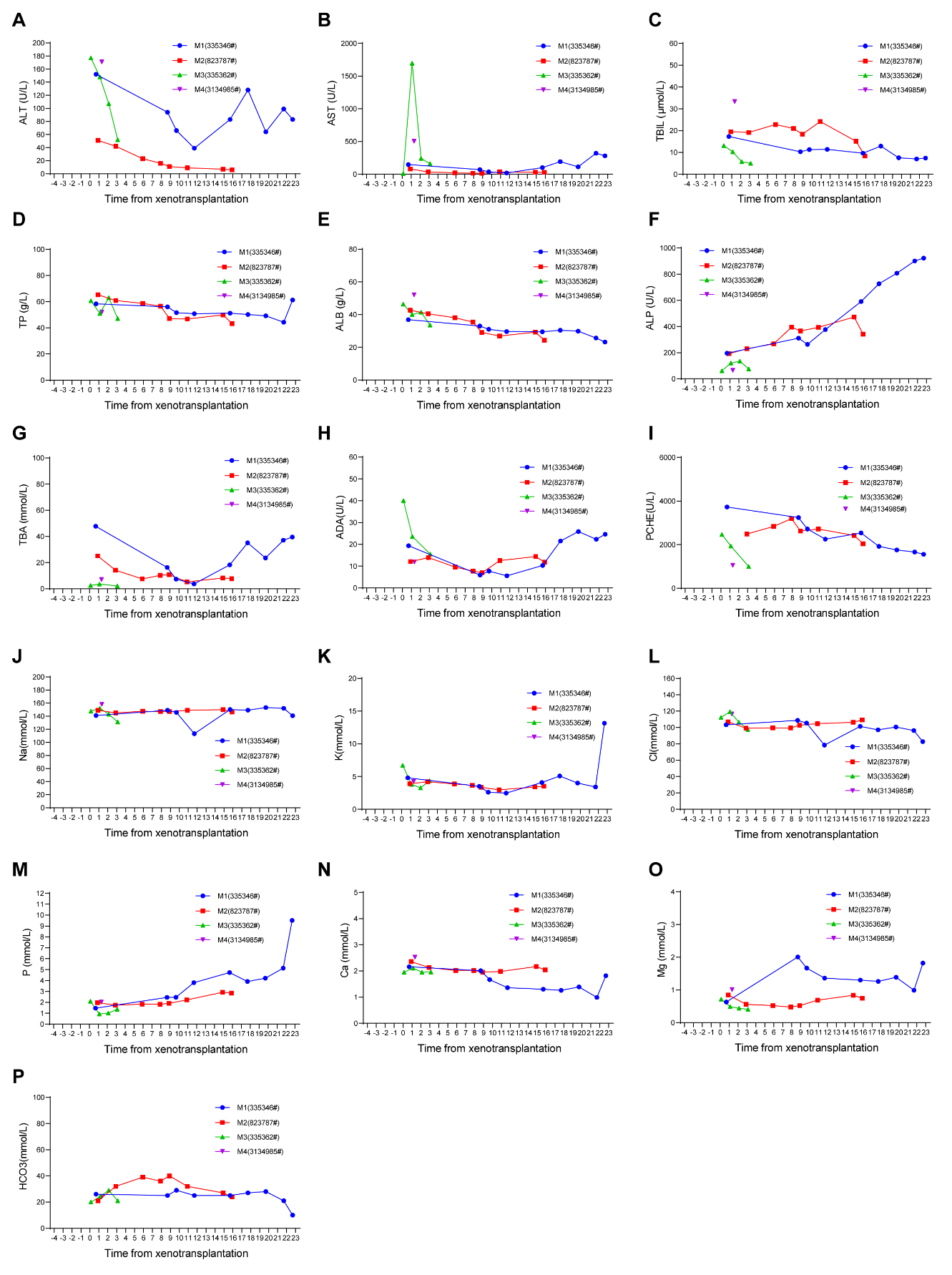
